# Iconic Manakins and Despicable Grackles: Comparing Bird-Related Cultural Ecosystem Services across Birdwatchers, Farmers, and Urbanites in Northwestern Costa Rica

**DOI:** 10.1101/548982

**Authors:** Alejandra Echeverri, Robin Naidoo, Daniel S. Karp, Kai M.A. Chan, Jiaying Zhao

## Abstract

Despite the great cultural and economic benefits associated with birdwatching and other bird-related cultural ecosystem services (CES), little is known about the bird-related CES perceived by people, and how they differ across stakeholder groups and species. The goal of this study was to explore CES across three stakeholder groups in northwestern Costa Rica. We conducted surveys (n=404 total) in which we presented farmers (n=140), urbanites (n=149), and birdwatchers (n=115) with pictures and songs of bird species and collected participants’ ratings on items designed to measure multiple CES. We found bird-related CES were perceived as six different constructs: identity, bequest, education, birdwatching, acoustic aesthetic, and disservices. The three stakeholder groups varied across these constructs and across species. Specifically, birdwatchers ranked species higher in terms of their education scores and lower in disservices scores compared to the other two groups. Positive correlations across CES, and negative correlations with disservices, suggest that the affect heuristic (by which generalized positive or negative feelings sway judgements of risks and benefits) might be informing bird-related CES. Our approach represents a novel method for assessing CES that can be adapted and modified for different taxa and multiple geographical contexts.

## 1 Introduction

Considering local communities’ knowledge and perceptions of biodiversity in conservation decisions is critical for the long-term protection of biodiversity (Berkes, 2004). Increasingly, the conservation and wildlife management communities are calling for more integrated approaches that incorporate peoples’ diverse values of nature, including how they perceive and value other species (Chan et al., 2016; Pascual et al., 2017). Such values vary across different groups of people, as they are shaped by cultural and socio-demographic contexts (Kenter, 2016; Peterson et al., 2010).

Cultural ecosystem services (CES) defined as “ecosystem’s contributions to the non-material benefits, such as capabilities and experiences, that arise from human-ecosystem relationships” (Chan et al., 2012b) are one of many theoretical frameworks used to characterize relationships between humans and ecosystems, and between humans and non-human animals (Echeverri et al., 2018). Though they are likely to motivate people’s connections with nature (Chan et al., 2012a; 2012b), little is known about how much and how particular species contribute to CES, and how this varies across stakeholder groups with different relationships to the non-human world (Gould and Lincoln, 2017; Milcu et al., 2013). Empirical work characterizing CES has focused mostly on landscapes and their associated services (e.g., place values) (Gould et al., 2014; Klain et al., 2014; Pascua et al., 2017). Fewer studies have analyzed the CES provided by and constructed with species or specific taxonomic groups (Milcu et al., 2013).

To date, research has focused on understanding the biophysical services that species provide to people (e.g., pest control, pollination) by identifying key species that act as ecosystem-service providers (Karp et al., 2013; Luck et al., 2012; Peisley et al., 2017; Whelan et al., 2008). Despite the great cultural and economic benefits associated with CES provided by species, such as wildlife viewing and aesthetic benefits, very little is known about the kinds of CES perceived by people. Aesthetic beauty is a commonly cited CES and is often related to biodiversity (Graves et al., 2017). However, the contribution of species to other CES categories such as identity, or importance for education benefits remain largely unexplored. Moreover, the psychological mechanisms that inform peoples’ perceptions towards biodiversity are a growing field of study, but remain largely unexplored (Clayton et al., 2013). For example, the affect heuristic (Finucane et al., 2000), first introduced in the psychology of risk perception, proposes that the inverse relationship between perceived risk and perceived benefits of specific hazards occur because people rely on affect (positive feelings) when judging them. In the fields of conservation and wildlife management, studies evaluating the perception of species that cause human-wildlife conflicts have evaluated the role of affect on informing such perceptions (Bruskotter and Wilson, 2014; Slagle et al., 2012), but little is known about how affect can inform perceived benefits from species, such as aesthetic beauty or other CES.

Birds are globally distributed, fill various ecological roles, and provide many ecosystem services to people (Sekercioglu, 2006; Whelan et al., 2008). For example, birds provide game meat for food (Fernandes-Ferreira et al., 2011), regulate pest populations (Karp et al., 2013), act as scavengers in agricultural landscapes (Peisley et al., 2017), and disperse seeds (Pigot et al., 2016). Culturally, they drive bird-watching tourism industries (e.g., Puhakka et al., 2011; U.S. Fish & Widlife Service, 2009). In the United States alone, estimates suggest that 46 million birdwatchers spend $32 billion each year, which contributes US$85 billion in economic output annually (Pullis La Rouche, 2006). Birds are also important characters in folk tales (Enríquez Rocha and Rangel Salazar, 2015), are common themes in the arts and have been portrayed in human languages, proverbs, and ceremonial activities for millennia (Watkins and Stockland, 2007). Thus, birds are great study organisms for characterizing CES associated with species.

Few studies have evaluated bird-related CES, and they have either examined single groups of people (e.g., Veríssimo et al., 2009), single services (e.g., Cox et al., 2018; Puhakka et al., 2011), or single species (e.g., Cortés-Avizanda et al., 2018). A study that compares CES across different groups of people is long overdue for informing current and future bird conservation actions. Moreover, similar studies are scarce in the growing body of literature attempting to characterize the plurality of perspectives on how people relate to the natural world (3-6). Thus, our main research question was: How do bird-related CES vary across bird species and stakeholder groups? Akin to biophysical services, we predicted that bird-related CES are perceived as separate constructs (e.g., birdwatching vs. bequest). We also predicted CES exhibit different rankings, such that different stakeholder groups perceive and value birds for different reasons.

## 2 Methods

### 2.1 Study region

This research took place in Northwestern Costa Rica (encompassing the Guanacaste and Puntarenas provinces). The region is rich in biodiversity, hosting ∼250 bird species and two Important Bird Areas (Devenish et al., 2009). Costa Rica is appropriate for studying bird-related CES because its biodiversity contributes significantly to its tourism industry (5-7% of GDP) (WTTC, 2014). Moreover, conservation discourses and economies have predominated in the country’s recent history, eliciting widespread awareness of biodiversity among Costa Ricans (Vivanco, 2006). People were surveyed in urban towns farmland and protected areas (e.g., Parque Nacional Barra Honda, Parque Nacional Palo Verde, Parque Nacional Diriá) across the region.

### 2.2 Data collection

In June-July 2016, we collected pilot data from 50 in-person surveys to identify key stakeholders in the region and to tailor the survey instrument to the local context. Moreover, a colleague conducted 20 semi-structured interviews with farmers for another study, but the interview protocol had two open-ended questions about the birds that they saw in their surroundings, and the species they found interesting, appealing, and problematic (Chapman, 2017). Results from the pilot data indicated that 8 species were the most prevalent for their positive and negative aspects. Based on the pilot data, we developed a survey to evaluate the perceived CES associated with birds by different stakeholder groups.

This study was conducted under the auspices of the University of British Columbia with Behavioral Research Ethics Board approval (#H16-00693). Surveys were administered in-person and online to 404 people during November and December 2017. Specifically, we surveyed farmers (n=140), urbanites (n=149), and international and local birdwatchers and birdwatching guides (n=115). Surveys were available in Spanish and English and were administered by the first author and six local field assistants. Sampling efforts were tailored for each stakeholder group. Specifically, urbanites were recruited by going to public spaces in urban towns across the peninsula (e.g., Liberia, Nicoya, Hojancha, Cañas, Sámara, Tambor). We visited central town parks, senior homes, universities, schools, and local fairs to recruit participants. The sample included people with a wide range of ages, different education levels, and 50% women, to get a representative sample of the population (Table 1). Farmers were recruited via partnerships with the Ministry of Agriculture and Cattle Ranching (MAG) in Nicoya and Hojancha and by attending cattle ranching fairs. We sampled both small-scale and large-scale farmers (e.g., 100m^2^ vs. 6.000 ha). We also sampled farmers who grew a variety of crops (e.g., sugar cane, rice, corn, oranges, mangoes) or who had cattle pastures. Even though women are less likely to be farmers in this region, we tried to sample as many women farmers as possible to minimize any bias in the data due to gender, however, only 22% of the farmers surveyed were women (Table 1). Lastly, to recruit birdwatchers we advertised the survey in Neotropical and European birdwatching forums and listservs (e.g., NEOORN-Neotropical Ornithology discussion list), in Facebook pages of Costa Rican birdwatching sites, and through the online bulletin of the Costa Rican ornithological association. We also attended two Christmas bird counts in Monteverde and Volcán Arenal (December 2017) and conducted in-person surveys during the meetings prior to the counts. Even though birdwatching is an activity that is mostly dominated (>75%) by males over the age of 45 in North America and Europe (Vas, 2017), we were able to cover a more demographically diverse sample (Table 1).

**Table 1.**
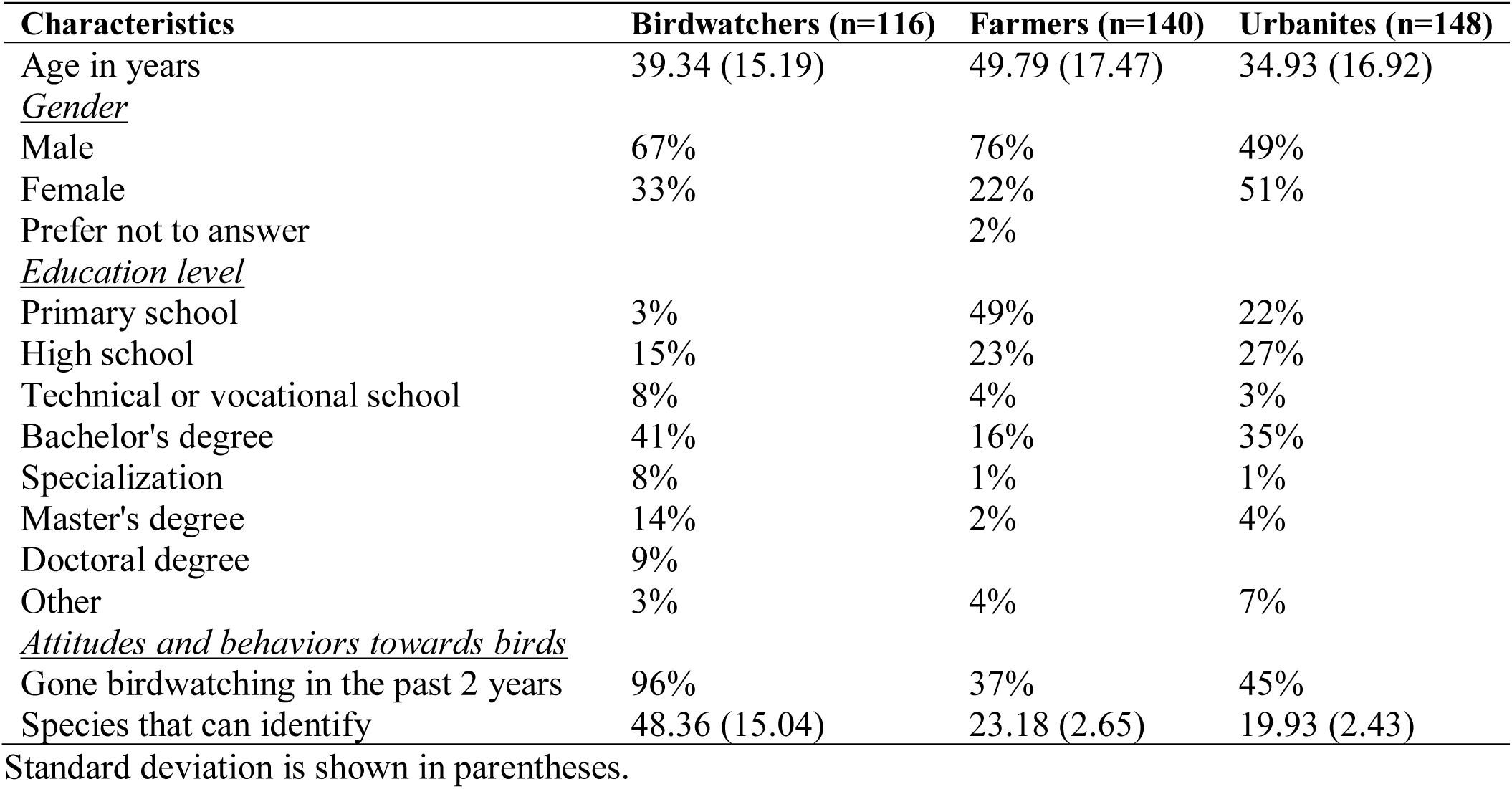
Characterization of participants according to demographic information and attitudes and behavior towards birds.

All survey responses for farmers and urbanites were recorded in person, but birdwatchers’ responses were recorded both online and in-person. Online responses (n=75) were mostly composed of international birdwatchers who had been birding in northwestern Costa Rica in the past but were not present at the time of sampling. All data were recorded on Qualtrics (a software for designing surveys).

### 2.3 Survey design

The survey had six sections. First, participants viewed a page with an introduction to the research, and the consent form. Then, they self-identified as either a birdwatcher, birdwatching guide, farmer, or urbanite; and answered questions tailored to each group. For instance, if they were birdwatchers or birdwatching guides, they were asked where they had been birdwatching. If they were farmers, they were asked what type of farm they owned or operated. Next, participants answered three open-ended questions about which birds they enjoyed watching or hearing, which birds they would like to protect for future generations, and which birds they perceived annoying or harmful. Then, participants ranked 12 or 13 species by answering 5-point Likert scale times (see below), and they answered three attitudinal questions about personal interest and self-reported behavior towards birds (e.g., birdwatching in the past, or reading books about birds). Lastly, they answered demographic questions (e.g., gender, education).

Each survey presented a set of 12 or 13 species that were presented in random order (Table S1). Surveys covered questions about 199 species detected in the region (Karp et al., 2018), but from those 199, 8 were focal species that appeared more frequently in species sets (Table S1). Those 8 focal species were selected as the most prevalently named species in the pilot data, perceived in both positive and negative terms (Table 2). For instance, pilot data had indicated that urbanites perceived the national bird (Clay-colored Thrush) and the Long-tailed Manakin as iconic in the region, while farmers indicated that the Great-tailed Grackle and the Orange-chinned Parakeet were eating their crops. Thus, we chose the 8 most salient species in the pilot data to become focal species for evaluating how different CES varied across species and stakeholder groups. We collected ratings on the other 191 species for a different research project, thus in this study we primarily report results for the 8 focal species and we focus on the comparisons across stakeholder groups.

**Table 2.**
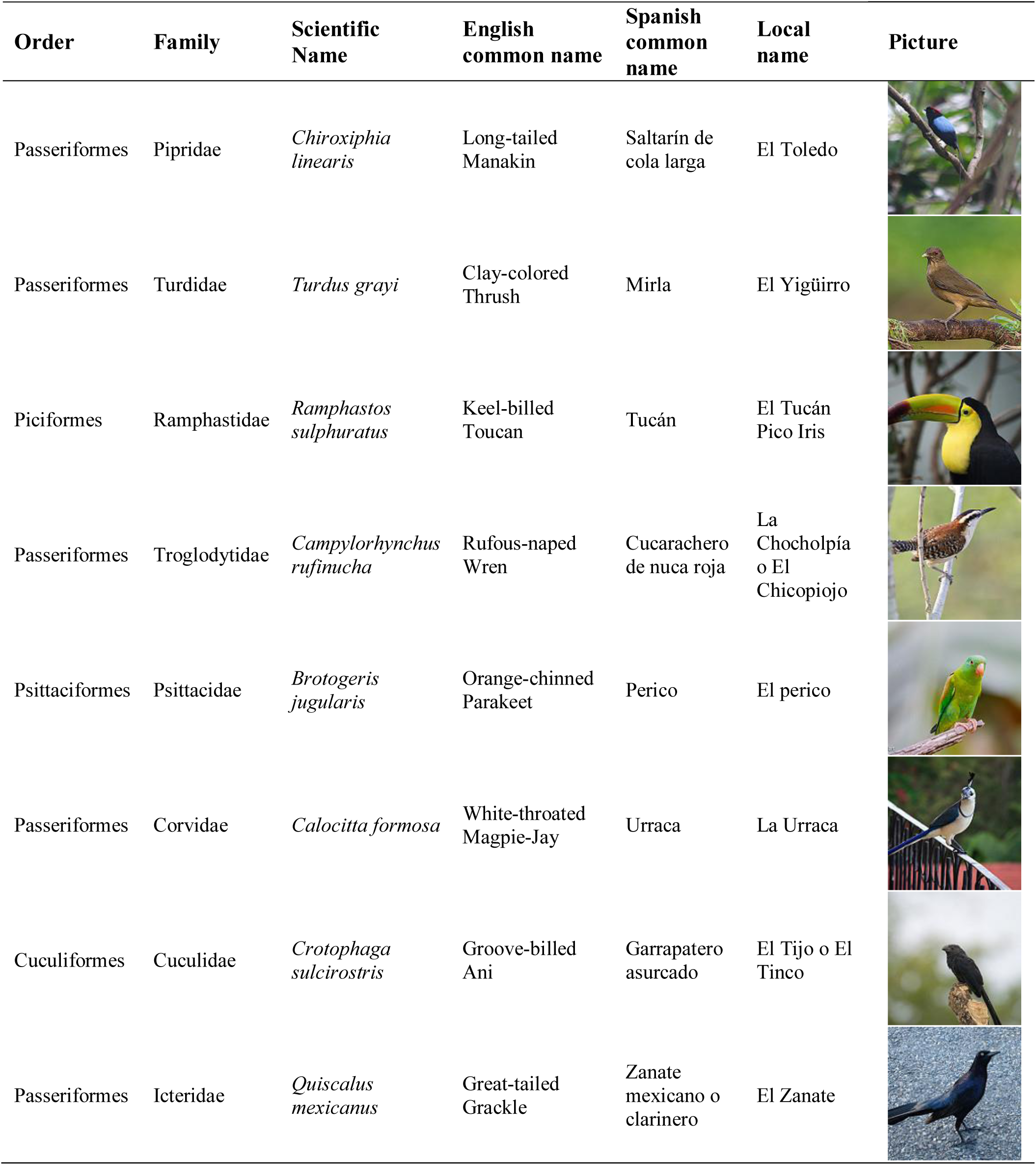
Focal species with their scientific, English, Spanish and local names.

Each species was represented with a visual illustration of a male individual (Garrigues and Dean, 2014) and an auditory clip of their song/call (xeno-canto.org; Table S2). For each species, participants were asked how much they liked each species, how frequently they saw the species in a given month, and whether they knew what species it was. If participants knew what the species was, then they were asked the name of the species and their subjective agreement on 12 different 5-point Likert scale items (Table 3) that ranged from strongly disagree (1) to strongly agree (5). Likert scale items were designed to measure different CES (beneficial and detrimental) that birds provide to people building on the categories from Gould et al. (2014), and Belaire et al. (2015). Items were tested and refined after collecting pilot data to ensure that the language in the items was simple enough so that a wide range of people could understand it.

**Table 3.** Factor analysis results indicating six different constructs that represent various cultural ecosystem services and disservices.

| Construct | Survey item | F1 | F2 | F3 | F4 | F5 | F6 |
| --- | --- | --- | --- | --- | --- | --- | --- |
| **Disservices**<br>(Cronbach's<br>alpha= 0.77) | This bird causes problems to other species that are important for me (reversed coded) | 0.56 | 0.00 | 0.03 | 0.04 | 0.00 | 0.07 |
|  | This bird causes problems to the crops or the farms by for example eating the crop (reversed coded) | 0.79 | 0.01 | 0.00 | -0.10 | -0.01 | 0.03 |
|  | I find this bird annoying because it is too noisy | 0.58 | 0.00 | 0.02 | 0.12 | 0.16 | -0.07 |
|  | I dislike this bird because their droppings make a mess or they build nests in inconvenient places | 0.54 | 0.05 | 0.03 | 0.21 | -0.07 | -0.04 |
| **Education**<br>(Cronbach's<br>alpha=0.83) | I like learning about or studying this bird, where it lives and what it does | 0.00 | 0.80 | 0.02 | 0.03 | -0.02 | 0.01 |
|  | I like teaching others about this bird and its habitat | -0.01 | 0.88 | -0.01 | -0.02 | 0.02 | 0.00 |
| **Bequest**<br>(Cronbach's<br>alpha=0.84) | This bird should be protected for future generations | -0.01 | 0.00 | 1.01 | -0.02 | 0.01 | -0.01 |
|  | It would be sad if this bird would no longer exist | 0.06 | 0.06 | 0.57 | 0.18 | -0.01 | 0.07 |
| **Birdwatching**<br>(Cronbach's<br>alpha=0.84) | This bird is beautiful and I enjoy watching it | -0.02 | -0.01 | 0.02 | 0.84 | 0.03 | 0.02 |
|  | I am excited to find this bird | 0.08 | 0.12 | 0.03 | 0.66 | 0.04 | 0.06 |
| **Acoustic**<br>**aesthetic**<br>(NA) | This bird has a beautiful song | 0.00 | 0.01 | 0.01 | 0.01 | 0.99 | 0.01 |
| **Identity**<br>(Cronbach's<br>alpha=0.58) | This bird is like my neighbor and makes me feel at home | -0.02 | 0.00 | -0.08 | 0.01 | 0.07 | 0.63 |
|  | This bird helps make this place what it is | 0.03 | 0.04 | 0.12 | 0.06 | -0.04 | 0.59 |
| NA | This bird is good for the farms, by for example eating rodent pests | 0.19 | 0.02 | 0.03 | -0.12 | 0.10 | 0.01 |
| SS loadings |  | 1.79 | 1.58 | 1.55 | 1.54 | 1.08 | 0.93 |
| Inertia explained (%) |  | 13% | 11% | 11% | 11% | 8% | 7% |
| Inertia accumulated (%) |  | 13% | 24% | 35% | 46% | 54% | 61% |

If participants did not know what the species was, then they were only presented with 2 Likert-scale items measuring bequest also with a 5-point Likert scale (Table 3, bequest). See supplementary material for a copy of the survey.

### 2.4 Data analysis

We first examined whether there was a difference between the patterns in responses of the Likert scale items between local and international birdwatchers, between birdwatchers and birdwatching guides, and between online and in-person responses. We found no significant differences between any of these comparisons after conducting one-way ANOVAs (p’s>0.05), suggesting that birdwatchers could be considered one stakeholder group in subsequent analyses. Therefore, we pooled data from all these groups and called them “birdwatchers”.

We performed an exploratory factor analysis with all complete observations on the 14 different Likert-scale items designed to measure various CES categories. To examine the number of factors, we used the “fa.parallel” function instituted in the “psych” package (Revelle, 2017) in the statistical software R (R Development Core Team, 2008). Then, we conducted a factor analysis with “oblimin” rotation and maximum likelihood instituted in the “GPA rotation” package also in R. Factor analysis operates on the notion that measurable and observable variables can be reduced to fewer latent variables that share a common variance and are unobservable, which is known as reducing dimensionality (Bartholomew et al., 2011).

We used a factor loading threshold of 0.5 to assign Likert scale items to different factors and calculated Cronbach’s alpha for internal consistency. The factor analysis yielded 6 different factors, representing different bird-related service categories. Only one of the 14 items did not load to any factor, so we excluded that item for posterior analyses (Table 3; NA). With the results from the factor loadings, we then calculated the mean scores for the items in each factor to create 6 constructs and used them as dependent variables in two analyses (Table 3). For the first analysis, we pooled data from all species (including non-focal species) and created linear mixed-effects models to predict the effect of stakeholder group on the 6 dependent variables. For such analyses, we treated both ‘species’ and ‘participant’ as random effects. For the second analysis, we only used data from the 8 focal species and regressed each dependent variable against species, stakeholder group and the interaction between the two via using linear mixed-effects models. We also used a random intercept of ‘participant’. For all models, we did posterior checks to test for normality and heteroscedasticity assumptions. We then conducted type II ANOVAs to test for the significance of the main effects and used Tukey HSD as post-hoc tests.

In the second analysis, the acoustic aesthetic and bequest constructs did not conform to normality/heteroscedasticity assumptions. Therefore, we used a multinomial regression to evaluate the effects of species and stakeholder groups on the acoustic aesthetic construct. For the multinomial regression, we treated each of the 5 points in the Likert scale as a potential outcome (i.e., response variable), and calculated the probabilities of each stakeholder group ranking each species on any of the five points. Moreover, bequest scores were analyzed as a binary variable because the data were dominated by responses in the “agree” and “strongly agree” categories (i.e., categories 4 and 5). We collapsed scores lower or equal than 3.5 and called them “disagree”, and scores higher than 3.5 and called them “agree”. The 3.5 threshold represented the most conservative estimate for “agreement”, as it assumed that for one of two Likert scale items measuring bequest, a person would have to score at least one item as 4 (somewhat agree) and the other as 3 (neutral), or one as 5 (strongly agree) and one as 2 (somewhat disagree). With the newly constructed binary variable, we conducted a logistic regression predicting bequest with species, stakeholder group, and the interaction between the two. All analyses were done in R (R Development Core Team, 2008). See supplementary material for details from all statistical models and model fits.

We coded the qualitative data from the open-ended questions to identify the most common species mentioned by birdwatchers, farmers, and urbanites when prompted with birdwatching, bequest, and disservices questions. We counted the frequency of mentions for species (e.g., Long-tailed Manakin) or bird groups (e.g., Toucans) by each stakeholder group, and used frequencies as a metric of how salient that bird species/group was for each stakeholder group. The complete data is presented in Tables S9-S11.

## 3 Results

### 3.1 Comparing cultural ecosystem services across species and stakeholder groups

Results from the factor analysis indicated the presence of six factors among the Likert scale items (Table 3). Those factors were interpreted as different constructs that represented various CES and disservices. Below we present how each of them varied by species and stakeholder groups.

### 3.2 Disservices

Results from the open-ended questions showed that all three stakeholder groups found the Great-tailed Grackle to be most harmful and most annoying (Table 4). Birdwatchers and urbanites found vultures second most harmful/annoying, unlike farmers who found parakeets most harmful/annoying (Table 4).

**Table 4.**
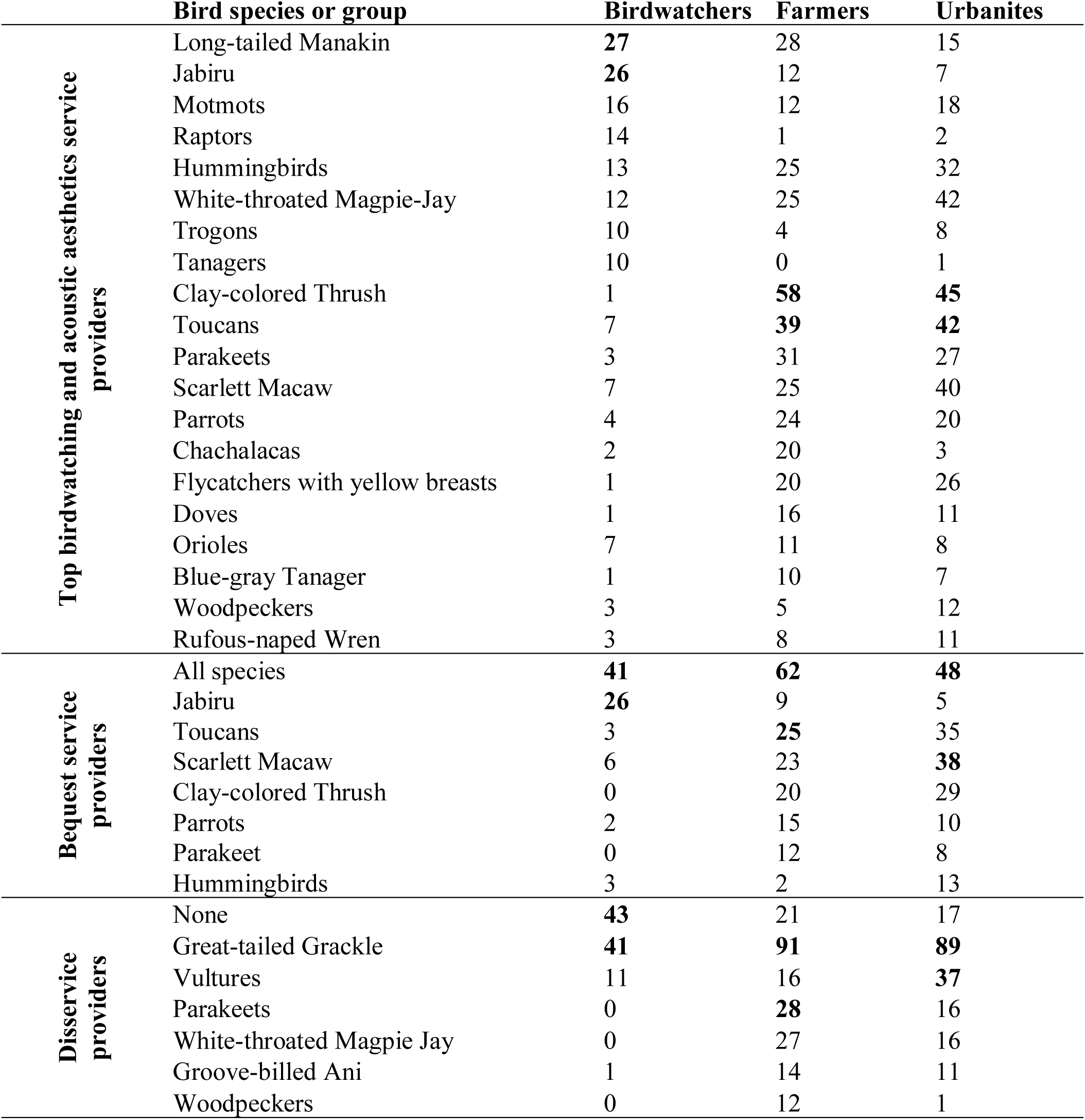
Most frequently mentioned bird species or groups by each stakeholder group in the open-ended questions.

|  | **Bird species or group** | **Birdwatchers** | **Farmers** | **Urbanites** |
| --- | --- | --- | --- | --- |
| **Top birdwatching and acoustic aesthetics service providers** | Long-tailed Manakin | **27** | 28 | 15 |
|  | Jabiru | **26** | 12 | 7 |
|  | Motmots | 16 | 12 | 18 |
|  | Raptors | 14 | 1 | 2 |
|  | Hummingbirds | 13 | 25 | 32 |
|  | White-throated Magpie-Jay | 12 | 25 | 42 |
|  | Trogons | 10 | 4 | 8 |
|  | Tanagers | 10 | 0 | 1 |
|  | Clay-colored Thrush | 1 | **58** | **45** |
|  | Toucans | 7 | **39** | **42** |
|  | Parakeets | 3 | 31 | 27 |
|  | Scarlett Macaw | 7 | 25 | 40 |
|  | Parrots | 4 | 24 | 20 |
|  | Chachalacas | 2 | 20 | 3 |
|  | Flycatchers with yellow breasts | 1 | 20 | 26 |
|  | Doves | 1 | 16 | 11 |
|  | Orioles | 7 | 11 | 8 |
|  | Blue-gray Tanager | 1 | 10 | 7 |
|  | Woodpeckers | 3 | 5 | 12 |
|  | Rufous-naped Wren | 3 | 8 | 11 |
| **Bequest service providers** | All species | **41** | **62** | **48** |
|  | Jabiru | **26** | 9 | 5 |
|  | Toucans | 3 | **25** | 35 |
|  | Scarlett Macaw | 6 | 23 | **38** |
|  | Clay-colored Thrush | 0 | 20 | 29 |
|  | Parrots | 2 | 15 | 10 |
|  | Parakeet | 0 | 12 | 8 |
|  | Hummingbirds | 3 | 2 | 13 |
| **Diservice providers** | None | **43** | 21 | 17 |
|  | Great-tailed Grackle | **41** | **91** | **89** |
|  | Vultures | 11 | 16 | **37** |
|  | Parakeets | 0 | **28** | 16 |
|  | White-throated Magpie Jay | 0 | 27 | 16 |
|  | Groove-billed Ani | 1 | 14 | 11 |
|  | Woodpeckers | 0 | 12 | 1 |

Regarding the quantitative data, we found that birdwatchers perceived fewer disservices than farmers and urbanites across all species (Fig. 1; df=2, *χ* ^2^ =50.56, p<0.0001). We also found that disservices varied across species (df=7, *χ* ^2^ =1211.83, p<0. 0001), stakeholder groups (df=2, *χ* ^2^ =56.96, p<0.0001), and the interaction between the two (df=14, *χ* ^2^ =112.06, p<0.0001). Tukey HSD post-hoc tests showed that birdwatchers perceived less disservices from four species compared to farmers and urbanites (Rufous-naped Wren, Orange-chinned Parakeet, White-throated Magpie-Jay, and Great-tailed Grackle, p<0.05; Fig. 2). Results from the pairwise comparisons for all post-hoc tests are presented in the supplementary material.

**Figure 1.**
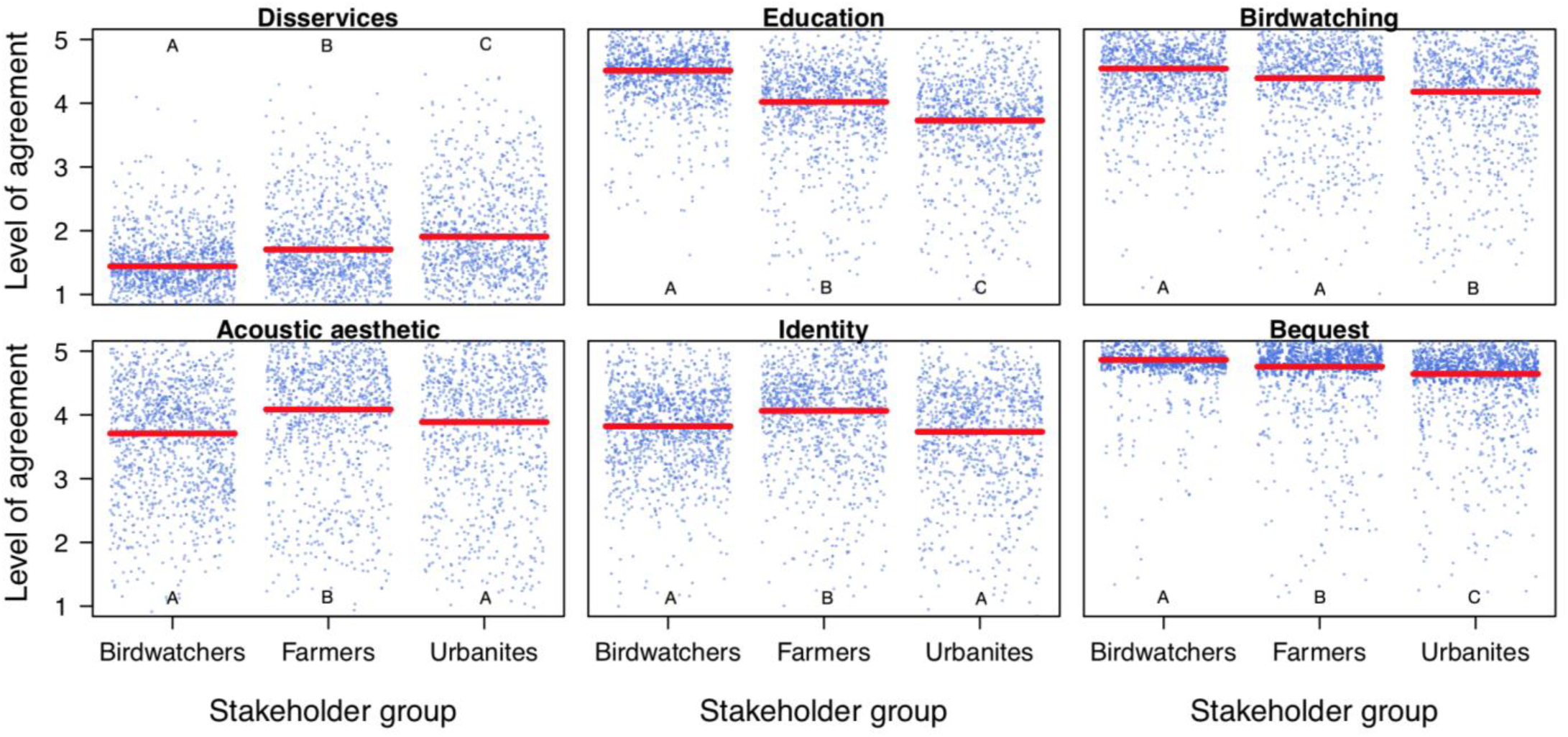
Relationship between stakeholder groups and their perceived cultural ecosystem service categories across species. Each dot represents an individuals’ ranking for a species for each service category. Red lines represent the modelled estimate in the linear mixed-effects models for each stakeholder group in each service category. Distributions marked with the same letters are not statistically different from one another, while those not sharing any letters in common are significantly different distributions according to post-hoc tests (p< 0.05).

**Figure 2.**
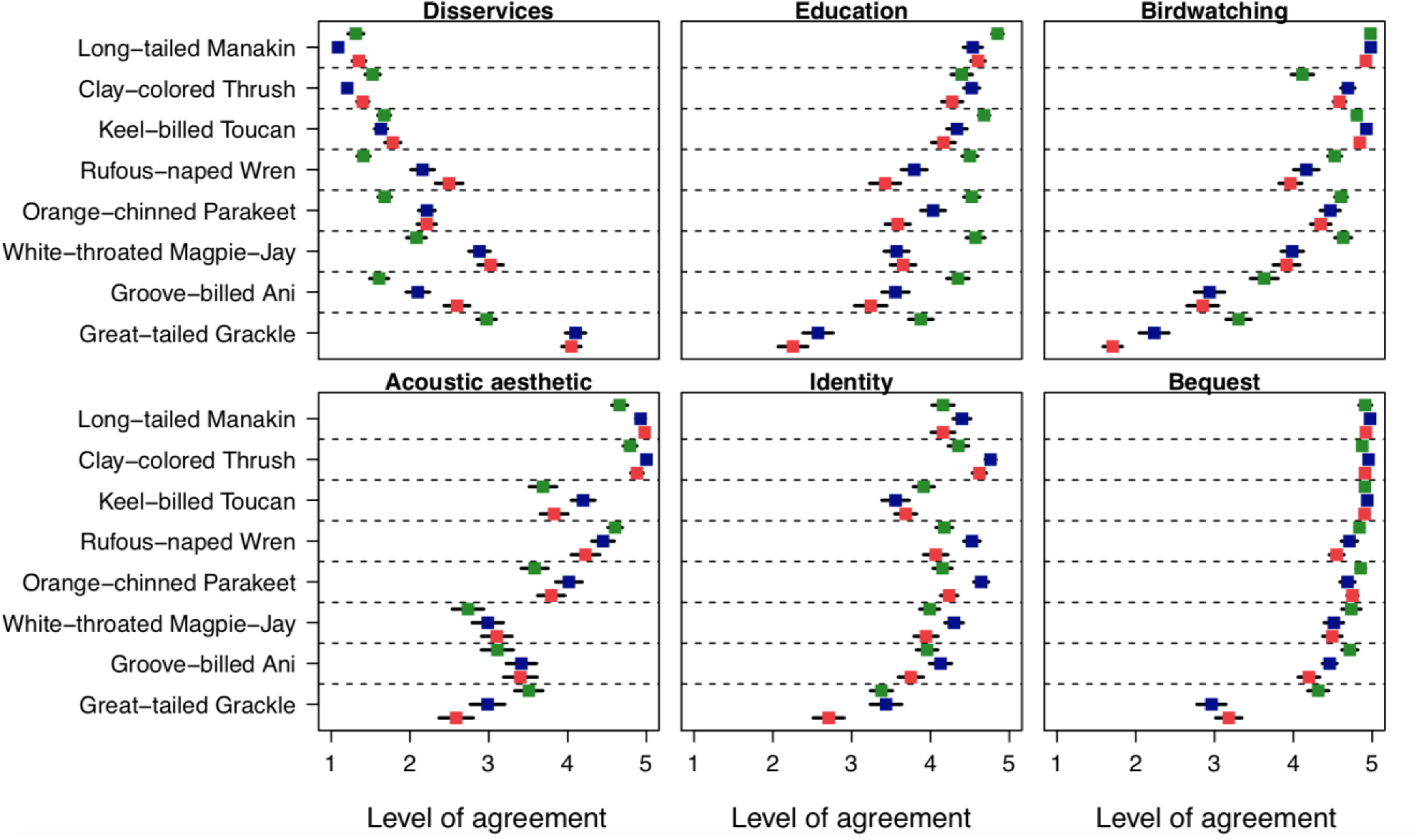
Scores for six ecosystem service categories across eight focal species and stakeholder groups. Each square represents the mean value for the level of agreement for birdwatchers (green), farmers (blue), and urbanites (red). The level of agreement was assessed on a five-point Likert scale: 1-strongly disagree, 2-somewhat disagree, 3-neither agree nor disagree, 4-somewhat agree, 5-strongly agree. We present the mean scores for each species with 95% confidence intervals (black lines).

### 3.3 Education

Birdwatchers allocated higher education scores than farmers and urbanites across all species (Fig. 1) (df=2, *χ* ^2^ =53.80, p<0.0001). We also found that education varied across species (df=7, *χ* ^2^ =451.02, p< 0.0001), stakeholder groups (df=2, *χ* =62.14, p<0.0001), and the interaction between the two (df=14, *χ* ^2^ =71.64, p<0.0001). Post-hoc tests showed no differences across stakeholder groups (p>0.05) for the Long-tailed Manakin and the Clay-colored Thrush, as all groups displayed strong education in these species (Fig. 2). Birdwatchers scored 5 other species significantly higher than farmers and urbanites. We did not find statistically significant differences between farmers and urbanites for any of the 8 focal species (p>0.05) (Fig. 2).

### 3.4 Birdwatching and acoustic aesthetics

We found that birdwatchers mentioned 156 different species or groups of birds (e.g., raptors) that they enjoyed watching or hearing. Of those, 77 were only mentioned once. The most mentioned species among birdwatchers were the Long-tailed Manakin (n=26) and the Jabiru (n=26; Table 4). Conversely, farmers mentioned 60 species or groups, 15 of which were only mentioned once. The most mentioned were the Clay-colored Thrush (n=58) and toucans (n=39). Finally, urbanites mentioned 70 different species or groups that they enjoyed watching or hearing, 22 of which were only mentioned once. The most-mentioned species were the Clay-colored Thrush (n=45), toucans (n=42), and the White-throated Magpie-Jay (n=42) (Table 4).

Birdwatchers and farmers ranked species higher on birdwatching scores than urbanites across all species (df=2, *χ* ^2^ =28.24, p<0.0001), while farmers perceived higher acoustic aesthetics than the other two groups across species (df=2, *χ* ^2^ =18.14, p<0.001) (Fig. 1). We found that birdwatching varied across species (df=7, *χ* ^2^ =1214.81, p<0.0001), stakeholder groups (df=2, *χ* ^2^ =29.26, p<0.0001), and the interaction between the two (df=14, *χ* ^2^ =111.02, p<0.0001). Post-hoc tests indicated no significant differences between groups for 3 species (Orange-chinned Parakeet, Long-tailed Manakin, and Keel-billed Toucan; p>0.05), as all groups found them visually appealing (Fig. 2). Birdwatchers perceived 2 species as having higher birdwatching scores compared to farmers and urbanites (Great-tailed Grackle, and White-throated Magpie-Jay; p<0.001), and one species lower than farmers and urbanites (Clay-colored Thrush, p<0.001).

The multinomial regression showed that the acoustic aesthetics varied by species (df=28, LR *χ* ^2^ =469.74, p<0.0001), stakeholder groups (df=8, *χ* ^2^ =47.88, p<0.0001), and the interaction between the two (df=56, LR *χ* ^2^ =82.20, p<0.05). We did not find significant differences across species and stakeholder groups for 6 species (Groove-billed Ani, Rufous-naped Wren, Orange-chinned Parakeet, White-throated Magpie-Jay, Keel-billed Toucan and Clay-colored Thrush; p>0.05). Birdwatchers rated the song of the Long-tailed Manakin significantly lower than the other two groups (p<0.001).

### 3.5 Identity

Farmers ranked species’ identity scores higher than birdwatchers and urbanites across species (df=2, *χ* ^2^ =16.83, p<0.001) (Fig. 1). We also found that identity scores varied across species (df=7, *χ* ^2^ =262.20, p<0.0001), stakeholder groups (df=2, *χ* ^2^ =15.32, p<0.001), and the interaction between the two (df=14, *χ* ^2^ =32.39, p<0.005). A trend observed from Fig. 2 is that farmers’ identity scores across species were higher for all but one species (Keel-billed Toucan). For example, farmers scored Orange-chinned Parakeet and Clay-colored Thrush significantly higher in terms of their identity scores compared to both birdwatchers and urbanites (p<0.005). Additionally, urbanites scored the Great-tailed Grackle lower than the other two groups (p<0.05). Post-hoc tests found no significant differences across stakeholder groups for four species (Groove-billed Ani, Long-tailed Manakin, White-throated Magpie-Jay, and Keel-billed Toucan; p>0.05).

### 3.6 Bequest

When participants were prompted with bequest questions (i.e., which species would you like to protect for future generations?), the most mentioned answer across all three groups was “All species” (birdwatchers=41, farmers=62, urbanites=48). Birdwatchers named 59 species or groups when prompted with bequest questions, farmers named 41, and urbanites named 56. The most mentioned species or groups are presented in Table 4.

All stakeholder groups assigned high scores on bequest across species (Fig. 1): the means for all 3 groups were higher than 4.5, indicating a high level of agreement on bequest items. Nonetheless, we found that birdwatchers had higher bequest scores compared to the other two groups (df=2, *χ* ^2^ =19.38, p<0.0001). After analyzing bequest as a binary variable (>=3.5 was coded as agree, and <=3.5 was coded as disagree), we found that they varied with species (df=7, LR *χ* ^2^ =260.27, p<0.0001), stakeholder groups (df=2, LR *χ* ^2^ =32.28, p<0.0001), but not the interaction between the two (df=14, LR *χ* ^2^ =17.99, p=0.207). Post-hoc tests showed that the Great-tailed Grackle was the only species for which bequest scores differed by stakeholder groups, as birdwatchers ranked its bequest scores higher than farmers and urbanites (p<0.00001; see supplementary material). The other seven species did not show statistically significant differences across their bequest scores when comparing different stakeholder groups (p>0.05).

### 3.7 Correlations across cultural ecosystem services

Pairwise correlations across cultural ecosystem services and disservices categories indicated that for all three stakeholder groups, disservices were negatively correlated with the other five CES categories. Meanwhile identity, education, bequest, birdwatching and acoustic aesthetics were positively correlated with one another. Farmers and urbanites exhibit strong correlations (Pearson’s r>0.5) between birdwatching and education, acoustic aesthetics, and bequest (Fig. 3). Additionally, they exhibit strong negative correlations (Pearson’s r<-0.5) between disservices and birdwatching and bequest (Fig. 3). These correlations suggest that for farmers and urbanites, when birds are perceived as beautiful, they are also perceived as having a beautiful song, worthy of conserving, and worthy of studying (or vice versa). However, when looking at birdwatchers, the correlations trends are similar, but the strength of the relationship is weaker (Fig. 3).

**Figure 3.**
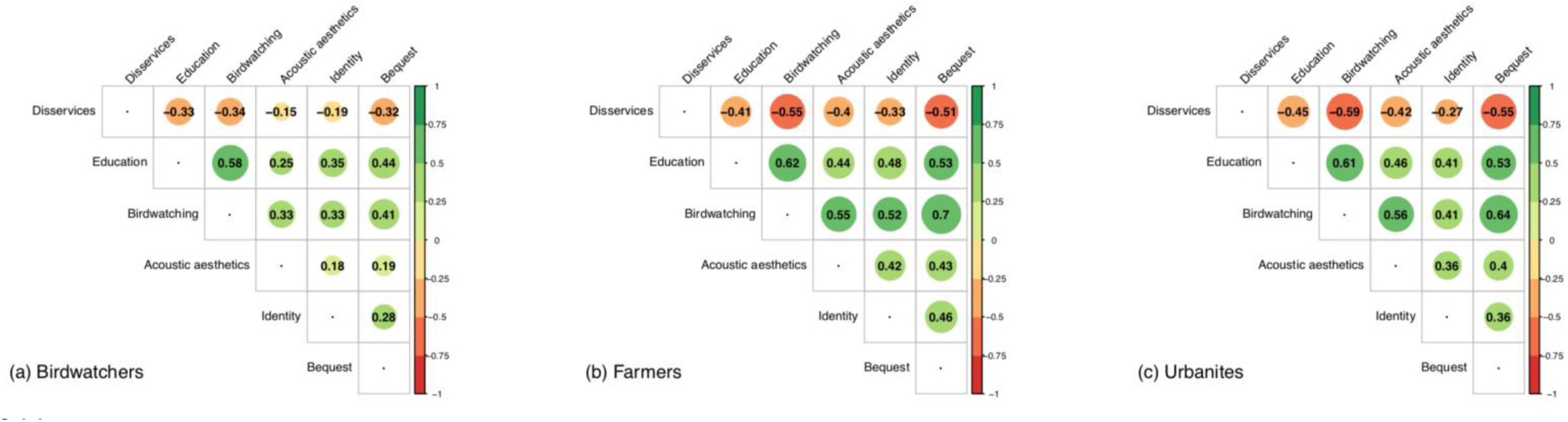
Pairwise correlations across cultural ecosystem service categories for each stakeholder group. The size of the circles represents the magnitude of the pairwise correlation, where bigger circles mean higher correlation. The numbers refer to the correlation coefficients (Pearson’s r) warmer colors (yellow, orange, red) indicate negative correlations, and cooler colors (greens) indicate positive correlations.

## 4 Discussion

Birdwatchers, farmers, and urbanites all expressed high interest in birds and identified various benefits that they derive from and construct with birds. Overall trends indicated that birdwatchers perceived higher scores regarding education, birdwatching, and bequest across species compared to the other two groups. In contrast, farmers scored species highest on identity and acoustic services compared to the other groups (Fig. 1). These results suggest that CES perceived by people vary strongly across stakeholder groups. Moreover, we identified strong trade-offs between the species that were perceived as CES providers vs. those that were disservice providers. For instance, when looking at the qualitative data, it was evident that the birds that people enjoyed watching were not the only ones that they wanted to protect for future generations (Table 4). However, regarding the quantitative data, we found positive correlations among identity, bequest, birdwatching, acoustic aesthetics and education, suggesting that birds that got high ratings on one of those CES categories also received high ratings on the other categories. In fact, the strongest correlation was found between birdwatching and education for all three stakeholder groups (Fig. 3), which is consistent with the taxonomic bias present in biological data gathering where biologists are more motivated to study charismatic species (Troudet et al., 2017).

While most birds were perceived positively, people also identified negative aspects of some birds. Our results echo those of Cox et al. (2018) who showed that in urban areas of southern England, people perceived 2.5 times as many bird species to be positive for people’s well-being relative to those whose behaviors caused conflict. We found similar results in our study where the birds that people found harmful to the crops or infrastructure were also the ones they perceived as annoying or noisy. The species perceived as most harmful/annoying was by far the Great-tailed Grackle. It was viewed as a pest in agricultural landscapes by farmers, as a nuisance in urban areas by urbanites, and as an invasive species in ecosystems by some birdwatchers. These results are consistent with previous studies on endangered species, where disliked species were also perceived as threatening (Echeverri et al., 2017).

We found negative correlations between disservices and the other five CES categories (Fig. 3). This finding can be explained by the affect heuristic (Finucane et al., 2000), which suggests that people rely on affect when judging risks (i.e., disservices) and benefits (i.e., beneficial CES, such as birdwatching) associated with species. In our study, general likeability (i.e., positive affect) of the birds was positively correlated with education, birdwatching, acoustic aesthetics, identity, and bequest, but was negatively correlated with disservices (Table S12). Importantly, the strength of these correlations was weaker for birdwatchers than for farmers or urbanites (Fig. 3). We believe this finding can be explained by the fact that birdwatchers are more knowledgeable (i.e., experts) about bird species and therefore are able to engage in rational reasoning more than relying on the affect heuristic to inform their perceptions towards species, unlike non-birdwatchers who inform their perceptions through affective measures (Markowitz et al., 2013).

bird-related CES differed strongly across species and stakeholder groups. For the most part, farmers and urbanites had very similar perceptions of birds regarding all species and all services. Only the Groove-billed Ani was perceived as more harmful for urbanites than for farmers (Fig. 2), mostly because farmers identified it as a species that gleans parasitic insects off of cattle, while urbanites often confused it with the Great-tailed Grackle (viewed as a pest locally). Given that farmers interact with birds through their livelihood activities more often than urbanites, we expected them to have differing perceptions. However, we believe the similarity between the perceptions of farmers and urbanites regarding birds speaks directly to the Costa Rican context and the deep association between Costa Ricans and the natural environment (Vivanco, 2006).

Birds are an important component of Costa Rican lifestyles. For instance, over 150,000 parrots are kept as pets in Costa Rican households (Drews, 2001). Also, birds are frequently mentioned in local Indigenous stories and folk tales. For example, in local stories, barn owls (*Tyto alba*) are believed to announce deaths in neighborhoods (Enríquez Rocha and Rangel Salazar, 2015). Similarly, the Clay-colored Thrush (*Turdus grayi)* and the Laughing Falcon (*Herpetotheres cachinnans*) are believed to announce when the rains are coming (Sault, 2010). Additionally, birds are present in Costa Rican folklore; they are often depicted in murals, local art, and are an important part of how Costa Ricans advertise their country to foreigners. The strong connection between people and birds in this context might also explain the species egalitarianism elicited across people when prompted with bequest questions (Schmidtz, 1998).

Regionally, our results have implications for environmental education and conservation campaigns. For instance, by far the most iconic species for all three stakeholder groups was the Long-tailed Manakin (*Chiroxiphia linearis*). It received the highest rankings on birdwatching, acoustic aesthetics, bequest and education, and the lowest rankings on disservices (Fig. 2). Current conservation and environmental awareness campaigns highlight the Jabiru (*Jabiru mycteria*) as a focal species, and it is the logo of the protected areas network and the Guanacaste Conservation Area (SINAC-ACG). However, we found that Jabirus were not very prominent in peoples’ minds when doing our pilot surveys and interviews, in fact not everyone could recognize this species. Also, Jabirus were mentioned more often by birdwatchers than by farmers and urbanites (Table 4). Our results suggest that the Long-tailed Manakin would be a more appropriate species to highlight in education and conservation campaigns as it is liked by all stakeholder groups, has cultural significance, a high degree of familiarity, and widely appreciated charisma (Bowen-Jones and Entwistle, 2002). Birdwatchers, farmers, and urbanites all commented positively on this species’ courtship dances, color and song. Given that this species is associated with wet forests, and species associated with wetter, more forested sites are more vulnerable to land-use and climate change (Karp et al., 2018), this species might be a good candidate for raising awareness on ecological issues, such as the expected droughts for the region (Hund et al., 2018).

The fact that CES are perceived differently by different people has been of central concern in decision-making for conservation planning and wildlife management (Daniel et al., 2012; Teel and Manfredo, 2010). Many conservation initiatives begin with stakeholders discussing realistic scenarios for the conservation of species’ habitats and how such scenarios might affect their own livelihoods or practices (Rosa et al., 2017). This process identifies salient ecosystem services and often involves people recognizing the cultural aspects of their relationships with species that are not yet captured by most protocols. For example, regarding fish conservation, projects might balance the competing cultural ecosystem services of recreational, commercial, and charter fisheries, as well as diving companies that prioritize tourism benefits — all including stakeholders who relate to fish in different ways (Beaudreau et al., 2011). Our method is well-suited to identify the diverse CES that people derive from and construct with species and could be easily adapted to existing conservation planning initiatives for birds and for other taxa. It provides a more systematic and nuanced assessment of the CES perceived by different stakeholder groups. For example, we have showed that species can rank differently on disservices and other CES categories (Fig. 2). Such rankings enable a systematic evaluation of trade-offs associated with different species.

Our study built on the method developed by Belaire et al. (2015) by adding survey items that elicit other CES (e.g., bequest, education), and by allowing people to discuss individual species instead of commenting on birds as a whole group. An important contribution of our paper is therefore the methodological advancement for measuring and eliciting CES associated with species. With our method, we identified species that act as CES and disservice providers. Identifying ecosystem-service providers is important for managing ecosystem services (Kremen, 2005; Luck et al., 2009) and for making conservation decisions that require assessing competing trade-offs between species or taxa (Karp et al., 2015). For example, stakeholders in the recreation sector could conduct conservation planning in areas that target populations of the species deemed most preferred for birdwatching purposes, and wildlife managers could identify people’s perceptions towards species that are culled and garner public support for their actions.

Interestingly, most people (not only birdwatchers) were able to identify birds by their songs (more so than the images), perhaps because they perceived them to be important elements of their daily soundscapes. Our method may thus be useful for researchers and practitioners attempting to capture people’s relationships with non-human animals, as it can be adapted to any geographical context and many species. We received positive feedback from participants regarding the enjoyment they derived from completing an interactive survey that showed both pictures and songs of birds. Many participants stated that this was a novel tool, which was different from traditional paper-based surveys. However, we also received negative feedback about the length of the survey from participants who knew most species (and so had to score all or most species). Thus, we recommend future applications of the method to reduce the number of species presented so that the survey takes at most half an hour to complete.

We acknowledge that our methods are rooted in Western science, which can be deemed as inappropriate for analyzing Traditional Ecological Knowledge (Bartlett et al., 2012). Therefore, future research directions may include an ethno-ornithological analysis of individual species in our dataset and their associated myths, folklore, proverbs, and knowledge. A more qualitative study could help understand more in depth the relationships between people and birds that are not captured by our methods. Additionally, future research could focus on using our methods in other geographical contexts to test for the generalizability of peoples’ perceptions of species across their distribution range.

## 5 Conclusions

Although birds provide many important biophysical ecosystem services (Whelan et al., 2008), perhaps one of their most important roles is their contribution to the non-material benefits that people derive and construct with them (Echeverri et al., 2018). Birds of northwestern Costa Rica carried many stories associated with death, peace, as indicators of climate, as characters in religious references and proverbs. Responses from different stakeholder groups revealed that for birdwatchers, urbanites, and farmers, the surrounding landscape is not quiet or still: it is alive, full of songs they want to hear when they wake up, and full of birds they think beautify the landscapes and the skies. Relationships between people and birds in this place might be leveraged to motivate pro-environmental actions and support ongoing conservation initiatives, such as reforestation efforts.

Here we presented a method that allowed us to analyze people’s relationships with birds by characterizing various cultural ecosystem services and by assessing multiple species. Importantly, our methodological contribution is timely considering the prominent pushes to incorporate diverse values in conservation and environmental management (Chan et al., 2016; Pascual et al., 2017).

## 6 Acknowledgments

We thank study participants for taking their time to answer the survey. We thank A. Valverde, J. Valverde, A. Zúñiga, A.Rojas, E. Obando, and W. Lázaro for their assistance with data collection. Special thanks to B. Freeman, S. David, R. May, J. Zook, J.D. Vargas, R. Guindon, T. Karwinkel, M. Tacke and Asociación de Ornitólogos de Costa Rica for helping us recruit birdwatchers for this study. We thank D. Schluter, J.A. Belaire, L.O. Frishkoff, and M. Chapman for their insightful comments. Special thanks to Christian Echeverri for his invaluable help with analysis. Research was funded by a National Geographic Young Explorers Grant (#C335-16) awarded to AE, a U.S. Forest Service grant (16-IJ-11242309-057) awarded to JZ, and a SSHRC Insight Grant (Human-nature relationships and values from meanings to mitigation and practice Insight Grant) awarded to KMAC. AE was supported by a Graduate Global Leadership Doctoral Fellowship and a Killam Doctoral Fellowship awarded through UBC.

