## Supplementary material for "Iconic Manakins and Despicable Grackles: Comparing Bird-Related Cultural Ecosystem Services across Birdwatchers, Farmers, and Urbanites in Northwestern Costa Rica"

### 1 EXPERIMENTAL DESIGN: SPECIES PRESENTED

**Table S1. Species presented in each one of the 22 versions of the survey. The eight focal species reported in this study are bolded.**

| Survey version | English name | Order | Family | Scientific name |
|---|---|---|---|---|
| 1 | Barred Antshrike | Passeriformes | Thamnophilidae | *Thamnophilus doliatus* |
| 1 | Brown-crested Flycatcher | Passeriformes | Tyrannidae | *Myiarchus tyrannulus* |
| **1** | **Clay-colored Thrush** | **Passeriformes** | **Turdidae** | ***Turdus grayi*** |
| **1** | **Groove-billed Ani** | **Cuculiformes** | **Cuculidae** | ***Crotophaga sulcirostris*** |
| 1 | Laughing Falcon | Falconiformes | Falconidae | *Herpetotheres cachinnans* |
| 1 | Lesser Swallow-tailed Swift | Apodiformes | Apodidae | *Panyptila cayennensis* |
| **1** | **Long-tailed Manakin** | **Passeriformes** | **Pipridae** | ***Chiroxiphia linearis*** |
| 1 | Olivaceous Woodcreeper | Passeriformes | Furnariidae | *Sittasomus griseicapillus* |
| 1 | Rufous-browed Peppershrike | Passeriformes | Vireonidae | *Cyclarhis gujanensis* |
| 1 | Scarlet Macaw | Psittaciformes | Psittacidae | *Ara macao* |
| 1 | American Swallow-tailed Kite | Accipitriformes | Accipitridae | *Elanoides forficatus* |
| 1 | Wood Stork | Ciconiiformes | Ciconiidae | *Mycteria americana* |
| **1** | **White-throated Magpie-Jay** | **Passeriformes** | **Corvidae** | ***Calocitta formosa*** |
| 2 | Blue-black Grassquit | Passeriformes | Thraupidae | *Volatinia jacarina* |
| 2 | Blue Ground Dove | Columbiformes | Columbidae | *Claravis pretiosa* |
| **2** | **Clay-colored Thrush** | **Passeriformes** | **Turdidae** | ***Turdus grayi*** |
| 2 | Common Black-Hawk | Accipitriformes | Accipitridae | *Buteogallus anthracinus* |
| 2 | Common Tody-Flycatcher | Passeriformes | Tyrannidae | *Todirostrum cinereum* |
| **2** | **Groove-billed Ani** | **Cuculiformes** | **Cuculidae** | ***Crotophaga sulcirostris*** |
| **2** | **Long-tailed Manakin** | **Passeriformes** | **Pipridae** | ***Chiroxiphia linearis*** |
| 2 | Ruddy Woodcreeper | Passeriformes | Furnariidae | *Dendrocincla homochroa* |
| 2 | Vaux's Swift | Apodiformes | Apodidae | *Chaetura vauxi* |
| 2 | White-fronted Parrot | Psittaciformes | Psittacidae | *Amazona albifrons* |
| **2** | **White-throated Magpie-Jay** | **Passeriformes** | **Corvidae** | ***Calocitta formosa*** |
| 2 | Yellow-green Vireo | Passeriformes | Vireonidae | *Vireo flavoviridis* |
| 2 | Yellow-headed Caracara | Accipitriformes | Accipitridae | *Milvago chimachima* |
| 3 | Blue-gray Tanager | Passeriformes | Thraupidae | *Thraupis episcopus* |
| 3 | Bare-throated Tiger-Heron | Pelecaniformes | Ardeidae | *Tigrisoma mexicanum* |
| **3** | **Clay-colored Thrush** | **Passeriformes** | **Turdidae** | ***Turdus grayi*** |
| 3 | Common Ground-Dove | Columbiformes | Columbidae | *Columbina passerina* |
| 3 | Crane Hawk | Accipitriformes | Accipitridae | *Geranospiza caerulescens* |
| 3 | Dusky-capped Flycatcher | Passeriformes | Tyrannidae | *Myiarchus tuberculifer* |
| **3** | **Groove-billed Ani** | **Cuculiformes** | **Cuculidae** | ***Crotophaga sulcirostris*** |

| Survey version | English name | Order | Family | Scientific name |
|---|---|---|---|---|
| 3 | Great Curassow | Craciformes | Cracidae | *Crax rubra* |
| **3** | **Long-tailed Manakin** | **Passeriformes** | **Pipridae** | ***Chiroxiphia linearis*** |
| 3 | Streak-headed Woodcreeper | Passeriformes | Furnariidae | *Lepidocolaptes souleyetii* |
| 3 | White-collared Swift | Apodiformes | Apodidae | *Streptoprocne zonaris* |
| **3** | **White-throated Magpie-Jay** | **Passeriformes** | **Corvidae** | ***Calocitta formosa*** |
| 3 | Yellow-naped Parrot | Psittaciformes | Psittacidae | *Amazona auropalliata* |
| 4 | Barn Swallow | Passeriformes | Hirundinidae | *Hirundo rustica* |
| 4 | Black-crowned Night-Heron | Pelecaniformes | Ardeidae | *Nycticorax nycticorax* |
| 4 | Buff-throated Saltator | Passeriformes | Thraupidae | *Saltator maximus* |
| 4 | Canivet's Emerald | Apodiformes | Trochilidae | *Chlorostilbon canivetii* |
| **4** | **Clay-colored Thrush** | **Passeriformes** | **Turdidae** | ***Turdus grayi*** |
| 4 | Double-toothed Kite | Accipitriformes | Accipitridae | *Harpagus bidentatus* |
| 4 | Ferruginous Pygmy-Owl | Strigiformes | Strigidae | *Glaucidium brasilianum* |
| **4** | **Groove-billed Ani** | **Cuculiformes** | **Cuculidae** | ***Crotophaga sulcirostris*** |
| 4 | Gray-headed Chachalaca | Galliformes | Cracidae | *Ortalis cinereiceps* |
| 4 | Gray-headed Dove | Columbiformes | Columbidae | *Leptotila plumbeiceps* |
| 4 | Great Kiskadee | Passeriformes | Tyrannidae | *Pitangus sulphuratus* |
| **4** | **Long-tailed Manakin** | **Passeriformes** | **Pipridae** | ***Chiroxiphia linearis*** |
| **4** | **White-throated Magpie-Jay** | **Passeriformes** | **Corvidae** | ***Calocitta formosa*** |
| 5 | Boat-billed Heron | Pelecaniformes | Ardeidae | *Cochlearius cochlearius* |
| **5** | **Clay-colored Thrush** | **Passeriformes** | **Turdidae** | ***Turdus grayi*** |
| 5 | Cinnamon Hummingbird | Apodiformes | Trochilidae | *Amazilia rutila* |
| **5** | **Groove-billed Ani** | **Cuculiformes** | **Cuculidae** | ***Crotophaga sulcirostris*** |
| 5 | Great Black Hawk | Accipitriformes | Accipitridae | *Buteogallus urubitinga* |
| 5 | Gray-breasted Martin | Passeriformes | Hirundinidae | *Progne chalybea* |
| 5 | Greenish Elaenia | Passeriformes | Tyrannidae | *Myiopagis viridicata* |
| 5 | Gray-headed Tanager | Passeriformes | Thraupidae | *Eucometis penicillata* |
| 5 | Inca Dove | Columbiformes | Columbidae | *Columbina inca* |
| **5** | **Long-tailed Manakin** | **Passeriformes** | **Pipridae** | ***Chiroxiphia linearis*** |
| 5 | Mottled Owl | Strigiformes | Strigidae | *Ciccaba virgata* |
| 5 | Plain Chachalaca | Galliformes | Cracidae | *Ortalis vetula* |
| **5** | **White-throated Magpie-Jay** | **Passeriformes** | **Corvidae** | ***Calocitta formosa*** |
| 6 | Cattle Egret | Pelecaniformes | Ardeidae | *Bubulcus ibis* |
| **6** | **Clay-colored Thrush** | **Passeriformes** | **Turdidae** | ***Turdus grayi*** |
| 6 | Crested Bobwhite | Galliformes | Odontophoridae | *Colinus cristatus* |
| **6** | **Groove-billed Ani** | **Cuculiformes** | **Cuculidae** | ***Crotophaga sulcirostris*** |
| 6 | Gray Hawk | Accipitriformes | Accipitridae | *Buteo plagiatus* |

| Survey version | English name | Order | Family | Scientific name |
|---|---|---|---|---|
| 6 | Grayish Saltator | Passeriformes | Thraupidae | *Saltator coerulescens* |
| **6** | **Long-tailed Manakin** | **Passeriformes** | **Pipridae** | ***Chiroxiphia linearis*** |
| 6 | Green-breasted Mango | Apodiformes | Trochilidae | *Anthracothorax prevostii* |
| 6 | Mangrove Swallow | Passeriformes | Hirundinidae | *Tachycineta albilinea* |
| 6 | Northern Beardless-Tyrannulet | Passeriformes | Tyrannidae | *Camptostoma imberbe* |
| 6 | Pacific Screech-Owl | Strigiformes | Strigidae | *Megascops cooperi* |
| 6 | Plain-breasted Ground-Dove | Columbiformes | Columbidae | *Columbina minuta* |
| **6** | **White-throated Magpie-Jay** | **Passeriformes** | **Corvidae** | ***Calocitta formosa*** |
| 7 | Bronzed Cowbird | Passeriformes | Icteridae | *Molothrus aeneus* |
| **7** | **Clay-colored Thrush** | **Passeriformes** | **Turdidae** | ***Turdus grayi*** |
| **7** | **Groove-billed Ani** | **Cuculiformes** | **Cuculidae** | ***Crotophaga sulcirostris*** |
| 7 | Gray-headed Kite | Columbiformes | Columbidae | *Leptodon cayanensis* |
| 7 | Great Egret | Pelecaniformes | Ardeidae | *Ardea alba* |
| 7 | Least Grebe | Podicipediformes | Podicipedidae | *Tachybaptus dominicus* |
| **7** | **Long-tailed Manakin** | **Passeriformes** | **Pipridae** | ***Chiroxiphia linearis*** |
| 7 | Northern Bentbill | Passeriformes | Tyrannidae | *Oncostoma cinereigulare* |
| 7 | Palm Tanager | Passeriformes | Thraupidae | *Thraupis palmarum* |
| 7 | Plain-capped Starthroat | Apodiformes | Trochilidae | *Heliomaster constantii* |
| 7 | Red-billed Pigeon | Columbiformes | Columbidae | *Patagioenas flavirostris* |
| 7 | Spectacled Owl | Strigiformes | Strigidae | *Pulsatrix perspicillata* |
| **7** | **White-throated Magpie-Jay** | **Passeriformes** | **Corvidae** | ***Calocitta formosa*** |
| 8 | Barn Owl | Strigiformes | Tytonidae | *Tyto alba* |
| 8 | Green Heron | Pelecaniformes | Ardeidae | *Butorides virescens* |
| **8** | **Clay-colored Thrush** | **Passeriformes** | **Turdidae** | ***Turdus grayi*** |
| 8 | Eastern Meadowlark | Passeriformes | Icteridae | *Sturnella magna* |
| **8** | **Groove-billed Ani** | **Cuculiformes** | **Cuculidae** | ***Crotophaga sulcirostris*** |
| 8 | Harris's Hawk | Accipitriformes | Accipitridae | *Parabuteo unicinctus* |
| 8 | Limpkin | Gruiformes | Aramidae | *Aramus guarauna* |
| **8** | **Long-tailed Manakin** | **Passeriformes** | **Pipridae** | ***Chiroxiphia linearis*** |
| 8 | Nutting's Flycatcher | Passeriformes | Tyrannidae | *Myiarchus nuttingi* |
| 8 | Ruddy Ground-Dove | Columbiformes | Columbidae | *Columbina talpacoti* |
| 8 | Red-legged Honeycreeper | Passeriformes | Thraupidae | *Cyanerpes cyaneus* |
| 8 | Rufous-tailed Hummingbird | Apodiformes | Trochilidae | *Amazilia tzacatl* |
| **8** | **White-throated Magpie-Jay** | **Passeriformes** | **Corvidae** | ***Calocitta formosa*** |
| 9 | Anhinga | Suliformes | Anhingidae | *Anhinga anhinga* |
| **9** | **Clay-colored Thrush** | **Passeriformes** | **Turdidae** | ***Turdus grayi*** |
| **9** | **Groove-billed Ani** | **Cuculiformes** | **Cuculidae** | ***Crotophaga sulcirostris*** |

| Survey version | English name | Order | Family | Scientific name |
|---|---|---|---|---|
| 9 | Giant Cowbird | Passeriformes | Icteridae | *Molothrus oryzivorus* |
| 9 | Hook-billed Kite | Accipitriformes | Accipitridae | *Chondrohierax uncinatus* |
| 9 | Little Blue Heron | Pelecaniformes | Ardeidae | *Egretta caerulea* |
| **9** | **Long-tailed Manakin** | **Passeriformes** | **Pipridae** | ***Chiroxiphia linearis*** |
| 9 | Ochre-bellied Flycatcher | Passeriformes | Tyrannidae | *Mionectes oleaginus* |
| 9 | Purple Gallinule | Gruiformes | Rallidae | *Porphyrio martinicus* |
| 9 | Ruddy Quail-Dove | Columbiformes | Columbidae | *Geotrygon montana* |
| 9 | Steely-vented Hummingbird | Apodiformes | Trochilidae | *Amazilia saucerrottei* |
| 9 | Variable Seedeater | Passeriformes | Emberizidae | *Sporophila corvina* |
| **9** | **White-throated Magpie-Jay** | **Passeriformes** | **Corvidae** | ***Calocitta formosa*** |
| **10** | **Clay-colored Thrush** | **Passeriformes** | **Turdidae** | ***Turdus grayi*** |
| 10 | Neotropic Cormorant | Suliformes | Phalacrocoracidae | *Phalacrocorax brasilianus* |
| **10** | **Groove-billed Ani** | **Cuculiformes** | **Cuculidae** | ***Crotophaga sulcirostris*** |
| 10 | Russet-naped Wood-Rail | Gruiformes | Rallidae | *Aramides albiventris* |
| 10 | Stripe-throated Hermit | Apodiformes | Trochilidae | *Phaethornis striigularis* |
| **10** | **Long-tailed Manakin** | **Passeriformes** | **Pipridae** | ***Chiroxiphia linearis*** |
| 10 | Melodious Blackbird | Passeriformes | Icteridae | *Dives dives* |
| 10 | Pearl Kite | Accipitriformes | Accipitridae | *Gampsonyx swainsonii* |
| 10 | Piratic Flycatcher | Passeriformes | Tyrannidae | *Legatus leucophaius* |
| 10 | Snowy Egret | Pelecaniformes | Ardeidae | *Egretta thula* |
| 10 | White-collared Seedeater | Passeriformes | Emberizidae | *Sporophila torqueola* |
| 10 | White-tipped Dove | Columbiformes | Columbidae | *Leptotila verreauxi* |
| **10** | **White-throated Magpie-Jay** | **Passeriformes** | **Corvidae** | ***Calocitta formosa*** |
| **11** | **Clay-colored Thrush** | **Passeriformes** | **Turdidae** | ***Turdus grayi*** |
| 11 | Common Pauraque | Caprimulgiformes | Caprimulgidae | *Nyctidromus albicollis* |
| **11** | **Groove-billed Ani** | **Cuculiformes** | **Cuculidae** | ***Crotophaga sulcirostris*** |
| 11 | Little Tinamou | Tinamiformes | Tinamidae | *Crypturellus soui* |
| **11** | **Long-tailed Manakin** | **Passeriformes** | **Pipridae** | ***Chiroxiphia linearis*** |
| 11 | Montezuma Oropendola | Passeriformes | Icteridae | *Psarocolius montezuma* |
| 11 | Plumbeous Kite | Accipitriformes | Accipitridae | *Ictinia plumbea* |
| 11 | Royal Flycatcher | Passeriformes | Tyrannidae | *Onychorhynchus coronatus* |
| 11 | Spotted Rail | Gruiformes | Rallidae | *Pardirallus maculatus* |
| 11 | Tricolored Heron | Pelecaniformes | Ardeidae | *Egretta tricolor* |
| **11** | **White-throated Magpie-Jay** | **Passeriformes** | **Corvidae** | ***Calocitta formosa*** |
| 11 | White-winged Dove | Columbiformes | Columbidae | *Zenaida asiatica* |
| 11 | Yellow-faced Grassquit | Passeriformes | Thraupidae | *Tiaris olivaceus* |
| 12 | Amazon Kingfisher | Coraciiformes | Alcedinidae | *Chloroceryle amazona* |

| Survey version | English name | Order | Family | Scientific name |
|---|---|---|---|---|
| 12 | Black-crowned Tityra | Passeriformes | Tityridae | *Tityra inquisitor* |
| 12 | Brown Pelican | Pelecaniformes | Pelecanidae | *Pelecanus occidentalis* |
| **12** | **Great-tailed Grackle** | **Passeriformes** | **Icteridae** | ***Quiscalus mexicanus*** |
| **12** | **Keel-billed Toucan** | **Piciformes** | **Ramphastidae** | ***Ramphastos sulfuratus*** |
| 12 | Lesser Nighthawk | Caprimulgiformes | Caprimulgidae | *Chordeiles acutipennis* |
| 12 | Northern Potoo | Caprimulgiformes | Nyctibiidae | *Nyctibius jamaicensis* |
| **12** | **Orange-chinned Parakeet** | **Psittaciformes** | **Psittacidae** | ***Brotogeris jugularis*** |
| **12** | **Rufous-naped Wren** | **Passeriformes** | **Troglodytidae** | ***Campylorhynchus rufinucha*** |
| 12 | Roadside Hawk | Accipitriformes | Accipitridae | *Rupornis magnirostris* |
| 12 | Red-winged Blackbird | Passeriformes | Icteridae | *Agelaius phoeniceus* |
| 12 | Slate-headed Tody-Flycatcher | Passeriformes | Tyrannidae | *Poecilotriccus sylvia* |
| 12 | Thicket Tinamou | Tinamiformes | Tinamidae | *Crypturellus cinnamomeus* |
| 13 | Blue-black Grosbeak | Passeriformes | Cardinalidae | *Cyanocompsa cyanoides* |
| 13 | Black-headed Trogon | Trogoniformes | Trogonidae | *Trogon melanocephalus* |
| 13 | Black Vulture | Accipitriformes | Cathartidae | *Coragyps atratus* |
| 13 | Green Kingfisher | Coraciiformes | Alcedinidae | *Chloroceryle americana* |
| **13** | **Great-tailed Grackle** | **Passeriformes** | **Icteridae** | ***Quiscalus mexicanus*** |
| **13** | **Keel-billed Toucan** | **Piciformes** | **Ramphastidae** | ***Ramphastos sulfuratus*** |
| 13 | Masked Tityra | Passeriformes | Tityridae | *Tityra semifasciata* |
| **13** | **Orange-chinned Parakeet** | **Psittaciformes** | **Psittacidae** | ***Brotogeris jugularis*** |
| **13** | **Rufous-naped Wren** | **Passeriformes** | **Troglodytidae** | ***Campylorhynchus rufinucha*** |
| 13 | Roseate Spoonbill | Ciconiiformes | Threskiornithidae | *Platalea ajaja* |
| 13 | Social Flycatcher | Passeriformes | Tyrannidae | *Myiozetetes similis* |
| 13 | Spot-breasted Oriole | Passeriformes | Icteridae | *Icterus pectoralis* |
| 13 | Short-tailed Hawk | Accipitriformes | Accipitridae | *Buteo brachyurus* |
| 14 | Elegant Trogon | Trogoniformes | Trogonidae | *Trogon elegans* |
| **14** | **Great-tailed Grackle** | **Passeriformes** | **Icteridae** | ***Quiscalus mexicanus*** |
| **14** | **Keel-billed Toucan** | **Piciformes** | **Ramphastidae** | ***Ramphastos sulfuratus*** |
| 14 | King Vulture | Accipitriformes | Cathartidae | *Sarcoramphus papa* |
| **14** | **Orange-chinned Parakeet** | **Psittaciformes** | **Psittacidae** | ***Brotogeris jugularis*** |
| 14 | Red-crowned Ant-tanager | Passeriformes | Thraupidae | *Habia rubica* |
| 14 | Ringed Kingfisher | Coraciiformes | Alcedinidae | *Megaceryle torquata* |
| **14** | **Rufous-naped Wren** | **Passeriformes** | **Troglodytidae** | ***Campylorhynchus rufinucha*** |
| 14 | Rose-throated Becard | Passeriformes | Tityridae | *Pachyramphus aglaiae* |
| 14 | Streak-backed Oriole | Passeriformes | Icteridae | *Icterus pustulatus* |
| 14 | Snail Kite | Accipitriformes | Accipitridae | *Rostrhamus sociabilis* |
| 14 | Streaked Flycatcher | Passeriformes | Tyrannidae | *Myiodynastes maculatus* |

| Survey version | English name | Order | Family | Scientific name |
|---|---|---|---|---|
| 14 | White Ibis | Pelecaniformes | Threskiornithidae | *Eudocimus albus* |
| 15 | Blue-crowned Motmot | Coraciiformes | Momotidae | *Momotus momota* |
| 15 | Banded Wren | Passeriformes | Troglodytidae | *Thryothorus pleurostictus* |
| **15** | **Great-tailed Grackle** | **Passeriformes** | **Icteridae** | ***Quiscalus mexicanus*** |
| **15** | **Keel-billed Toucan** | **Piciformes** | **Ramphastidae** | ***Ramphastos sulfuratus*** |
| **15** | **Orange-chinned Parakeet** | **Psittaciformes** | **Psittacidae** | ***Brotogeris jugularis*** |
| 15 | Olive Sparrow | Passeriformes | Passerellidae | *Arremonops rufivirgatus* |
| **15** | **Rufous-naped Wren** | **Passeriformes** | **Troglodytidae** | ***Campylorhynchus rufinucha*** |
| 15 | Stub-tailed Spadebill | Passeriformes | Tyrannidae | *Platyrinchus cancrominus* |
| 15 | Turkey Vulture | Accipitriformes | Cathartidae | *Cathartes aura* |
| 15 | Gartered Trogon | Trogoniformes | Trogonidae | *Trogon caligatus* |
| 15 | White-necked Puffbird | Galbuliformes | Bucconidae | *Notharchus hyperrhynchus* |
| 15 | White-tailed Hawk | Accipitriformes | Accipitridae | *Geranoaetus albicaudatus* |
| 15 | Yellow-billed Cacique | Passeriformes | Icteridae | *Amblycercus holosericeus* |
| 16 | Double-striped Thick-knee | Charadriiformes | Burhinidae | *Burhinus bistriatus* |
| 16 | Gray-crowned Yellowthroat | Passeriformes | Parulidae | *Geothlypis poliocephala* |
| **16** | **Great-tailed Grackle** | **Passeriformes** | **Icteridae** | ***Quiscalus mexicanus*** |
| 16 | Hoffmann's Woodpecker | Piciformes | Picidae | *Melanerpes hoffmannii* |
| 16 | House Wren | Passeriformes | Troglodytidae | *Troglodytes aedon* |
| **16** | **Keel-billed Toucan** | **Piciformes** | **Ramphastidae** | ***Ramphastos sulfuratus*** |
| **16** | **Orange-chinned Parakeet** | **Psittaciformes** | **Psittacidae** | ***Brotogeris jugularis*** |
| **16** | **Rufous-naped Wren** | **Passeriformes** | **Troglodytidae** | ***Campylorhynchus rufinucha*** |
| 16 | Sulphur-bellied Flycatcher | Passeriformes | Tyrannidae | *Myiodynastes luteiventris* |
| 16 | Stripe-headed Sparrow | Passeriformes | Passerellidae | *Peucaea ruficauda* |
| 16 | Turquoise-browed Motmot | Coraciiformes | Momotidae | *Eumomota superciliosa* |
| 16 | White-tailed Kite | Accipitriformes | Accipitridae | *Elanus leucurus* |
| 17 | Black-bellied Plover | Charadriiformes | Charadriidae | *Pluvialis squatarola* |
| 17 | Tricolored Munia | Passeriformes | Estrildidae | *Lonchura malacca* |
| **17** | **Great-tailed Grackle** | **Passeriformes** | **Icteridae** | ***Quiscalus mexicanus*** |
| **17** | **Keel-billed Toucan** | **Piciformes** | **Ramphastidae** | ***Ramphastos sulfuratus*** |
| 17 | Lesser Ground-Cuckoo | Cuculiformes | Cuculidae | *Morococcyx erythropygus* |
| 17 | Lineated Woodpecker | Piciformes | Picidae | *Dryocopus lineatus* |
| **17** | **Orange-chinned Parakeet** | **Psittaciformes** | **Psittacidae** | ***Brotogeris jugularis*** |
| 17 | Plain wren | Passeriformes | Troglodytidae | *Cantorchilus modestus* |
| 17 | Rufous-capped Warbler | Passeriformes | Parulidae | *Basileuterus rufifrons* |
| **17** | **Rufous-naped Wren** | **Passeriformes** | **Troglodytidae** | ***Campylorhynchus rufinucha*** |
| 17 | Tropical Kingbird | Passeriformes | Tyrannidae | *Tyrannus melancholicus* |

| Survey version | English name | Order | Family | Scientific name |
|---|---|---|---|---|
| 17 | Zone-tailed Hawk | Accipitriformes | Accipitridae | *Buteo albonotatus* |
| 18 | Blue Grosbeak | Passeriformes | Cardinalidae | *Passerina caerulea* |
| **18** | **Great-tailed Grackle** | **Passeriformes** | **Icteridae** | ***Quiscalus mexicanus*** |
| **18** | **Keel-billed Toucan** | **Piciformes** | **Ramphastidae** | ***Ramphastos sulfuratus*** |
| **18** | **Orange-chinned Parakeet** | **Psittaciformes** | **Psittacidae** | ***Brotogeris jugularis*** |
| 18 | Osprey | Accipitriformes | Accipitridae | *Pandion haliaetus* |
| 18 | Pale-billed Woodpecker | Piciformes | Picidae | *Campephilus guatemalensis* |
| 18 | Rufous-and-white Wren | Passeriformes | Troglodytidae | *Thryophilus rufalbus* |
| **18** | **Rufous-naped Wren** | **Passeriformes** | **Troglodytidae** | ***Campylorhynchus rufinucha*** |
| 18 | Scrub Euphonia | Passeriformes | Thraupidae | *Euphonia affinis* |
| 18 | Southern Lapwing | Charadriiformes | Charadriidae | *Vanellus chilensis* |
| 18 | Squirrel Cuckoo | Cuculiformes | Cuculidae | *Piaya cayana* |
| 18 | Western/Eastern Wood-Pewee | Passeriformes | Tyrannidae | *Contopus sordilus/virens* |
| 19 | Black-bellied Whistling-Duck | Anseriformes | Anatidae | *Dendrocygna autumnalis* |
| 19 | Collared Aracari | Piciformes | Ramphastidae | *Pteroglossus torquatus* |
| **19** | **Great-tailed Grackle** | **Passeriformes** | **Icteridae** | ***Quiscalus mexicanus*** |
| 19 | House Sparrow | Passeriformes | Passeridae | *Passer domesticus* |
| **19** | **Keel-billed Toucan** | **Piciformes** | **Ramphastidae** | ***Ramphastos sulfuratus*** |
| 19 | Northern Jacana | Gruiformes | Rallidae | *Jacana spinosa* |
| 19 | Orange-billed Nightingale Thrush | Passeriformes | Turdidae | *Catharus aurantiirostris* |
| **19** | **Orange-chinned Parakeet** | **Psittaciformes** | **Psittacidae** | ***Brotogeris jugularis*** |
| **19** | **Rufous-naped Wren** | **Passeriformes** | **Troglodytidae** | ***Campylorhynchus rufinucha*** |
| 19 | Striped Cuckoo | Cuculiformes | Cuculidae | *Tapera naevia* |
| 19 | White-winged Becard | Passeriformes | Tityridae | *Pachyramphus polychopterus* |
| 19 | Yellow-crowned Euphonia | Passeriformes | Thraupidae | *Euphonia luteicapilla* |
| 20 | Alder Flycatcher | Passeriformes | Tyrannidae | *Empidonax alnorum* |
| 20 | Bat Falcon | Falconiformes | Falconidae | *Falco rufigularis* |
| 20 | Black-necked Stilt | Charadriiformes | Recurvirostridae | *Himantopus mexicanus* |
| 20 | Crimson-fronted Parakeet | Psittaciformes | Psittacidae | *Psittacara finschi* |
| 20 | Fulvous Whistling Duck | Anseriformes | Anatidae | *Dendrocygna bicolor* |
| **20** | **Great-tailed Grackle** | **Passeriformes** | **Icteridae** | ***Quiscalus mexicanus*** |
| **20** | **Keel-billed Toucan** | **Piciformes** | **Ramphastidae** | ***Ramphastos sulfuratus*** |
| 20 | Long-billed Gnatwren | Passeriformes | Polioptilidae | *Ramphocaenus melanurus* |
| **20** | **Orange-chinned Parakeet** | **Psittaciformes** | **Psittacidae** | ***Brotogeris jugularis*** |
| **20** | **Rufous-naped Wren** | **Passeriformes** | **Troglodytidae** | ***Campylorhynchus rufinucha*** |
| 20 | Yellow-bellied Elaenia | Passeriformes | Tyrannidae | *Elaenia flavogaster* |
| 20 | Yellow-throated Euphonia | Passeriformes | Thraupidae | *Euphonia hirundinacea* |

| Survey version | English name | Order | Family | Scientific name |
|---|---|---|---|---|
| 21 | Boat-billed Flycatcher | Passeriformes | Tyrannidae | *Megarynchus pitangua* |
| 21 | Collared Forest-Falcon | Falconiformes | Falconidae | *Micrastur semitorquatus* |
| **21** | **Great-tailed Grackle** | **Passeriformes** | **Icteridae** | ***Quiscalus mexicanus*** |
| 21 | Ivory-billed Woodcreeper | Passeriformes | Furnariidae | *Xiphorhynchus flavigaster* |
| **21** | **Keel-billed Toucan** | **Piciformes** | **Ramphastidae** | ***Ramphastos sulfuratus*** |
| 21 | Muscovy Duck | Anseriformes | Anatidae | *Cairina moschata* |
| **21** | **Orange-chinned Parakeet** | **Psittaciformes** | **Psittacidae** | ***Brotogeris jugularis*** |
| 21 | Orange-fronted Parakeet | Psittaciformes | Psittacidae | *Eupsittula canicularis* |
| **21** | **Rufous-naped Wren** | **Passeriformes** | **Troglodytidae** | ***Campylorhynchus rufinucha*** |
| 21 | Tropical Gnatcatcher | Passeriformes | Polioptilidae | *Polioptila plumbea* |
| 21 | Whimbrel | Charadriiformes | Scolopacidae | *Numenius phaeopus* |
| 21 | Yellow-olive Flycatcher | Passeriformes | Tyrannidae | *Tolmomyias sulphurescens* |
| 22 | Northern Barred-Woodcreeper | Passeriformes | Furnariidae | *Dendrocolaptes sanctithomae* |
| 22 | Black Swift | Apodiformes | Apodidae | *Cypseloides niger* |
| 22 | Bright-rumped Attila | Passeriformes | Tyrannidae | *Attila spadiceus* |
| 22 | Crested Caracara | Accipitriformes | Accipitridae | *Caracara cheriway* |
| **22** | **Great-tailed Grackle** | **Passeriformes** | **Icteridae** | ***Quiscalus mexicanus*** |
| 22 | Jabiru | Ciconiiformes | Ciconiidae | *Jabiru mycteria* |
| **22** | **Keel-billed Toucan** | **Piciformes** | **Ramphastidae** | ***Ramphastos sulfuratus*** |
| 22 | Lesser Greenlet | Passeriformes | Vireonidae | *Pachysylvia decurtata* |
| **22** | **Orange-chinned Parakeet** | **Psittaciformes** | **Psittacidae** | ***Brotogeris jugularis*** |
| 22 | Red-lored Parrot | Psittaciformes | Psittacidae | *Amazona autumnalis* |
| **22** | **Rufous-naped Wren** | **Passeriformes** | **Troglodytidae** | ***Campylorhynchus rufinucha*** |
| 22 | White-lored Gnatcatcher | Passeriformes | Polioptilidae | *Polioptila albiloris* |

#### 2 INFORMATION FOR THE SONG AND AUDIO RECORDINGS USED IN THE SURVEY

**Table S2. Information regarding the recordings used in the survey downloaded from the Xeno-canto website**

| Species # | English name | Recording author | Recording Website | Recording country | Recording type | Xeno-canto ID number |
|---|---|---|---|---|---|---|
| 1 | Amazon Kingfisher | Andrew Spencer | Xeno-Canto | Panama | Song | XC31704 |
| 2 | Alder Flycatcher | Oliver Komar | Xeno-canto | Honduras | Call | XC252894 |
| 3 | Anhinga | Mike Nelson | Xeno-canto | Costa Rica | Call | XC166158 |
| 4 | Barred Antshrike | Dan Mennill | Xeno-Canto | Costa Rica | Song | XC572 |
| 5 | Bat Falcon | Patrick ODonnell | Xeno-Canto | Costa Rica | Call | XC338634 |
| 6 | Barn Owl | Edwin Calderón | Xeno-Canto | El Salvador | Call | XC319074 |
| 7 | Barn Swallow | Fernando Mondaca Fernandez | Xeno-Canto | Mexico | Song | XC384074 |
| 8 | Northern Barred-Woodcreeper | Peter Boesman | Xeno-Canto | Panama | Song | XC271225 |
| 9 | Boat-billed Flycatcher | Oscar Ramírez Alán | Xeno-Canto | Costa Rica | Call, alarm call | XC169329 |
| 10 | Blue-black Grosbeak | George Wagner | Xeno-Canto | Costa Rica | Song | XC278977 |
| 11 | Blue-black Grassquit | Peter Boesman | Xeno-canto | Costa Rica | Song | XC274562 |
| 12 | Boat-billed Heron | Mike Nelson | Xeno-canto | Costa Rica | Call | XC166160 |
| 13 | Black-bellied Plover | John V Moore | Xeno-canto | Ecuador | Take-off and flight calls | XC257551 |
| 14 | Black-bellied Whistling-Duck | Oswaldo Cortés | Xeno-canto | Colombia | Song | XC86111 |
| 15 | Brown-crested Flycatcher | Albert Lastukhin | Xeno-canto | Costa Rica | Call | XC378411 |
| 16 | Blue-crowned Motmot | Manuel Grosselet | Xeno-Canto | Mexico | Song | XC319766 |
| 17 | Black-crowned Night-Heron | Johana Zuluaga-Bonilla | Xeno-Canto | Costa Rica | Song | XC153095 |
| 18 | Black-crowned Tityra | Paul Smith | Xeno-Canto | Paraguay | Call | XC15928 |
| 19 | Blue-gray Tanager | Peter Boesman | Xeno-Canto | Costa Rica | Song | XC274501 |
| 20 | Black-headed Trogon | David Bradley | Xeno-canto | Costa Rica | Song | XC6763 |
| 21 | Blue Grosbeak | Scott Olmstead | Xeno-Canto | Costa Rica | Song | XC202749 |
| 22 | Blue Ground Dove | Peter Boesman | Xeno-canto | Costa Rica | Song | XC221154 |
| 23 | Black Swift | Nathan Peiplow | Xeno-Canto | Mexico | Call | XC255569 |
| 24 | Black Vulture | Marcelo Araya-Salas | Xeno-canto | Costa Rica | Call | XC154016 |
| 25 | Black-necked Stilt | Albert Lastukhin | Xeno-Canto | Costa Rica | Call | XC383317 |
| 26 | Banded Wren | Michelle L Hall | Xeno-Canto | Costa Rica | Song | XC210808 |
| 27 | Bright-rumped Attila | Mary Beth Stowe | Xeno-canto | Costa Rica | Song | XC338413 |
| 28 | Bronzed Cowbird | Manuel Grosselet | Xeno-canto | Mexico | Begging call | XC382045 |
| 29 | Brown Pelican | Judith Priam | Xeno-canto | France | Call, juvenile | XC357928 |

| Species # | English name | Recording author | Recording Website | Recording country | Recording type | Xeno-canto ID number |
|---|---|---|---|---|---|---|
| 30 | Buff-throated Saltator | Peter Boesman | Xeno-Canto | Costa Rica | Song | XC274021 |
| 31 | Bare-throated Tiger-Heron | Albert Lastukhin | Xeno-canto | Costa Rica | Flight call | XC371459 |
| 32 | Green Heron | Albert Lastukhin | Xeno-Canto | Costa Rica | Call | XC372645 |
| 33 | Cattle Egret | Albert Lastukhin | Xeno-canto | Costa Rica | Flight call | XC371472 |
| 34 | Canivet's Emerald | Paul Driver | Xeno-Canto | Costa Rica | Call | XC137688 |
| 35 | Clay-colored Robin | Peter Boesman | Xeno-canto | Costa Rica | Song | XC274554 |
| 36 | Crimson-fronted Parakeet | Don Jones | Xeno-canto | Costa Rica | Flight call | XC15529 |
| 37 | Tricolored Munia | Juan Pablo López O | Xeno-Canto | Colombia | Call | XC49035 |
| 38 | Cinnamon Hummingbird | Marcelo Araya-Salas | Xeno-Canto | Costa Rica | Song | XC154050 |
| 39 | Collared Aracari | Albert Lastukhin | Xeno-canto | Costa Rica | Call | XC376487 |
| 40 | Common Black-Hawk | Albert Lastukhin | Xeno-Canto | Costa Rica | Call | XC372644 |
| 41 | Collared Forest-Falcon | Oscar Ramírez Alán | Xeno-Canto | Costa Rica | Song | XC175262 |
| 42 | Common Ground-Dove | Manuel Grosselet | Xeno-canto | Mexico | Song | XC62707 |
| 43 | Common Pauraque | Mike Nelson | Xeno-Canto | Costa Rica | Song | XC166197 |
| 44 | Neotropic Cormorant | Peter Boesman | Xeno-canto | Brazil | Call | XC227630 |
| 45 | Crested Bobwhite | Ken Allaire | Xeno-canto | Panama | Call | XC6412 |
| 46 | Crested Caracara | Oswaldo Cortés | Xeno-canto | Colombia | Song | XC92402 |
| 47 | Crane Hawk | Peter Boesman | Xeno-canto | Venezuela | Call | XC223472 |
| 48 | Common Tody-Flycatcher | Oscar Ramírez Alán | Xeno-canto | Costa Rica | Song | XC180510 |
| 49 | Dusky-capped Flycatcher | Mary Beth Stowe | Xeno-Canto | Costa Rica | Call | XC338420 |
| 50 | Double-striped Thick-knee | Carlos Funes | Xeno-Canto | El Salvador | Call | XC45014 |
| 51 | Double-toothed Kite | Sander Bolt | Xeno-canto | Panama | Call | XC112868 |
| 52 | Eastern Meadowlark | Paul Driver | Xeno-Canto | Costa Rica | Song | XC140147 |
| 53 | Elegant Trogon | David Bradley | Xeno-Canto | Costa Rica | Song | XC6770 |
| 54 | Ferruginous Pygmy-Owl | Robin Carter | Xeno-canto | Costa Rica | Call | XC6797 |
| 55 | Fulvous Whistling Duck | Manuel Grosselet | Xeno-canto | Mexico | Song | XC325493 |
| 56 | Groove-billed Ani | Albert Lastukhin | Xeno-canto | Costa Rica | Flight call | XC372871 |
| 57 | Great Black Hawk | John V. Moore | Xeno-canto | Ecuador | Call | XC257362 |
| 58 | Gray-breasted Martin | Danny Zapata-Henao | Xeno-Canto | Colombia | Song | XC213474 |
| 59 | Gray-crowned Yellowthroat | Bobby Wilcox | Xeno-Canto | Costa Rica | Song | XC332260 |

| Species # | English name | Recording author | Recording Website | Recording country | Recording type | Xeno-canto ID number |
|---|---|---|---|---|---|---|
| 60 | Gray-headed Chachalaca | Peter Boesman | Xeno-canto | Panama | Call | XC271478 |
| 61 | Gray-headed Dove | Daniel Lane | Xeno-Canto | Belize | Song | XC28435 |
| 62 | Gray-headed Kite | Peter Boesman | Xeno-canto | Costa Rica | Song | XC274517 |
| 63 | Giant Cowbird | Ken Allaire | Xeno-Canto | Panama | Call | XC60672 |
| 64 | Green Kingfisher | Andrew Spencer | Xeno-Canto | Panama | Call | XC31834 |
| 65 | Russet-naped Wood-Rail | Oscar Campbell | Xeno-canto | Costa Rica | Call | XC278669 |
| 66 | Great Curassow | Johan Chaves | Xeno-canto | Costa Rica | Call | XC370071 |
| 67 | Great Egret | Doug Knapp | Xeno-canto | Nicaragua | Call | XC10884 |
| 68 | Greenish Elaenia | Patrick ODonnell | Xeno-Canto | Costa Rica | Call | XC246645 |
| 69 | Gray Hawk | Peter Boesman | Xeno-Canto | Costa Rica | Call | XC274482 |
| 70 | Great Kiskadee | Peter Boesman | Xeno-canto | Costa Rica | Song | XC274495 |
| 71 | Grayish Saltator | Marcelo Araya-Salas | Xeno-canto | Costa Rica | Song | XC154186 |
| 72 | Gray-headed Tanager | Robin Carter | Xeno-Canto | Costa Rica | Song | XC1015 |
| 73 | Great-tailed Grackle | Peter Boesman | Xeno-canto | Costa Rica | Call | XC274570 |
| 74 | Harris's Hawk | Manuel Grosselet | Xeno-canto | Mexico | Call | XC376776 |
| 75 | Hook-billed Kite | Mayron McKewy Mejía | Xeno-canto | Honduras | Call | XC121472 |
| 76 | House Sparrow | Guillermo Funes | Xeno-Canto | El Salvador | Call | XC306629 |
| 77 | Hoffmann's Woodpecker | Jim Holmes | Xeno-Canto | Costa Rica | Call | XC192887 |
| 78 | House Wren | Bobby Wilcox | Xeno-Canto | Costa Rica | Song | XC332269 |
| 79 | Ivory-billed Woodcreeper | Andrés Jimenez | Xeno-Canto | Costa Rica | Song | XC132737 |
| 80 | Inca Dove | Johan Chaves | Xeno-canto | Costa Rica | Call, Song | XC384746 |
| 81 | Jabiru | Miguel Castelino | Xeno-canto | Argentina | Call | XC60513 |
| 82 | Keel-billed Toucan | Dan Mennill | Xeno-canto | Costa Rica | Song | XC588 |
| 83 | King Vulture | NA | NA | NA | NA | NA |
| 84 | Laughing Falcon | Peter Boesman | Xeno-Canto | Costa Rica | Song | XC224106 |
| 85 | Long-billed Gnatwren | Tom Stevens | Xeno-Canto | Costa Rica | Song | XC93316 |
| 86 | Little Blue Heron | Doug Knapp | Xeno-canto | Nicaragua | Call | XC10883 |
| 87 | Least Grebe | Ken Allaire | Xeno-canto | Panama | Call | XC24210 |
| 88 | Lesser Greenlet | Jim Holmes | Xeno-Canto | Costa Rica | Song | XC192885 |
| 89 | Lesser Nighthawk | John V Moore | Xeno-canto | Ecuador | Song | XC257766 |
| 90 | Lesser Ground-Cuckoo | Richard Garrigues | Xeno-Canto | Costa Rica | Call | XC5797 |
| 91 | Stripe-throated Hermit | Peter Boesman | Xeno-Canto | Costa Rica | Song | XC274600 |
| 92 | Limpkin | Melvin Bonilla | Xeno-canto | El Salvador | Alarm call | XC195696 |
| 93 | Little Tinamou | Sander Bot | Xeno-Canto | Costa Rica | Song | XC112977 |
| 94 | Lineated Woodpecker | Oswaldo Cortes | Xeno-Canto | Colombia | Song | XC106275 |

| Species # | English name | Recording author | Recording Website | Recording country | Recording type | Xeno-canto ID number |
|---|---|---|---|---|---|---|
| 95 | Lesser Swallow-tailed Swift | Marco Cruz | Xeno-Canto | Brazil | Song | XC199071 |
| 96 | Long-tailed Manakin | Dugan Maynard | Xeno-canto | Costa Rica | Song | XC147697 |
| 97 | Green-breasted Mango | Mary Beth Stowe | Xeno-canto | Costa Rica | Call | XC338421 |
| 98 | Mangrove Swallow | Peter Boesman | Xeno-Canto | Costa Rica | Call | XC274357 |
| 99 | Masked Tityra | Robin Carter | Xeno-Canto | Costa Rica | Song | XC1020 |
| 100 | Melodious Blackbird | Albert Lastukhin | Xeno-Canto | Costa Rica | Call/song | XC380880 |
| 101 | Muscovy Duck | Oscar Ramírez Alán | Xeno-Canto | Costa Rica | Alarm call | XC165832 |
| 102 | Montezuma Oropendola | Peter Boesman | Xeno-canto | Costa Rica | Song | XC274470 |
| 103 | Mottled Owl | Micah Reigner | Xeno-canto | Mexico | Song | XC358832 |
| 104 | Northern Beardless-Tyrannulet | Albert Lastukhin | Xeno-Canto | Costa Rica | Song | XC372653 |
| 105 | Northern Bentbill | Mike Nelson | Xeno-Canto | Costa Rica | Song | XC166194 |
| 106 | Northern Jacana | Peter Boesman | Xeno-Canto | Costa Rica | Call | XC274390 |
| 107 | Northern Potoo | Peter Boesman | Xeno-canto | Mexico | Song | XC226750 |
| 108 | Nutting's Flycatcher | Orlando Jarquín G | Xeno-Canto | Nicaragua | Song | XC284440 |
| 109 | Ochre-bellied Flycatcher | Robin Carter | Xeno-canto | Costa Rica | Song | XC1022 |
| 110 | Orange-billed Nightingale-Thrush | Mike Nelson | Xeno-canto | Costa Rica | Song | XC106715 |
| 111 | Orange-chinned Parakeet | Albert Lastukhin | Xeno-Canto | Costa Rica | Call | XC371455 |
| 112 | Orange-fronted Parakeet | Peter Boesman | Xeno-Canto | Costa Rica | Call | XC274581 |
| 113 | Olive Sparrow | Jorge Gabriel Campos | Xeno-Canto | Costa Rica | Call, song | XC328940 |
| 114 | Olivaceous Woodcreeper | Rick Bowers | Xeno-Canto | Costa Rica | Song | XC29624 |
| 115 | Osprey | Doug Knapp | Xeno-canto | Nicaragua | Call | XC10887 |
| 116 | Pacific Screech-Owl | Jim Holmes | Xeno-canto | Costa Rica | Song | XC189030 |
| 117 | Palm Tanager | Peter Boesman | Xeno-Canto | Panama | Song | XC271627 |
| 118 | Plain-breasted Ground-Dove | Peter Boesman | Xeno-Canto | Venezuela | Song | XC221413 |
| 119 | Pale-billed Woodpecker | Robin Carter | Xeno-Canto | Costa Rica | Song | XC1106 |
| 120 | Plain-capped Starthroat | Andrew Spencer | Xeno-Canto | Costa Rica | Call | XC191100 |
| 121 | Pearl Kite | Marcos A. Melo | Xeno-canto | Brazil | Male, Song | XC240803 |
| 122 | Piratic Flycatcher | Peter Boesman | Xeno-Canto | Costa Rica | Song | XC274136 |
| 123 | Plain Chachalaca | Jorge Gabriel Campos | Xeno-canto | Costa Rica | Male song | XC328936 |

| Species # | English name | Recording author | Recording Website | Recording country | Recording type | Xeno-canto ID number |
|---|---|---|---|---|---|---|
| 124 | Plumbeous Kite | Mitch Lysinger | Xeno-canto | Ecuador | Flight call | XC260333 |
| 125 | Plain wren | Albert Lastukhin | Xeno-canto | Costa Rica | Song | XC380754 |
| 126 | Purple Gallinule | Mike Nelson | Xeno-Canto | Costa Rica | Call | XC166168 |
| 127 | Rufous-and-white Wren | Paul Driver | Xeno-canto | Costa Rica | Song | XC137724 |
| 128 | Red-billed Pigeon | Ryan P. O'Donnell | Xeno-Canto | Costa Rica | Song | XC72518 |
| 129 | Rufous-browed Peppershrike | Paul Driver | Xeno-canto | Costa Rica | Song | XC137731 |
| 130 | Red-crowned Ant-tanager | Mike Nelson | Xeno-canto | Panama | Song | XC92094 |
| 131 | Rufous-capped Warbler | David Bradley | Xeno-Canto | Costa Rica | Song | XC7396 |
| 132 | Ruddy Ground-Dove | Guillermo Funes | Xeno-canto | El Salvador | Song | XC198411 |
| 133 | Ringed Kingfisher | Gary Stiles | Xeno-canto | Colombia | Song | XC148288 |
| 134 | Red-lored Parrot | Diego Caiafa | Xeno-canto | Costa Rica | Song | XC195077 |
| 135 | Red-legged Honeycreeper | Gary Stiles | Xeno-canto | Colombia | Song | XC148077 |
| 136 | Rufous-naped Wren | Beatrix Saadi-Varchmin | Xeno-canto | Costa Rica | Song | XC377946 |
| 137 | Royal Flycatcher | Peter Boesman | Xeno-canto | Costa Rica | Call | XC274493 |
| 138 | Roadside Hawk | Iain | Xeno-Canto | Costa Rica | Call | XC233494 |
| 139 | Roseate Spoonbill | Paul Marvin | Xeno-canto | United States | Call | XC161508 |
| 140 | Rose-throated Beccard | Peter Boesman | Xeno-Canto | Costa Rica | Song | XC274224 |
| 141 | Rufous-tailed Hummingbird | Peter Boesman | Xeno-Canto | Costa Rica | Call | XC274522 |
| 142 | Ruddy Quail-Dove | Oswaldo Cortés | Xeno-canto | Colombia | Song | XC96233 |
| 143 | Ruddy Woodcreeper | Niels Krabbe | Xeno-Canto | Colombia | Song | XC235681 |
| 144 | Red-winged Blackbird | Guillermo Funes | Xeno-canto | El Salvador | Song | XC300863 |
| 145 | Sulphur-bellied Flycatcher | Paul Driver | Xeno-Canto | Costa Rica | Call | XC140145 |
| 146 | Streak-backed Oriole | Mario Trejo | Xeno-Canto | El Salvador | Song | XC175349 |
| 147 | Scrub Euphonia | Oscar Ramírez Alán | Xeno-Canto | Costa Rica | Song | XC170081 |
| 148 | Scarlet Macaw | Scott Olmstead | Xeno-canto | Costa Rica | Call | XC374502 |
| 149 | Stripe-headed Sparrow | David Bradley | Xeno-canto | Costa Rica | Song | XC6397 |
| 150 | Slate-headed Tody-Flycatcher | Dan Mennill | Xeno-Canto | Costa Rica | Song | XC655 |
| 151 | Streak-headed Woodcreeper | Peter Boesman | Xeno-Canto | Costa Rica | Song | XC274606 |
| 152 | Snowy Egret | John V Moore | Xeno-canto | Ecuador | Call | XC257313 |
| 153 | Snail Kite | Thore Noernberg | Xeno-canto | Panama | Call | XC127155 |
| 154 | Social Flycatcher | Robin Carter | Xeno-Canto | Costa Rica | Call | XC1028 |
| 155 | Southern Lapwing | Albert Lastukhin | Xeno-canto | Costa Rica | Alarm call | XC375519 |

| Species # | English name | Recording author | Recording Website | Recording country | Recording type | Xeno-canto ID number |
|---|---|---|---|---|---|---|
| 156 | Spot-breasted Oriole | Robin Carter | Xeno-canto | Costa Rica | Song | XC6463 |
| 157 | Spectacled Owl | Olaf Jahn | Xeno-canto | Ecuador | Song | XC261641 |
| 158 | Spotted Rail | Jorge Gabriel Campos | Xeno-Canto | Costa Rica | Song | XC378763 |
| 159 | Squirrel Cuckoo | Mike Nelson | Xeno-canto | Costa Rica | Song | XC107189 |
| 160 | Striped Cuckoo | Peter Boesman | Xeno-Canto | Costa Rica | Song | XC274487 |
| 161 | Streaked Flycatcher | John V Moore | Xeno-Canto | Costa Rica | Call | XC275473 |
| 162 | Short-tailed Hawk | Guillermo Funes | Xeno-canto | El Salvador | Call, song | XC197642 |
| 163 | American Swallow-tailed Kite | Mike Nelson | Xeno-canto | Costa Rica | Call | XC107216 |
| 164 | Stub-tailed Spadebill | Ian Davies | Xeno-canto | Mexico | Song | XC118754 |
| 165 | Steely-vented Hummingbird | Albert Lastukhin | Xeno-Canto | Costa Rica | Call | XC372720 |
| 166 | Turquoise-browed Motmot | Scott Connop | Xeno-canto | Costa Rica | Song | XC108610 |
| 167 | Thicket Tinamou | Jorge Gabriel Campos | Xeno-canto | Costa Rica | Song | XC328942 |
| 168 | Tropical Gnatcatcher | David Bradley | Xeno-Canto | Costa Rica | Song | XC6395 |
| 169 | Tricolored Heron | Doug Knapp | Xeno-canto | Nicaragua | Call | XC10882 |
| 170 | Tropical Kingbird | Paul Driver | Xeno-canto | Costa Rica | Dawn song | XC137689 |
| 171 | Turkey Vulture | Charlie Vogt | Xeno-Canto | Ecuador | Call | XC16325 |
| 172 | Variable Seedeater | Peter Boesman | Xeno-canto | Costa Rica | Call | XC274057 |
| 173 | Vaux's Swift | Allet T Chartier | Xeno-canto | Costa Rica | Call | XC8894 |
| 174 | Gartered Trogon | Peter Boesman | Xeno-Canto | Costa Rica | Song | XC274382 |
| 175 | White-collared Seedeater | Andrew Spencer | Xeno-canto | Costa Rica | Song | XC72319 |
| 176 | White-collared Swift | Peter Boesman | Xeno-canto | Costa Rica | Call | XC274462 |
| 177 | White-fronted Parrot | Robin Carter | Xeno-canto | Costa Rica | Song | XC7831 |
| 178 | White Ibis | Doug Knapp | Xeno-canto | Nicaragua | Call | XC10881 |
| 179 | Whimbrel | Mike Nelson | Xeno-canto | Costa Rica | Call, Song | XC166313 |
| 180 | White-lored Gnatcatcher | Jim Holmes | Xeno-Canto | Costa Rica | Call | XC189094 |
| 181 | White-necked Puffbird | Albert Lastukhin | Xeno-Canto | Costa Rica | Call | XC381137 |
| 182 | Wood Stork | Bernabe Lopez-Lanus | Xeno-canto | Argentina | Call | XC45147 |
| 183 | White-tipped Dove | Dan Mennill | Xeno-canto | Costa Rica | Song | XC661 |
| 184 | White-tailed Hawk | John van Dort | Xeno-canto | Honduras | Call | XC85826 |
| 185 | White-tailed Kite | Manuel Grosselet | Xeno-Canto | Mexico | Call | XC293103 |
| 186 | White-throated Magpie-Jay | Peter Boesman | Xeno-canto | Costa Rica | Call | XC274498 |
| 187 | White-winged Becard | Andrew Spencer | Xeno-canto | Costa Rica | Call | XC72400 |
| 188 | White-winged Dove | Albert Lastukhin | Xeno-Canto | Costa Rica | Song | XC378630 |

| Species # | English name | Recording author | Recording Website | Recording country | Recording type | Xeno-canto ID number |
|---|---|---|---|---|---|---|
| 189 | Western/Eastern Wood-Pewee | Sander Bolt | Xeno-canto | Panama | Song | XC113559 |
| 190 | Yellow-billed Cacique | Scott Olmstead | Xeno-canto | Costa Rica | Song | XC48977 |
| 191 | Yellow-bellied Elaenia | Paul Driver | Xeno-canto | Costa Rica | Song | XC137682 |
| 192 | Yellow-crowned Euphonia | Mike Nelson | Xeno-canto | Costa Rica | Song | XC166837 |
| 193 | Yellow-faced Grassquit | Peter Boesman | Xeno-canto | Costa Rica | Song | XC274459 |
| 194 | Yellow-green Vireo | Jeff Norris | Xeno-Canto | Costa Rica | Song | XC334255 |
| 195 | Yellow-headed Caracara | Mike Nelson | Xeno-canto | Costa Rica | Call | XC166318 |
| 196 | Yellow-naped Parrot | Patrick ODonnell | Xeno-Canto | Costa Rica | Call | XC335408 |
| 197 | Yellow-olive Flycatcher | David Bradley | Xeno-Canto | Costa Rica | Call | XC6837 |
| 198 | Yellow-throated Euphonia | Peter Boesman | Xeno-Canto | Costa Rica | Song | XC274504 |
| 199 | Zone-tailed Hawk | Andrew Spencer | Xeno-canto | Mexico | Call | XC265410 |

### 3 STATISTICAL ANALYSES AND MODEL FITS

#### 3.1 DISSERVICES: MIXED-EFFECTS MODEL

| **Model** | **Disservices = Species + Social group + Species*Social group + (1\|Participant)** |
|---|---|
| AIC= 3688.57; BIC=3824.82 | |

| Type II Anova | | | |
|---|---|---|---|
| **Term** | **Chi-squared** | **Degrees of Freedom** | **P value** |
| Species | 1211.83 | 7 | <2.26E-16 |
| Social group | 56.96 | 2 | 4.29E-13 |
| Species: social group | 12.06 | 14 | <2.2E-16 |

| Term | Estimate | Degrees of freedom | T-value |
|---|---|---|---|
| Intercept | 1.51 | 997 | 11.71 |
| Species Groove-billed Ani (GBAN) | 0.12 | 997 | 0.77 |
| Species Great-tailed Grackle (GTGR) | 1.47 | 997 | 8.5 |
| Species Keel-billed Toucan (KBTO) | 0.15 | 997 | 0.93 |
| Species Long-tailed Manakin (LTMA) | -0.2 | 997 | -1.27 |
| Species Orange-chinned Parakeet (OCPA) | 0.17 | 997 | 1 |
| Species Rufous-naped Wren (RNWR) | -0.1 | 997 | -0.59 |
| Species White-throated Magpie-Jay (WTMJ) | 0.56 | 997 | 3.61 |
| Social group Farmers | -0.3 | 396 | -1.75 |
| Social group Urbanites | -0.11 | 396 | -0.65 |
| GBAN: farmers | 0.76 | 997 | 3.63 |
| GTGR: farmers | 1.43 | 997 | 6.1 |
| KBTO: farmers | 0.26 | 997 | 1.11 |
| LTMA: farmers | 0.09 | 997 | 0.43 |
| OCPA: farmers | 0.84 | 997 | 3.6 |
| RNWR: farmers | 1.01 | 997 | 4.3 |
| WTMJ: farmers | 1.11 | 997 | 5.46 |
| GBAN: urbanites | 1.03 | 997 | 4.82 |
| GTGR: urbanites | 1.19 | 997 | 4.93 |
| KBTO: urbanites | 0.22 | 997 | 0.97 |
| LTMA: urbanites | 0.12 | 997 | 0.53 |
| OCPA: urbanites | 0.64 | 997 | 2.75 |
| RNWR: urbanites | 1.19 | 997 | 4.93 |
| WTMJ: urbanites | 1.06 | 997 | 5.07 |

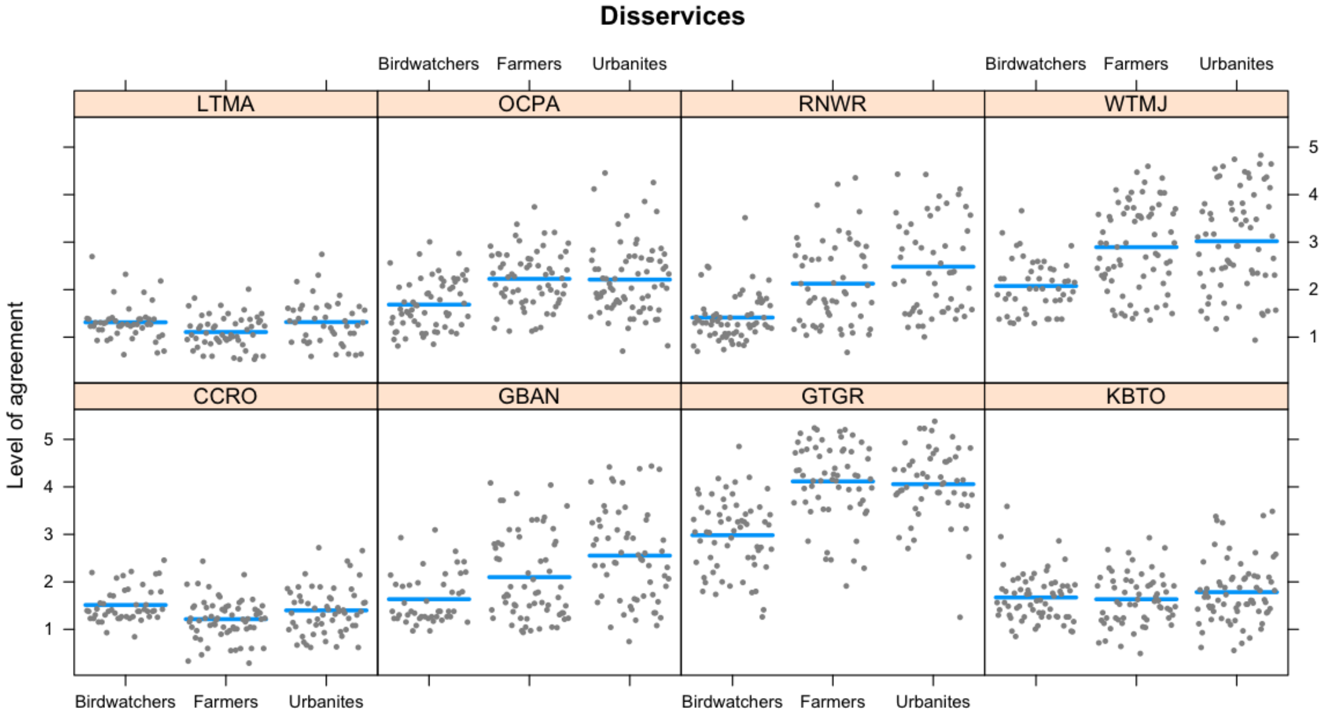

**Figure S1. Model fits for each species and three social groups regarding disservices scores.** Each panel represents a species (LTMA=Long-tailed Manakin, OCPA=Orange-chinned Parakeet, RNWR=Rufous-naped Wren, WTMJ=White-throated Magpie-Jay, CCRO=Clay-colored Thrush, GBAN=Groove-billed Ani, GTGR=Great-tailed Grackle, KBTO=Keel-billed Toucan). Each grey dot represents one person, and the blue lines represent the model.

**Table S3. Tukey HSD pairwise comparisons for disservices scores.**

| **Species** | **Pairwise comparison** | **P value** |
|---|---|---|
| Great-tailed Grackle | Birdwatchers vs. farmers | *p*<0.001*** |
| | Birdwatchers vs. urbanites | *p*=0.001** |
| | Farmers vs. urbanites | *p*>0.05 |
| Orange-chinned Parakeet | Birdwatchers vs. farmers | *p*<0.001*** |
| | Birdwatchers vs. urbanites | *p*=0.002 |
| | Farmers vs. urbanites | *p*>0.05 |
| White-throated Magpie-Jay | Birdwatchers vs. farmers | *p*=0.002** |
| | Birdwatchers vs. urbanites | *p*<0.001*** |
| | Farmers vs. urbanites | *p*>0.05 |
| Rufous-naped Wren | Birdwatchers vs. farmers | *p*<0.001*** |
| | Birdwatchers vs. urbanites | *p*=0.007** |
| | Farmers vs. urbanites | *p*>0.05 |
| Groove-billed Ani | Birdwatchers vs. farmers | *p*<0.001*** |
| | Birdwatchers vs. urbanites | *p*>0.05 |
| | Farmers vs. urbanites | *p*=0.019* |
| Clay-colored Thrush | Birdwatchers vs. farmers | *p*=0.002** |

| Species | Pairwise comparison | P value |
|---|---|---|
| | Birdwatchers vs. urbanites | *p*>0.05 |
| | Farmers vs. urbanites | *p*=0.019* |
| Keel-billed Toucan | Birdwatchers vs. farmers | *p*>0.05 |
| | Birdwatchers vs. urbanites | *p*>0.05 |
| | Farmers vs. urbanites | *p*>0.05 |
| Long-tailed Manakin | Birdwatchers vs. farmers | *p*>0.05 |
| | Birdwatchers vs. urbanites | *p*>0.05 |
| | Farmers vs. urbanites | *p*>0.05 |

*p<0.05, *p<0.01, ***p<0.001

#### 3.2 EDUCATION: MIXED-EFFECTS MODEL

**Model** **Education = Species + Social group + Species*Social group + (1|Participant)**

AIC= 4151.13; BIC=4287.35

| Type II Anova | | | |
|---|---|---|---|
| **Term** | **Chi-squared** | **Degrees of Freedom** | **P value** |
| Species | 451.02 | 7 | <2.2E-16 |
| Social group | 62.14 | 2 | 3.21E-14 |
| Species: social group | 71.64 | 14 | 9.71E-10 |

| Term | Estimate | Degrees of freedom | T-value |
|---|---|---|---|
| Intercept | 4.4 | 997 | 26.99 |
| Species Groove-billed Ani (GBAN) | -0.07 | 997 | -0.41 |
| Species Great-tailed Grackle (GTGR) | -0.55 | 997 | -2.57 |
| Species Keel-billed Toucan (KBTO) | 0.28 | 997 | 1.27 |
| Species Long-tailed Manakin (LTMA) | 0.46 | 997 | 2.69 |
| Species Rufous-naped Wren (RNWR) | 0.08 | 997 | 0.39 |
| Species White-throated Magpie-Jay (WTMJ) | 0.17 | 997 | 0.98 |
| Social group Farmers | 0.04 | 396 | 0.19 |
| Social group Urbanites | -0.17 | 396 | -0.8 |
| GBAN: farmers | -0.86 | 997 | -3.73 |
| GTGR: farmers | -1.3 | 997 | -4.42 |
| KBTO: farmers | -0.37 | 997 | -1.27 |
| LTMA: farmers | -0.46 | 997 | -1.95 |
| OCPA: farmers | -0.5 | 997 | -1.7 |
| RNWR: farmers | -0.74 | 997 | -2.48 |
| WTMJ: farmers | -1.06 | 997 | -4.7 |
| GBAN: urbanites | -0.96 | 997 | -4.08 |
| GTGR: urbanites | -1.5 | 997 | -4.96 |
| KBTO: urbanites | -0.35 | 997 | -1.19 |
| LTMA: urbanites | -0.22 | 997 | -0.91 |
| OCPA: urbanites | -0.74 | 997 | -2.53 |
| RNWR: urbanites | -0.9 | 997 | -3.01 |
| WTMJ: urbanites | -0.74 | 997 | -3.21 |

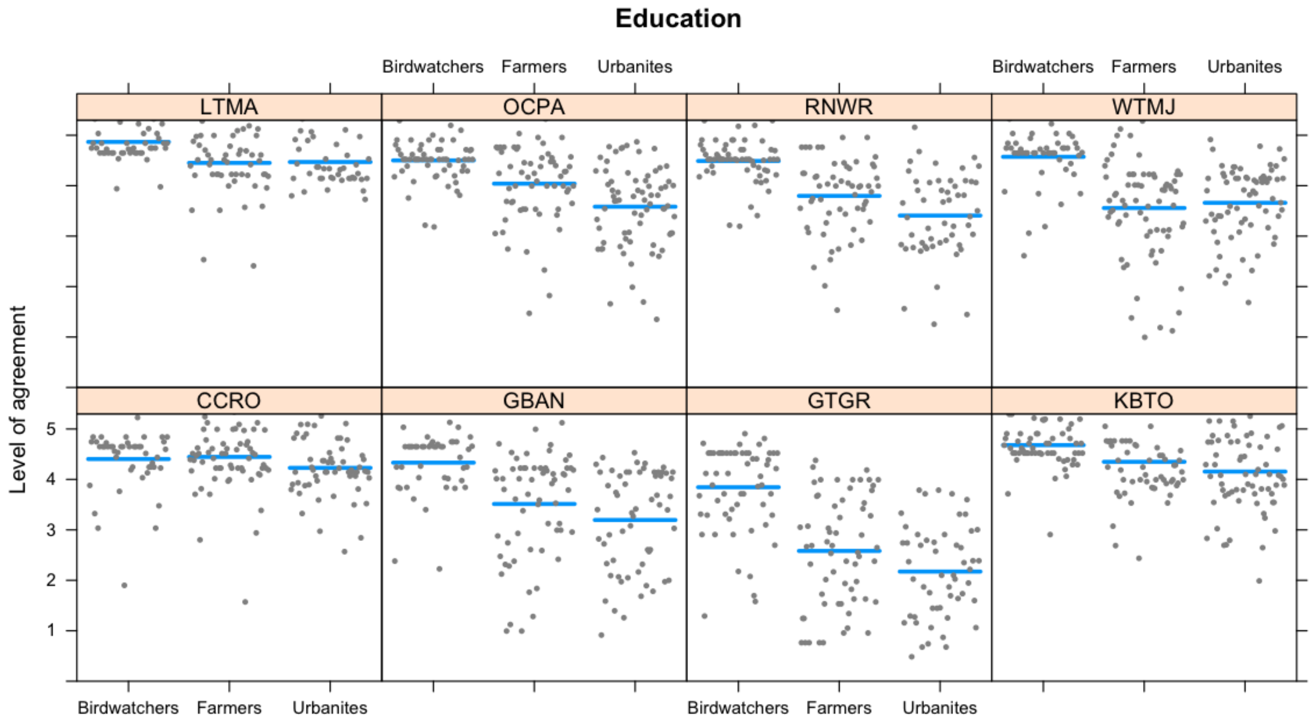

**Figure S2. Model fits for each species and three social groups regarding education scores.** Each panel represents a species (LTMA=Long-tailed Manakin, OCPA=Orange-chinned Parakeet, RNWR=Rufous-naped Wren, WTMJ=White-throated Magpie-Jay, CCRO=Clay-colored Thrush, GBAN=Groove-billed Ani, GTGR=Great-tailed Grackle, KBTO=Keel-billed Toucan). Each grey dot represents one person, and the blue lines represent the model.

**Table S4. Tukey HSD pairwise comparisons for education scores.**

| **Species** | **Pairwise comparison** | **P value** |
|---|---|---|
| Great-tailed Grackle | Birdwatchers vs. farmers | $p$<0.00001*** |
| | Birdwatchers vs. urbanites | $p$<0.00001*** |
| | Farmers vs. urbanites | $p$>0.05 |
| Orange-chinned Parakeet | Birdwatchers vs. farmers | $p$=0.044* |
| | Birdwatchers vs. urbanites | $p$<0.00001*** |
| | Farmers vs. urbanites | $p$>0.05 |
| White-throated Magpie-Jay | Birdwatchers vs. farmers | $p$<0.0001*** |
| | Birdwatchers vs. urbanites | $p$<0.001*** |
| | Farmers vs. urbanites | $p$>0.05 |
| Rufous-naped Wren | Birdwatchers vs. farmers | $p$=0.002** |
| | Birdwatchers vs. urbanites | $p$<0.001*** |
| | Farmers vs. urbanites | $p$>0.05 |
| Groove-billed Ani | Birdwatchers vs. urbanites | $p$<0.001 |
| | Birdwatchers vs. farmers | $p$=0.011* |
| | Farmers vs. urbanites | $p$>0.05 |
| Clay-colored Thrush | Birdwatchers vs. urbanites | $p$>0.05 |
| | Birdwatchers vs. farmers | $p$>0.05 |

| Species | Pairwise comparison | P value |
|---|---|---|
| | Farmers vs. urbanites | $p$>0.05 |
| Keel-billed Toucan | Birdwatchers vs. urbanites | $p$=0.006** |
| | Birdwatchers vs. farmers | $p$>0.05 |
| | Farmers vs. urbanites | $p$>0.05 |
| Long-tailed Manakin | Birdwatchers vs. urbanites | $p$>0.05 |
| | Birdwatchers vs. farmers | $p$>0.05 |
| | Farmers vs. urbanites | $p$>0.05 |

*p<0.05, *p<0.01, ***p<0.001

#### 3.3 BIRDWATCHING: MIXED-EFFECTS MODEL

| **Model** | **Birdwatching = Species + Social group + Species*Social group + (1\|Participant)** |
|---|---|
| AIC= 3973.30; BIC=4109.52 | |

| **Type II Anova** | | | |
|---|---|---|---|
| **Term** | **Chi-squared** | **Degrees of Freedom** | **P value** |
| Species | 1214.81 | 7 | <2.26E-16 |
| Social group | 29.26 | 2 | 4.43 E-07 |
| Species: social group | 111.02 | 14 | <2.2E-16 |

| **Term** | **Estimate** | **Degrees of freedom** | **T-value** |
|---|---|---|---|
| Intercept | 4.13 | 997 | 29.17 |
| Species Groove-billed Ani (GBAN) | -0.5 | 997 | -2.8 |
| Species Great-tailed Grackle (GTGR) | -0.84 | 997 | -4.43 |
| Species Keel-billed Toucan (KBTO) | 0.68 | 997 | 3.62 |
| Species Long-tailed Manakin (LTMA) | 0.87 | 997 | 4.91 |
| Species Orange-chinned Parakeet (OCPA) | 0.47 | 997 | 2.48 |
| Species Rufous-naped Wren (RNWR) | 0.39 | 997 | 2.09 |
| Species White-throated Magpie-Jay (WTMJ) | 0.5 | 997 | 2.86 |
| Social group Farmers | 0.56 | 396 | 2.99 |
| Social group Urbanites | 0.43 | 396 | 2.23 |
| GBAN: farmers | -1.26 | 997 | -5.31 |
| GTGR: farmers | -1.61 | 997 | -6.26 |
| KBTO: farmers | -0.43 | 997 | -1.69 |
| LTMA: farmers | -0.62 | 997 | -2.59 |
| OCPA: farmers | -0.68 | 997 | -2.66 |
| RNWR: farmers | -0.9 | 997 | -3.47 |
| WTMJ: farmers | -1.2 | 997 | -5.19 |
| GBAN: urbanites | -1.22 | 997 | -5.07 |
| GTGR: urbanites | -2 | 997 | -7.6 |
| KBTO: urbanites | -0.4 | 997 | -1.56 |
| LTMA: urbanites | -0.55 | 997 | -2.19 |
| OCPA: urbanites | -0.67 | 997 | -2.63 |
| RNWR: urbanites | -0.96 | 997 | -3.63 |
| WTMJ: urbanites | -1.14 | 997 | -4.88 |

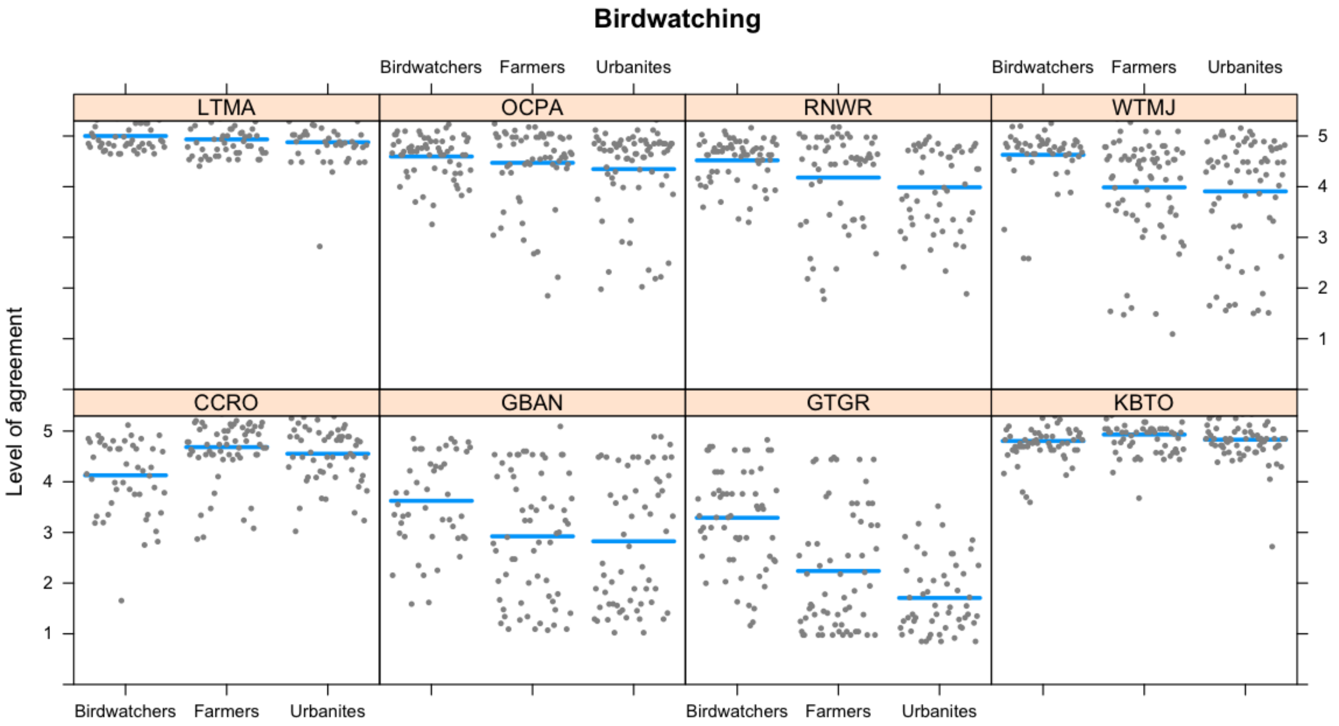

**Figure S3. Model fits for each species and three social groups regarding birdwatching scores.** Each panel represents a species (LTMA=Long-tailed Manakin, OCPA=Orange-chinned Parakeet, RNWR=Rufous-naped Wren, WTMJ=White-throated Magpie-Jay, CCRO=Clay-colored Thrush, GBAN=Groove-billed Ani, GTGR=Great-tailed Grackle, KBTO=Keel-billed Toucan). Each grey dot represents one person, and the blue lines represent the model.

**Table S5. Tukey HSD pairwise comparisons for birdwatching scores.**

| Species | Pairwise comparison | P value |
|---|---|---|
| Great-tailed Grackle | Birdwatchers vs. farmers | $p<0.0001$*** |
| | Birdwatchers vs. urbanites | $p<0.0001$*** |
| | Farmers vs. urbanites | $p>0.05$ |
| Orange-chinned Parakeet | Birdwatchers vs. farmers | $p>0.05$ |
| | Birdwatchers vs. urbanites | $p>0.05$ |
| | Farmers vs. urbanites | $p>0.05$ |
| White-throated Magpie-Jay | Birdwatchers vs. farmers | $p=0.008$** |
| | Birdwatchers vs. urbanites | $p=0.003$** |
| | Farmers vs. urbanites | $p>0.05$ |
| Rufous-naped Wren | Birdwatchers vs. farmers | $p<0.001$*** |
| | Birdwatchers vs. urbanites | $p<0.001$*** |
| | Farmers vs. urbanites | $p>0.05$ |
| Groove-billed Ani | Birdwatchers vs. farmers | $p<0.001$*** |
| | Birdwatchers vs. urbanites | $p<0.001$*** |
| | Farmers vs. urbanites | $p>0.05$ |
| Clay-colored Thrush | Birdwatchers vs. farmers | $p<0.0001$*** |
| | Birdwatchers vs. urbanites | $p=0.002$** |

| Species | Pairwise comparison | P value |
|---|---|---|
| | Farmers vs. urbanites | $p$>0.05 |
| Keel-billed Toucan | Birdwatchers vs. farmers | $p$>0.05 |
| | Birdwatchers vs. urbanites | $p$>0.05 |
| | Farmers vs. urbanites | $p$>0.05 |
| Long-tailed Manakin | Birdwatchers vs. farmers | $p$>0.05 |
| | Birdwatchers vs. urbanites | $p$>0.05 |
| | Farmers vs. urbanites | $p$>0.05 |

*p<0.05, *p<0.01, ***p<0.001

#### 3.4 ACOUSTIC AESTHETICS: MULTINOMIAL REGRESSION

**Model** **Multinomial (Acoustic aesthetics = Species + Social group + Species*Social group)**

| Type II Anova | | | |
|---|---|---|---|
| **Term** | **Chi-squared** | **Degrees of Freedom** | **P value** |
| Species | 469.74 | 28 | <2.2 E-16 |
| Social group | 47.88 | 8 | 1.04E-07 |
| Species: social group | 82.2 | 56 | 0.013 |

| Probabilities predicted by the multinomial model | | | | | | |
|---|---|---|---|---|---|---|
| **Species** | **Social Group** | **1 (strongly disagree)** | **2 (disagree)** | **3 (neutral)** | **4 (agree)** | **5 (strongly agree)** |
| Clay-colored Thrush | Birdwatchers | 0.00 | 0.02 | 0.02 | 0.10 | 0.85 |
| Groove-billed Ani | Birdwatchers | 0.17 | 0.15 | 0.26 | 0.22 | 0.20 |
| Great-tailed Grackle | Birdwatchers | 0.13 | 0.10 | 0.23 | 0.21 | 0.33 |
| Keel-billed Toucan | Birdwatchers | 0.08 | 0.17 | 0.16 | 0.17 | 0.42 |
| Long-tailed Manakin | Birdwatchers | 0.00 | 0.00 | 0.11 | 0.13 | 0.77 |
| Orange-chinned Parakeet | Birdwatchers | 0.08 | 0.15 | 0.26 | 0.15 | 0.37 |
| Rufous-naped Wren | Birdwatchers | 0.00 | 0.02 | 0.08 | 0.19 | 0.71 |
| White-throated Magpie-Jay | Birdwatchers | 0.27 | 0.20 | 0.16 | 0.27 | 0.10 |
| Clay-colored Thrush | Farmers | 0.00 | 0.00 | 0.00 | 0.00 | 1.00 |
| Groove-billed Ani | Farmers | 0.19 | 0.11 | 0.13 | 0.24 | 0.33 |
| Great-tailed Grackle | Farmers | 0.32 | 0.16 | 0.05 | 0.15 | 0.32 |
| Keel-billed Toucan | Farmers | 0.05 | 0.07 | 0.08 | 0.25 | 0.56 |
| Long-tailed Manakin | Farmers | 0.00 | 0.02 | 0.00 | 0.02 | 0.96 |
| Orange-chinned Parakeet | Farmers | 0.06 | 0.14 | 0.05 | 0.21 | 0.54 |
| Rufous-naped Wren | Farmers | 0.03 | 0.05 | 0.09 | 0.09 | 0.74 |
| White-throated Magpie-Jay | Farmers | 0.35 | 0.07 | 0.10 | 0.22 | 0.26 |
| Clay-colored Thrush | Urbanites | 0.02 | 0.00 | 0.02 | 0.02 | 0.95 |
| Groove-billed Ani | Urbanites | 0.25 | 0.07 | 0.10 | 0.20 | 0.38 |
| Great-tailed Grackle | Urbanites | 0.38 | 0.17 | 0.15 | 0.09 | 0.21 |
| Keel-billed Toucan | Urbanites | 0.13 | 0.09 | 0.13 | 0.14 | 0.51 |
| Long-tailed Manakin | Urbanites | 0.00 | 0.00 | 0.00 | 0.02 | 0.98 |
| Orange-chinned Parakeet | Urbanites | 0.10 | 0.15 | 0.12 | 0.11 | 0.51 |
| Rufous-naped Wren | Urbanites | 0.09 | 0.06 | 0.02 | 0.19 | 0.64 |
| White-throated Magpie-Jay | Urbanites | 0.23 | 0.20 | 0.12 | 0.13 | 0.32 |

**Table S6. Tukey HSD pairwise comparisons for acoustic aesthetics scores.**

| Species | Pairwise comparison | P value |
|---|---|---|
| Great-tailed Grackle | Birdwatchers vs. farmers | $p$>0.05 |
| | Birdwatchers vs. urbanites | $p$=0.005** |
| | Farmers vs. urbanites | $p$>0.05 |
| Orange-chinned Parakeet | Birdwatchers vs. farmers | $p$>0.05 |
| | Birdwatchers vs. urbanites | $p$>0.05 |
| | Farmers vs. urbanites | $p$>0.05 |
| White-throated Magpie-Jay | Birdwatchers vs. farmers | $p$>0.05 |
| | Birdwatchers vs. urbanites | $p$>0.05 |
| | Farmers vs. urbanites | $p$>0.05 |
| Rufous-naped Wren | Birdwatchers vs. farmers | $p$>0.05 |
| | Birdwatchers vs. urbanites | $p$>0.05 |

| Species | Pairwise comparison | P value |
|---|---|---|
| | Farmers vs. urbanites | *p*>0.05 |
| Groove-billed Ani | Birdwatchers vs. farmers | *p*>0.05 |
| | Birdwatchers vs. urbanites | *p*>0.05 |
| | Farmers vs. urbanites | *p*>0.05 |
| Clay-colored Thrush | Birdwatchers vs. farmers | *p*>0.05 |
| | Birdwatchers vs. urbanites | *p*>0.05 |
| | Farmers vs. urbanites | *p*>0.05 |
| Keel-billed Toucan | Birdwatchers vs. farmers | *p*>0.05 |
| | Birdwatchers vs. urbanites | *p*>0.05 |
| | Farmers vs. urbanites | *p*>0.05 |
| Long-tailed Manakin | Birdwatchers vs. farmers | *p*=0.014* |
| | Birdwatchers vs. urbanites | *p*=0.005** |
| | Farmers vs. urbanites | *p*>0.05 |

*p<0.05, *p<0.01, ***p<0.001

#### 3.5 IDENTITY: MIXED-EFFECTS MODEL

**Model** **Identity = Species + Social group + Species*Social group + (1|Participant)**

AIC= 4091.24; BIC=4227.46

| Type II Anova | | | |
|---|---|---|---|
| **Term** | **Chi-squared** | **Degrees of Freedom** | **P value** |
| Species | 262.2 | 7 | <2.26E-16 |
| Social group | 15.32 | 2 | 4.00E-04 |
| Species: social group | 32.39 | 14 | 3.50E-04 |

| **Term** | **Estimate** | **Degrees of freedom** | **T-value** |
|---|---|---|---|
| Intercept | 4.36 | 997 | 29.14 |
| Species Groove-billed Ani (GBAN) | -0.44 | 997 | -2.37 |
| Species Great-tailed Grackle (GTGR) | -0.99 | 997 | -5 |
| Species Keel-billed Toucan (KBTO) | -0.44 | 997 | -2.23 |
| Species Long-tailed Manakin (LTMA) | -0.19 | 997 | -1.05 |
| Species Orange-chinned Parakeet (OCPA) | -0.21 | 997 | -1.09 |
| Species Rufous-naped Wren (RNWR) | -0.19 | 997 | -0.96 |
| Species White-throated Magpie-Jay (WTMJ) | -0.38 | 997 | -2.11 |
| Social group Farmers | 0.39 | 396 | 2.02 |
| Social group Urbanites | 0.19 | 396 | 0.97 |
| GBAN: farmers | -0.18 | 997 | -0.75 |
| GTGR: farmers | -0.32 | 997 | -1.17 |
| KBTO: farmers | -0.75 | 997 | -2.77 |
| LTMA: farmers | -0.23 | 997 | -0.92 |
| OCPA: farmers | 0.11 | 997 | 0.4 |
| RNWR: farmers | -0.01 | 997 | -0.05 |
| WTMJ: farmers | -0.08 | 997 | -0.32 |
| GBAN: urbanites | -0.38 | 997 | -1.56 |
| GTGR: urbanites | -0.84 | 997 | -3.02 |
| KBTO: urbanites | -0.43 | 997 | -1.6 |
| LTMA: urbanites | -0.3 | 997 | -1.17 |
| OCPA: urbanites | -0.1 | 997 | -0.36 |
| RNWR: urbanites | -0.29 | 997 | -1.04 |
| WTMJ: urbanites | -0.24 | 997 | -0.97 |

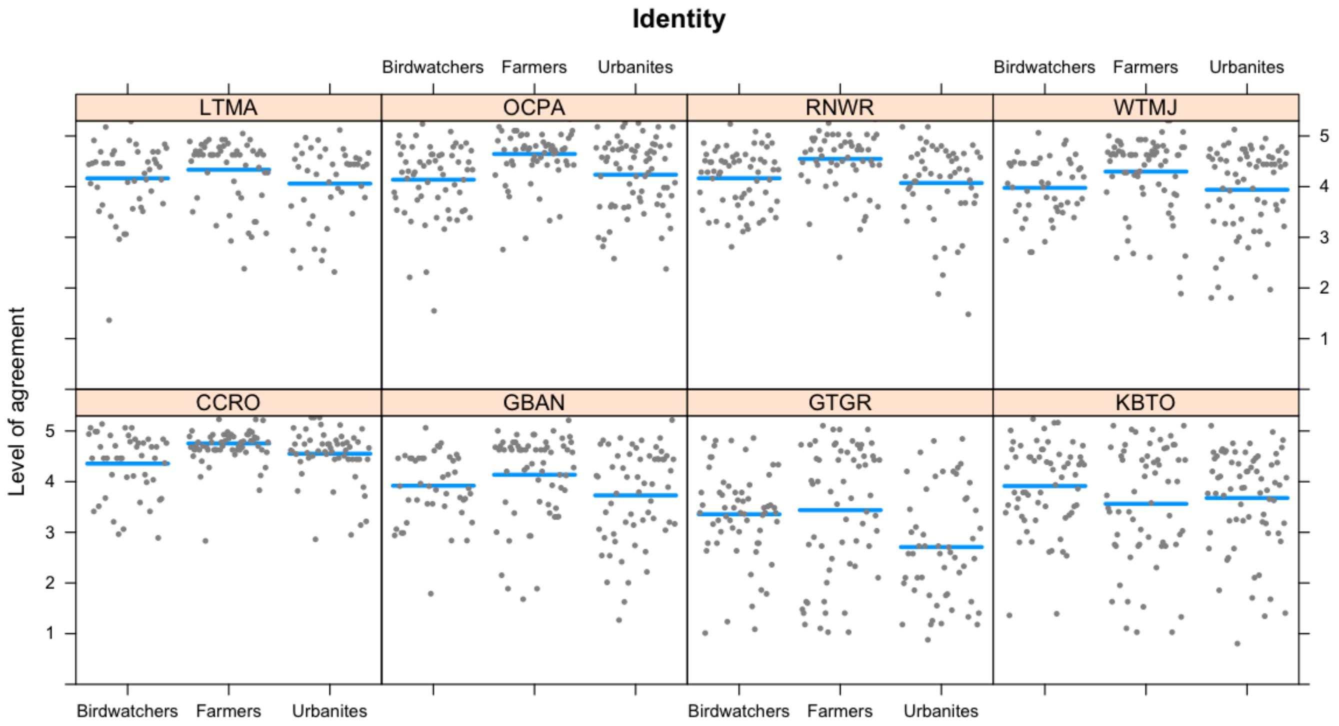

**Figure S4. Model fits for each species and three social groups regarding identity scores.** Each panel represents a species (LTMA=Long-tailed Manakin, OCPA=Orange-chinned Parakeet, RNWR=Rufous-naped Wren, WTMJ=White-throated Magpie-Jay, CCRO=Clay-colored Thrush, GBAN=Groove-billed Ani, GTGR=Great-tailed Grackle, KBTO=Keel-billed Toucan). Each grey dot represents one person, and the blue lines represent the model.

**Table S7. Tukey HSD pairwise comparisons for identity scores.**

| **Species** | **Pairwise comparison** | **P value** |
|---|---|---|
| Great-tailed Grackle | Birdwatchers vs. farmers | $p>0.05$ |
| | Birdwatchers vs. urbanites | $p=0.018$* |
| | Farmers vs. urbanites | $p=0.022$* |
| Orange-chinned Parakeet | Birdwatchers vs. farmers | $p=0.004$** |
| | Birdwatchers vs. urbanites | $p>0.05$ |
| | Farmers vs. urbanites | $p>0.05$ |
| White-throated Magpie-Jay | Birdwatchers vs. farmers | $p>0.05$ |
| | Birdwatchers vs. urbanites | $p>0.05$ |
| | Farmers vs. urbanites | $p>0.05$ |
| Rufous-naped Wren | Birdwatchers vs. farmers | $p>0.05$ |
| | Birdwatchers vs. urbanites | $p>0.05$ |
| | Farmers vs. urbanites | $p=0.02$* |
| Groove-billed Ani | Birdwatchers vs. farmers | $p>0.05$ |
| | Birdwatchers vs. urbanites | $p>0.05$ |
| | Farmers vs. urbanites | $p>0.05$ |
| Clay-colored Thrush | Birdwatchers vs. farmers | $p=0.003$** |
| | Birdwatchers vs. urbanites | $p>0.05$ |

| Species | Pairwise comparison | P value |
|---|---|---|
| | Farmers vs. urbanites | $p>0.05$ |
| Keel-billed Toucan | Birdwatchers vs. farmers | $p>0.05$ |
| | Birdwatchers vs. urbanites | $p>0.05$ |
| | Farmers vs. urbanites | $p>0.05$ |
| Long-tailed Manakin | Birdwatchers vs. farmers | $p>0.05$ |
| | Birdwatchers vs. urbanites | $p>0.05$ |
| | Farmers vs. urbanites | $p>0.05$ |

*p<0.05, *p<0.01, ***p<0.001

#### 3.6 BEQUEST: LOGISTIC REGRESSION

| | |
|---|---|
| | **Bequest (as a binary variable) ~ Species + Social group + Species*Social group** |
| **Model** | **Type of family= binomial (logit function)** |
| AIC= 957.29 | |

| **Type II Anova** | | | |
|---|---|---|---|
| **Term** | **Chi-squared** | **Degrees of Freedom** | **P value** |
| Species | 260.27 | 7 | <2.26E-16 |
| Social group | 32.28 | 2 | 9.78E-08 |
| Species: social group | 17.99 | 14 | 0.21 |

| **Term** | **Estimate** | **z-value** |
|---|---|---|
| Intercept | 3.2 | 4.43 |
| Species Groove-billed Ani (GBAN) | -0.98 | -1.14 |
| Species Great-tailed Grackle (GTGR) | -1.81 | -2.31 |
| Species Keel-billed Toucan (KBTO) | 0.25 | 0.25 |
| Species Long-tailed Manakin (LTMA) | 0.71 | 0.57 |
| Species Orange-chinned Parakeet (OCPA) | 0.25 | 0.25 |
| Species Rufous-naped Wren (RNWR) | -0.17 | -0.18 |
| Species White-throated Magpie-Jay (WTMJ) | -0.43 | -0.46 |
| Social group Farmers | 1.12 | 0.9 |
| Social group Urbanites | 1.12 | 0.9 |
| GBAN: farmers | -1.66 | -1.22 |
| GTGR: farmers | -2.88 | -2.22 |
| KBTO: farmers | -0.43 | -0.24 |
| LTMA: farmers | -0.71 | -0.34 |
| OCPA: farmers | -2.76 | -1.87 |
| RNWR: farmers | -1.68 | -1.16 |
| WTMJ: farmers | -2 | -1.42 |
| GBAN: urbanites | -2.17 | -1.6 |
| GTGR: urbanites | -3.02 | -2.32 |
| KBTO: urbanites | -0.31 | -0.18 |
| LTMA: urbanites | -1.42 | -0.81 |
| OCPA: urbanites | -1.97 | -1.31 |
| RNWR: urbanites | -2.43 | -1.73 |
| WTMJ: urbanites | -1.88 | -1.3 |

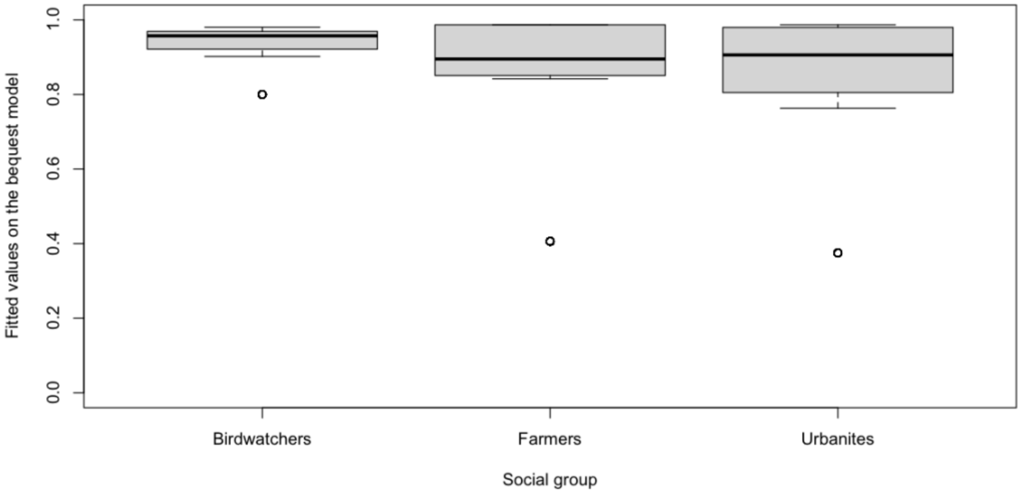

**Figure S5. Visualizing predicted values from the model by pooling all eight focal species and just comparing across social groups.**

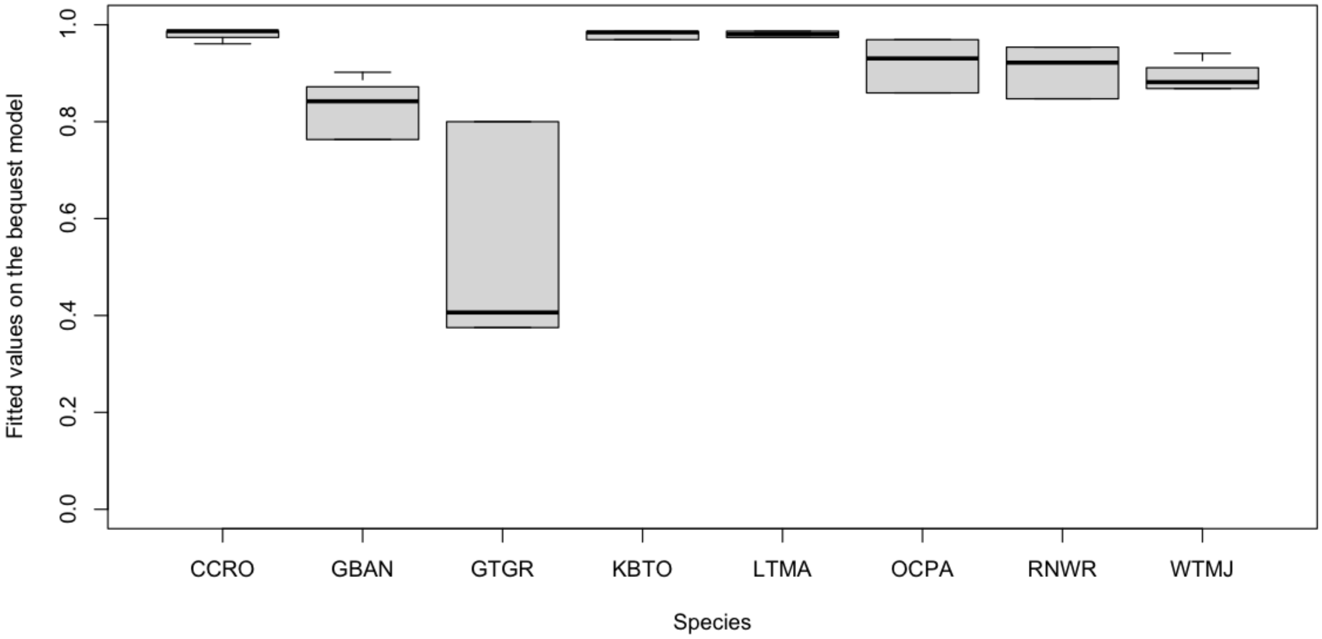

**Figure S6. Visualizing predicted values from the model by pooling all social groups and just comparing across species** (CCRO=Clay-colored Thrush, GBAN=Groove-billed Ani, GTGR=Great-tailed Grackle, KBTO=Keel-billed Toucan, LTMA=Long-tailed Manakin, OCPA=Orange-chinned Parakeet, RNWR=Rufous-naped Wren, WTMJ=White-throated Magpie-Jay).

**Table S8. Tukey HSD pairwise comparisons for bequest scores.**

| **Species** | **Pairwise comparison** | ***p* value** |
|---|---|---|
| Great-tailed Grackle | Birdwatchers vs. farmers | *p*<0.00001*** |
| | Birdwatchers vs. urbanites | *p*<0.00001*** |
| | Farmers vs. urbanites | *p*>0.05 |
| Orange-chinned Parakeet | Birdwatchers vs. farmers | *p*>0.05 |
| | Birdwatchers vs. urbanites | *p*>0.05 |
| | Farmers vs. urbanites | *p*>0.05 |
| White-throated Magpie-Jay | Birdwatchers vs. farmers | *p*>0.05 |
| | Birdwatchers vs. urbanites | *p*>0.05 |
| | Farmers vs. urbanites | *p*>0.05 |
| Rufous-naped Wren | Birdwatchers vs. farmers | *p*>0.05 |
| | Birdwatchers vs. urbanites | *p*>0.05 |
| | Farmers vs. urbanites | *p*>0.05 |
| Groove-billed Ani | Birdwatchers vs. farmers | *p*>0.05 |
| | Birdwatchers vs. urbanites | *p*>0.05 |
| | Farmers vs. urbanites | *p*<0.05* |
| Clay-colored Thrush | Birdwatchers vs. farmers | *p*>0.05 |
| | Birdwatchers vs. urbanites | *p*>0.05 |
| | Farmers vs. urbanites | *p*>0.05 |
| Keel-billed Toucan | Birdwatchers vs. farmers | *p*>0.05 |
| | Birdwatchers vs. urbanites | *p*>0.05 |
| | Farmers vs. urbanites | *p*>0.05 |
| Long-tailed Manakin | Birdwatchers vs. farmers | *p*>0.05 |
| | Birdwatchers vs. urbanites | *p*>0.05 |
| | Farmers vs. urbanites | *p*>0.05 |

*p<0.05, *p<0.01, ***p<0.001

### 4 RESULTS FROM THE OPEN-ENDED QUESTIONS

#### 4.1 DISSERVICES

**Table S9. List of species or groups with the associated mentions by birdwatchers, farmers, and urbanites.** Frequencies are reported in terms of how many times each species/group was mentioned when participants were asked to say which species they disliked or found annoying or harmful.

| Species or groups | Birdwatchers | Farmers | Urbanites |
|---|---|---|---|
| None | 43 | 21 | 17 |
| Great-tailed Grackle | 41 | 91 | 89 |
| Vultures | 11 | 16 | 37 |
| Doves | 8 | 9 | 7 |
| Cowbirds | 5 | | 1 |
| White-throated Magpie-Jay | 4 | 27 | 16 |
| Tricolored Munia | 4 | | |
| Rufous-naped Wren | 3 | 2 | 9 |
| House Sparrow | 3 | | |
| Invasive species | 2 | | |
| Groove-billed Ani | 1 | 14 | 11 |
| Parrots | 1 | 3 | 4 |
| Chachalaca | 1 | 3 | 1 |
| Black birds | 1 | 1 | |
| Toucan | 1 | | 1 |
| Blue-headed Parrot | 1 | | |
| Thrushes | 1 | | |
| Southern Lapwing | 1 | | |
| Warblers | 1 | | |
| Parakeets | | 28 | 16 |
| Woodpeckers | | 12 | 1 |
| Fulvous Whistling-Duck | | 9 | |
| Hawks | | 7 | 4 |
| Quails | | 2 | 1 |
| Crested Caracara | | 2 | |
| Melodious Blackbird | | 2 | |
| Anhinga | | 1 | |
| Oropendola | | 1 | |
| Potoos | | 1 | |
| Tytyras | | 1 | |
| Black guan | | 1 | |
| Purple Gallinule | | 1 | |
| Laughing Falcon | | 1 | |
| Tyrant Flycatcher | | 1 | |
| Magnificent Frigate bird | | 1 | |
| Peacocks | | | 3 |
| American Crow | | | 3 |

| Species or groups | Birdwatchers | Farmers | Urbanites |
|---|---|---|---|
| Barn Owl | | | 2 |
| Owls | | | 1 |
| Boat-billed Heron | | | 1 |
| Ostriches | | | 1 |

#### 4.2 BIRDWATCHING AND ACOUSTIC AESTHETICS

**Table S10. List of species or groups with the associated mentions by birdwatchers, farmers, and urbanites.** Frequencies are reported in terms of how many times each species/group was mentioned when participants were asked to say which species they enjoyed watching or hearing.

| Species or group | Birdwatchers | Farmers | Urbanites |
|---|---|---|---|
| Clay-colored Thrush | 1 | 58 | 45 |
| Toucans | 7 | 39 | 42 |
| Parakeets | 3 | 31 | 27 |
| Long-tailed Manakin | 27 | 28 | 15 |
| Hummingbirds | 13 | 25 | 32 |
| White-throated Magpie-Jay | 12 | 25 | 42 |
| Scarlett Macaw | 7 | 25 | 40 |
| Parrots | 4 | 24 | 20 |
| Chachalacas | 2 | 20 | 3 |
| Flycatchers with yellow breasts | 1 | 20 | 26 |
| Doves | 1 | 16 | 11 |
| Jabiru | 26 | 12 | 7 |
| Momots | 16 | 12 | 18 |
| Orioles | 7 | 11 | 8 |
| Blue-gray Tanager | 1 | 10 | 7 |
| Rufuous-naped Wren | 3 | 8 | 11 |
| Fulvous Whistling Duck | 1 | 7 | 4 |
| Crested Bobwhite | 1 | 7 | 3 |
| Roseate Spoonbill | 7 | 6 | 4 |
| All of them | 6 | 6 | 3 |
| Herons | 5 | 6 | 2 |
| Summer Tanager | | 6 | |
| Euphonia | 6 | 5 | |
| Hawks | 4 | 5 | 4 |
| Woodpeckers | 3 | 5 | 12 |
| Canaries | | 5 | 8 |
| Trogons | 10 | 4 | 8 |
| Ducks | 5 | 4 | 2 |
| Double-striped Thick-knee | 5 | 4 | |
| Red-winged Blackbird | | 4 | 7 |
| Great-tailed Grackle | | 4 | 5 |
| Brown doves | | 4 | 1 |
| Domestic chickens | | 4 | |

| Species or group | Birdwatchers | Farmers | Urbanites |
|---|---|---|---|
| Thicket tinamou | 6 | 3 | 2 |
| Great Currassow | 4 | 3 | 1 |
| Seed finches | 2 | 3 | |
| Great Green Macaw | | 3 | 2 |
| Peacock | | 3 | 2 |
| Vultures | | 3 | 2 |
| White-tipped Dove | | 3 | 1 |
| Laughing Falcon | 5 | 2 | 7 |
| Three-wattled Bell bird | 4 | 2 | 1 |
| Yellow-naped Parrot | 4 | 2 | |
| Oropendola | 3 | 2 | 3 |
| Tiger Heron | 1 | 2 | 2 |
| Swallows | | 2 | 3 |
| Raptors | 14 | 1 | 2 |
| King fishers | 4 | 1 | 3 |
| Scissor-tailed Flycatcher | 4 | 1 | |
| Crested Caracara | 3 | 1 | 4 |
| Owls | 3 | 1 | 2 |
| White Ibis | 3 | 1 | |
| Sea gulls | 2 | 1 | 1 |
| Great Egret | 1 | 1 | 2 |
| Violaceous Quail-Dove | 1 | 1 | |
| Common Pauraque | | 1 | 6 |
| Southern Lapwing | | 1 | 1 |
| Melodious black bird | | 1 | |
| Anhiga | | 1 | |
| Black Guan | | 1 | |
| Keel-billed Toucan | | 1 | |
| Tanagers | 10 | | 1 |
| Tody Motmot | 7 | | |
| Wrens | 6 | | 3 |
| Lesser Ground Cuckoo | 6 | | |
| Elegant Trogon | 5 | | 2 |
| Black-headed Trogon | 5 | | |
| Resplandescent Quetzal | 4 | | 5 |
| Barn Owl | 4 | | 3 |
| Shorebirds | 4 | | 1 |
| Manakins | 4 | | |
| Ornate Hawk Eagle | 4 | | |
| Water birds | 3 | | **3** |
| Boat-billed heron | 3 | | 1 |
| Painted Bunting | 3 | | 1 |
| Warblers | 3 | | 1 |
| Mangrove Hummingbird | 3 | | |

| Species or group | Birdwatchers | Farmers | Urbanites |
|---|---|---|---|
| Banded Wren | 3 | | |
| Ferruginous Pygmy Owl | 3 | | |
| Potoo | 3 | | |
| Bare-necked Umbrella Bird | 3 | | |
| Pearl Kite | 3 | | |
| Spot-breasted Oriole | 3 | | |
| White fronted Parrot | 3 | | |
| Rufous-naped Wood rail | 2 | | 3 |
| Migratory birds | 2 | | 1 |
| Collared Araçari | 2 | | 1 |
| Striped Cuckoo | 2 | | 1 |
| Chestnut-bellied Heron | 2 | | |
| Cinnamon Hummingbird | 2 | | |
| Birds endemic to Guanacaste | 2 | | |
| Marine birds | 2 | | |
| Cotingas | 2 | | |
| Cattle Egret | 2 | | |
| Tricolor Heron | 2 | | |
| Ant birds | 2 | | |
| King Vulture | 2 | | |
| Panama Flycatcher | 2 | | |
| Blue Groskbeak | 2 | | |
| Rufous-vented Ground-cuckoo | 2 | | |
| Rufous-capped Warbler | 2 | | |
| Yellow Warbler | 2 | | |
| Streak-backed Oriole | 2 | | |
| Mangrove Vireo | 2 | | |
| White-necked Puffbird | 2 | | |
| Peregrine Falcon | 1 | | 2 |
| Pacific Screech-Owl | 1 | | 1 |
| Brown Pelican | 1 | | 1 |
| Prothonotary Warbler | 1 | | 1 |
| Yellow-eared Toucanet | 1 | | 1 |
| American Oystercatcher | 1 | | |
| Endangered birds | 1 | | |
| Nocturnal birds | 1 | | |
| Resident birds | 1 | | |
| Mixed-species flocks birds | 1 | | |
| Barred Antshrike | 1 | | |
| Bat Falcon | 1 | | |
| Black-faced solitaire | 1 | | |
| Blue-black grassquit | 1 | | |
| Botteri's Sparrow | 1 | | |
| Brown-crested Flycatcher | 1 | | |

| Species or group | Birdwatchers | Farmers | Urbanites |
|---|---|---|---|
| Canivet's Emerald | 1 | | |
| Orange-billed Nightingale Thrush | 1 | | |
| Cinnamon Beccard | 1 | | |
| Crane Hawk | 1 | | |
| Red-legged Honeycreeper | 1 | | |
| Fiery-throated Hummingbird | 1 | | |
| Little Blue Heron | 1 | | |
| Snowy Egret | 1 | | |
| Golden-hooded Tanager | 1 | | |
| Gray-crowned Yellowthroat | 1 | | |
| Green-breasted Mango | 1 | | |
| Grey hawk | 1 | | |
| Hook-billed Kite | 1 | | |
| Glossy Ibis | 1 | | |
| Baltimore's Oriole | 1 | | |
| Little Tinamou | 1 | | |
| Lesser Swallow-tailed Swift | 1 | | |
| Mangrove Cuckoo | 1 | | |
| Collared Forest Falcon | 1 | | |
| Ochre-bellied Flycatcher | 1 | | |
| Montezuma Oropendola | 1 | | |
| Northern Beardless-Tyrannulet | 1 | | |
| Northern Scrub-Flycatcher | 1 | | |
| Olive Sparrow | 1 | | |
| Orange-fronted Parakeet | 1 | | |
| Osprey | 1 | | |
| Lineated Woodpecker | 1 | | |
| White doves | 1 | | |
| Western Tanager | 1 | | |
| Plain-brown Woodcreeper | 1 | | |
| Plain-breasted Ground Dove | 1 | | |
| Plumbeous Kite | 1 | | |
| Rock Wren | 1 | | |
| Royal Tern | 1 | | |
| Ruby-throated Hummingbird | 1 | | |
| Ruddy Turnstone | 1 | | |
| Rufous-and-white Wren | 1 | | |
| Rufous-browed Peppershrike | 1 | | |
| Rusty Sparrow | 1 | | |
| Sandwich Tern | 1 | | |
| Scaly-breasted Hummingbird | 1 | | |
| Sharpbill | 1 | | |
| Short-tailed Hawk | 1 | | |
| Slate-throated Redstarts | 1 | | |

| Species or group | Birdwatchers | Farmers | Urbanites |
|---|---|---|---|
| Steely-vented Hummingbird | 1 | | |
| Stripe-headed Sparrow | 1 | | |
| Boobies | 1 | | |
| Chesnut-mandibled Toucan or Swainson's Toucan | 1 | | |
| White-collared Manakin | 1 | | |
| White-lored Gnatcatcher | 1 | | |
| White-tailed Nightjar | 1 | | |
| Yellow-headed Caracara | 1 | | |
| Dickcissels | | | 1 |
| Grebes | | | 1 |
| Flycatchers | | | |
| Burrowing owl | | | |
| White Hawk | | | |
| Olivaceous Piculet | | | |
| Prevost's Ground-Sparrow | | | |
| Spotted Antbird | | | |
| Baird's Trogon | | | |
| Blue-footed boobies | | | 1 |
| Brown Noddy | | | 1 |

##### 4.3 BEQUEST

**Table S11. List of species or groups with the associated mentions by birdwatchers, farmers, and urbanites.** Frequencies are reported in terms of how many times each species/group was mentioned when participants were asked to say which species they would like to protect for future generations.

| Species or group | Birdwatchers | Farmers | Urbanites |
|---|---|---|---|
| All of them | 41 | 62 | 48 |
| Toucans | 3 | 25 | 35 |
| Scarlett Macaw | 6 | 23 | 38 |
| Clay-colored Thrush | | 20 | 29 |
| Parrots | 2 | 15 | 10 |
| Parakeet | | 12 | 8 |
| Jabiru | 26 | 9 | 5 |
| Chachalacas | 2 | 8 | 1 |
| Long-tailed Manakin | 3 | 7 | 8 |
| Doves | | 4 | 3 |
| Roseate Spoonbill | 3 | 4 | 1 |
| Crested Bobwhite | | 4 | 1 |
| Hawks | | 3 | |
| Great Currassow | 2 | 3 | |
| Hummingbirds | 3 | 2 | 13 |
| White-throated Magpie-Jay | 3 | 2 | 7 |
| Woodpeckers | | 2 | 5 |

| Species or group | Birdwatchers | Farmers | Urbanites |
|---|---|---|---|
| Orioles | | 2 | 5 |
| Motmots | 3 | 2 | 3 |
| Herons | 1 | 2 | 2 |
| Great Green Macaw | 3 | 2 | 2 |
| Three-wattled Bell-bird | 4 | 2 | 2 |
| Canaries | | 2 | 1 |
| Great-tailed Grackle | | 2 | 1 |
| Thicket Tinamou | 1 | 2 | |
| Resplandescent Quetzal | 2 | 1 | 5 |
| Flycatcher with yellow breast | | 1 | 4 |
| Peacock | | 1 | 2 |
| Red-winged Blackbird | | 1 | 2 |
| Ornate Hawk-Eagle | 2 | 1 | 1 |
| Vultures | | 1 | 1 |
| Owls | | 1 | 1 |
| Great Egret | | 1 | 1 |
| Tiger Heron | | 1 | 1 |
| Oropendolas | | 1 | 1 |
| Seed finches | | 1 | 1 |
| Euphonia | | 1 | |
| Water birds | 1 | 1 | |
| Seed dispersers | 2 | 1 | |
| Cattle Egret | | 1 | |
| Fulvous Whistling Duck | | 1 | |
| Yellow-naped Parrot | 4 | 1 | |
| Crested Caracara | 1 | | 4 |
| Rufous-naped Wren | | | 4 |
| Raptors | 5 | | 3 |
| Swallows | | | 2 |
| Barn Owl | | | 2 |
| King Vulture | | | 2 |
| Rufous-necked Wood Rail | | | 1 |
| Common Pauraque | | | 1 |
| Laughing Falcon | 1 | | 1 |
| King fishers | 1 | | 1 |
| Olivaceous Piculet | | | 1 |
| Ocellated Antbird | | | 1 |
| Ochre-breasted Antpitta | | | 1 |
| Painted bunting | 1 | | 1 |
| Potoo | | | 1 |
| White-tipped Dove | | | 1 |
| Ducks | | | 1 |
| Prevost's Ground Sparrow | | | 1 |
| Warblers | 1 | | 1 |

| Species or group | Birdwatchers | Farmers | Urbanites |
|---|---|---|---|
| Spotted Antbird | | | 1 |
| Tody Motmot | 2 | | 1 |
| Elegant Trogon | 2 | | 1 |
| Trogons | | | 1 |
| Yellow-eared Toucanet | | | 1 |
| Flycatchers | 1 | | |
| Dry forest birds | 4 | | |
| Mangrove birds | 1 | | |
| Endangered species | 1 | | |
| Birds endemic to Guanacaste | 2 | | |
| Migratory birds | 2 | | |
| Shorebirds | 2 | | |
| Resident species | 1 | | |
| Black-headed Trogon | 1 | | |
| Botteri's Sparrow | 1 | | |
| Burrowing Owl | 1 | | |
| Mangrove Hummingbird | 2 | | |
| Ecosystems | 1 | | |
| Boat-billed Night-Heron | 1 | | |
| White Hawk | 1 | | |
| Ant birds | 2 | | |
| Ivory-billed Woodcreeper | 1 | | |
| King Vulture | 1 | | |
| Lesser Swallow-tailed Swift | 1 | | |
| Manakins | 2 | | |
| Lineated Woodpecker | 1 | | |
| Bare-necked Umbrella Bird | 1 | | |
| Anhinga | | | |
| Crested Guan | 1 | | |
| Rock Wren | 1 | | |
| Rufous-Vented Ground-Cuckoo | 1 | | |
| Rusty Sparrow | 1 | | |
| Southern Lapwing | 1 | | |
| Striped Cuckoo | 1 | | |
| Baird's Trogon | 1 | | |
| Chestnut-mandibled Toucan or Swainsons' Toucan | 1 | | |
| White-tailed Nightjar | 1 | | |

#### 5 RESULTS FROM THE CORRELATIONS WITH GENERAL LIKING OF A SPECIES

**Table S12. Correlations between the general liking of species and their scores on all six CES categories.**

| Data | Variables | t | df | $R^2$ | significance level |
|---|---|---|---|---|---|
| All data | General liking-Disservices | -40.14 | 3380 | -0.57 | <0.0001 |
| | General liking-Education | 41.52 | 3380 | 0.58 | <0.0001 |
| | General liking-Birdwatching | 74.48 | 3380 | 0.79 | <0.0001 |
| | General liking-Acoustic aesthetics | 28.30 | 3380 | 0.44 | <0.0001 |
| | General liking-Identity | 21.66 | 3380 | 0.35 | <0.0001 |
| | General liking-Bequest | 48.56 | 5123 | 0.56 | <0.0001 |
| Birdwatchers | General liking-Disservices | -16.86 | 1277 | -0.42 | <0.0001 |
| | General liking-Education | 18.71 | 1277 | 0.46 | <0.0001 |
| | General liking-Birdwatching | 37.22 | 1277 | 0.72 | <0.0001 |
| | General liking-Acoustic aesthetics | 10.96 | 1277 | 0.29 | <0.0001 |
| | General liking-Identity | 10.91 | 1277 | 0.29 | <0.0001 |
| | General liking-Bequest | 21.29 | 1465 | 0.49 | <0.0001 |
| Farmers | General liking-Disservices | -24.69 | 1087 | -0.60 | <0.0001 |
| | General liking-Education | 24.64 | 1087 | 0.60 | <0.0001 |
| | General liking-Birdwatching | 44.29 | 1087 | 0.80 | <0.0001 |
| | General liking-Acoustic aesthetics | 19.81 | 1087 | 0.51 | <0.0001 |
| | General liking-Identity | 11.69 | 1087 | 0.33 | <0.0001 |
| | General liking-Bequest | 29.08 | 1774 | 0.57 | <0.0001 |
| Urbanites | General liking-Disservices | -22.56 | 1012 | -0.58 | <0.0001 |
| | General liking-Education | 22.75 | 1012 | 0.58 | <0.0001 |
| | General liking-Birdwatching | 43.63 | 1012 | 0.80 | <0.0001 |
| | General liking-Acoustic aesthetics | 20.48 | 1012 | 0.54 | <0.0001 |
| | General liking-Identity | 16.12 | 1012 | 0.45 | <0.0001 |
| | General liking-Bequest | 29.80 | 1880 | 0.57 | <0.0001 |

### 6 COPY OF THE SURVEY

**Survey about the birds of northwestern, Costa Rica**

Thank you for your interest in completing this survey. Completing this survey will require approximately 35 minutes. This study is conducted by researchers from the University of British Columbia, Canada, in partnership with local Birdwatchers in Costa Rica.

The goal of this survey is to better understand people's perceptions of birds in Guanacaste and is part of the doctoral research of Alejandra Echeverri Ochoa, a student at the University of British Columbia. All information received will remain confidential. Only aggregated results will be released. No risk is anticipated from your participation in this study.

By completing this questionnaire, you are consenting to participate in this research. Your participation in this study is voluntary and you have the right to refuse to participate. If you decide to take part, you may choose to exit the study at any time without giving a reason and without any negative consequence.

If you have any concerns or complaints about your rights as a research participant and/or your experiences while participating in this study, please contact the Research Participant Complaint Line in the UBC Office of Research Ethics at 604-822-8598 (in Canada), or, if long distance, or call toll free 1-877-822-8598.

All the images in this survey are from the book of The Birds of Costa Rica (A field Guide) by Richard Garrigues and Robert Dean. The sound recordings presented in this survey are from the Xeno-canto website with creative commons licenses. This survey is intended to have only academic purposes.

Please do not hesitate to contact us if you have any questions regarding our research project or how the results will be used.
Kai M.A. Chan, Principal Investigator and Full Professor, Institute for Resources, Environment, and Sustainability, University of British Columbia.
Alejandra Echeverri Ochoa, Co-investigator and PhD Candidate, Institute for Resources, Environment, and Sustainability, University of British Columbia.
Jiaying Zhao, Co-investigator, Department of Psychology, and Institute for Resources, Environment, and Sustainability. University of British Columbia.

Q1 Which of the following groups do you most identify with?

- ◯ A local resident in Costa Rica, not a farmer (1)
- ◯ A farmer in Costa Rica (2)
- ◯ A birder/ ornithologist (4)
- ◯ A bird-watching guide (3)

*Skip To: End of Block If Which of the following groups do you most identify with? = A local resident in Costa Rica, not a farmer*

*Skip To: Q3 If Which of the following groups do you most identify with? = A farmer in Costa Rica*

*Skip To: Q28 If Which of the following groups do you most identify with? = A birder/ ornithologist*

*Skip To: Q28 If Which of the following groups do you most identify with? = A bird-watching guide*

Q3 What type of farm do you have?

- ◯ Crop farm (2)
- ◯ Cattle farm (3)
- ◯ Mixed (4)

Q4 What do you grow on your farm?

________________________________________________________________

________________________________________________________________

________________________________________________________________

________________________________________________________________

________________________________________________________________

*Skip To: End of Block If What do you grow on your farm? Is Not Empty*

Q28 When birding in Guanacaste, where did you go? or where do you normally go? Please name the parks, protected areas, or approximate locations of the farms, mountains, wetlands, or other places that you visit when you go or went bird-watching (if not applicable, please write: I have not been in Guanacaste and please name the closest region you have visited instead)

________________________________________________________________

__________________________________________________________________

__________________________________________________________________

Q29 If you are an international birder, where did you stay in your trip?

__________________________________________________________________

__________________________________________________________________

__________________________________________________________________

__________________________________________________________________

__________________________________________________________________

**Start of Block: Prompting questions**

P1
**Think about birds from the Northwestern part of Costa Rica, specifically the Guanacaste region**
Can you think of any birds that you enjoy watching or hearing? Can you please tell me why you like them? All kinds of answers are useful, including about how they look, sound, act, or any other significance of the bird to you, your family, or your community.

__________________________________________________________________

__________________________________________________________________

__________________________________________________________________

__________________________________________________________________

__________________________________________________________________

P3 Now thinking specifically about the future, can you think of any birds that you would like to protect for your (future) children or future generations? Can you please tell me what it is about those birds that make them especially important to protect for the future?

__________________________________________________________________

__________________________________________________________________

__________________________________________________________________

__________________________________________________________________

__________________________________________________________________

P2 Apart from birds that you enjoy watching or hearing, are there any birds that you consider problematic? Are there any birds that you dislike? Can you please tell me why you dislike them? Think about the birds that you do not enjoy watching or hearing, or the ones that damage your properties or harm you in any way

________________________________________________________________

**Start of Block: Species presented in random order**

GBAN **Please watch and listen carefully**

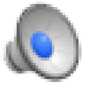

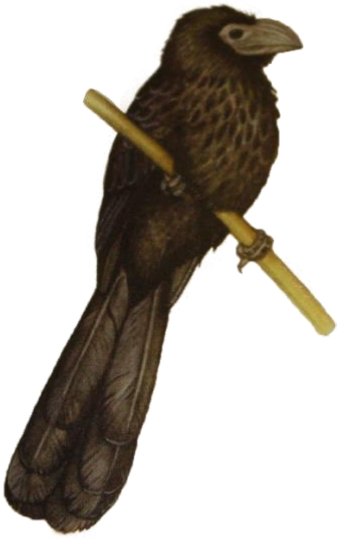

Illustration by Robert Dean (Garrigues & Dean, 2009), song from xeno-canto.org

GBAN_Q1 How much do you like this bird?

○ Dislike a great deal (1)

○ Dislike somewhat (2)

○ Neither like nor dislike (3)

○ Like somewhat (4)

○ Like a great deal (5)

GBAN_Q2 Why do you like or dislike this bird?

________________________________________________________________

________________________________________________________________

GBAN_Q3 How often do you see or hear this bird in a typical month?

- Never (1)
- Once (2)
- A few times (3)
- Many times (4)
- Every day (5)

GBAN_Q4 Do you know what this bird is?

- Yes (1)
- No (3)

*Skip To: GBAN_Q7 If Do you know what this bird is? = No*

GBAN_Q5 What do you call this bird? (You can enter N/A if you do not have a name for this bird)

________________________________________________________________

GBAN_Q6 On a scale from "strongly disagree" to "strongly agree" please tell me how much do you agree with each statement (Presented in random order)

| | **Strongly disagree** (1) | **Somewhat disagree** (2) | **Neither agree nor disagree** (3) | **Somewhat agree** (4) | **Strongly agree** (5) |
|---|---|---|---|---|---|
| This bird is like my neighbor and makes me feel at home | ○ | ○ | ○ | ○ | ○ |
| This bird helps make this place what it is | ○ | ○ | ○ | ○ | ○ |
| This bird is beautiful and I enjoy watching it | ○ | ○ | ○ | ○ | ○ |
| This bird has a beautiful song | ○ | ○ | ○ | ○ | ○ |
| I am excited to find this bird | ○ | ○ | ○ | ○ | ○ |
| This bird causes problems to the crops or the farms, for example by eating the crop | ○ | ○ | ○ | ○ | ○ |
| This bird helps the crops, for example by controlling insect pests or rodents | ○ | ○ | ○ | ○ | ○ |
| This bird causes problems to other species that are important for me | ○ | ○ | ○ | ○ | ○ |
| I like learning about or studying this bird, where it lives and what it does | ○ | ○ | ○ | ○ | ○ |
| I like teaching others about this bird and its habitat | ○ | ○ | ○ | ○ | ○ |
| I find this bird annoying because it's too noisy | ○ | ○ | ○ | ○ | ○ |
| I dislike this bird because their droppings make a mess or they build nests in inconvenient places | ○ | ○ | ○ | ○ | ○ |

GBAN_Q7 On a scale from "strongly disagree" to "strongly agree" please tell me how much do you agree with each statement

| | **Strongly disagree** (1) | **Somewhat disagree** (2) | **Neither agree nor disagree** (3) | **Somewhat agree** (4) | **Strongly agree** (5) |
|---|---|---|---|---|---|
| This bird should be protected for future generations | ○ | ○ | ○ | ○ | ○ |
| It would be sad if this bird would no longer exist | ○ | ○ | ○ | ○ | ○ |

GBAN_Q8 Do you have any memories with this bird? Do you have any comments about this bird?

________________________________________________________________

________________________________________________________________

________________________________________________________________

________________________________________________________________

________________________________________________________________

Start of Block: Clay-colored Thrush

CCRO **Please watch and listen carefully**

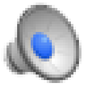

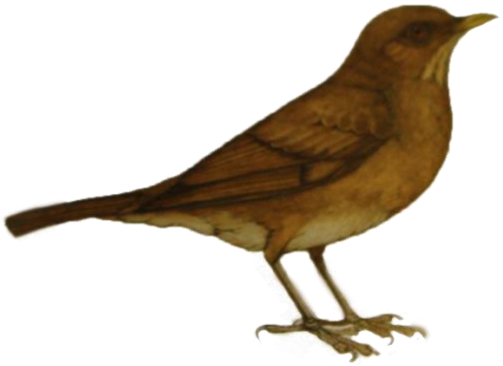

Illustration by Robert Dean (Garrigues & Dean, 2009), song from xeno-canto.org

CCRO_Q1 How much do you like this bird?

○ Dislike a great deal (1)

○ Dislike somewhat (2)

○ Neither like nor dislike (3)

○ Like somewhat (4)

○ Like a great deal (5)

CCRO_Q2 Why do you like or dislike this bird?

________________________________________________________________

________________________________________________________________

CCRO_Q3 How often do you see or hear this bird in a typical month?

- Never (1)
- Once (2)
- A few times (3)
- Many times (4)
- Every day (5)

CCRO_Q4 Do you know what this bird is?

- Yes (1)
- No (3)

*Skip To: CCRO_Q7 If Do you know what this bird is? = No*

CCRO_Q5 What do you call this bird? (You can enter N/A if you do not have a name for this bird)

________________________________________________________________

CCRO_Q6 On a scale from "strongly disagree" to "strongly agree" please tell me how much do you agree with each statement (Presented in random order)

| | **Strongly disagree** (1) | **Somewhat disagree** (2) | **Neither agree nor disagree** (3) | **Somewhat agree** (4) | **Strongly agree** (5) |
|---|---|---|---|---|---|
| This bird is like my neighbor and makes me feel at home | ○ | ○ | ○ | ○ | ○ |
| This bird helps make this place what it is | ○ | ○ | ○ | ○ | ○ |
| This bird is beautiful and I enjoy watching it | ○ | ○ | ○ | ○ | ○ |
| This bird has a beautiful song | ○ | ○ | ○ | ○ | ○ |
| I am excited to find this bird | ○ | ○ | ○ | ○ | ○ |
| This bird causes problems to the crops or the farms, for example by eating the crop | ○ | ○ | ○ | ○ | ○ |
| This bird helps the crops, for example by controlling insect pests or rodents | ○ | ○ | ○ | ○ | ○ |
| This bird causes problems to other species that are important for me | ○ | ○ | ○ | ○ | ○ |
| I like learning about or studying this bird, where it lives and what it does | ○ | ○ | ○ | ○ | ○ |
| I like teaching others about this bird and its habitat | ○ | ○ | ○ | ○ | ○ |
| I find this bird annoying because it's too noisy | ○ | ○ | ○ | ○ | ○ |
| I dislike this bird because their droppings make a mess or they build nests in inconvenient places | ○ | ○ | ○ | ○ | ○ |

CCRO_Q7 On a scale from "strongly disagree" to "strongly agree" please tell me how much do you agree with each statement

| | **Strongly disagree** (1) | **Somewhat disagree** (2) | **Neither agree nor disagree** (3) | **Somewhat agree** (4) | **Strongly agree** (5) |
|---|---|---|---|---|---|
| This bird should be protected for future generations | ○ | ○ | ○ | ○ | ○ |
| It would be sad if this bird would no longer exist | ○ | ○ | ○ | ○ | ○ |

CCRO_Q8 Do you have any memories with this bird? Do you have any comments about this bird?

________________________________________________________________

________________________________________________________________

________________________________________________________________

________________________________________________________________

________________________________________________________________

Start of Block: Long-tailed Manakin

LTMA **Please watch and listen carefully**

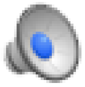

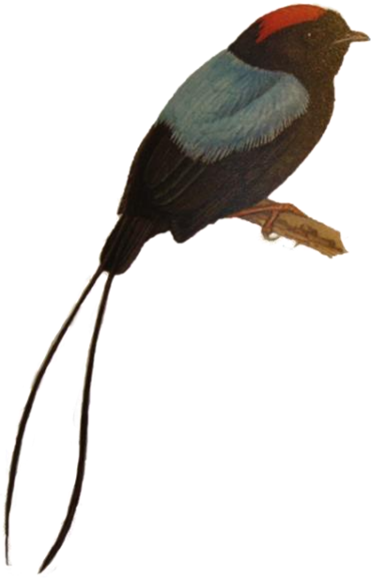

Illustration by Robert Dean (Garrigues & Dean, 2009), song from xeno-canto.org

LTMA_Q1 How much do you like this bird?

- ◯ Dislike a great deal (1)
- ◯ Dislike somewhat (2)
- ◯ Neither like nor dislike (3)
- ◯ Like somewhat (4)
- ◯ Like a great deal (5)

LTMA_Q2 Why do you like or dislike this bird?

________________________________________________________________

________________________________________________________________

LTMA_Q3 How often do you see or hear this bird in a typical month?

- ○ Never (1)
- ○ Once (2)
- ○ A few times (3)
- ○ Many times (4)
- ○ Every day (5)

LTMA_Q4 Do you know what this bird is?

- ○ Yes (1)
- ○ No (3)

*Skip To: LTMA_Q7 If Do you know what this bird is? = No*

LTMA_Q5 What do you call this bird? (You can enter N/A if you do not have a name for this bird)

________________________________________________________________

LTMA_Q6 On a scale from "strongly disagree" to "strongly agree" please tell me how much do you agree with each statement (Presented in random order)

| | **Strongly disagree** (1) | **Somewhat disagree** (2) | **Neither agree nor disagree** (3) | **Somewhat agree** (4) | **Strongly agree** (5) |
|---|---|---|---|---|---|
| This bird is like my neighbor and makes me feel at home | ○ | ○ | ○ | ○ | ○ |
| This bird helps make this place what it is | ○ | ○ | ○ | ○ | ○ |
| This bird is beautiful and I enjoy watching it | ○ | ○ | ○ | ○ | ○ |
| This bird has a beautiful song | ○ | ○ | ○ | ○ | ○ |
| I am excited to find this bird | ○ | ○ | ○ | ○ | ○ |
| This bird causes problems to the crops or the farms, for example by eating the crop | ○ | ○ | ○ | ○ | ○ |
| This bird helps the crops, for example by controlling insect pests or rodents | ○ | ○ | ○ | ○ | ○ |
| This bird causes problems to other species that are important for me | ○ | ○ | ○ | ○ | ○ |
| I like learning about or studying this bird, where it lives and what it does | ○ | ○ | ○ | ○ | ○ |
| I like teaching others about this bird and its habitat | ○ | ○ | ○ | ○ | ○ |
| I find this bird annoying because it's too noisy | ○ | ○ | ○ | ○ | ○ |
| I dislike this bird because their droppings make a mess or they build nests in inconvenient places | ○ | ○ | ○ | ○ | ○ |

LTMA_Q7 On a scale from "strongly disagree" to "strongly agree" please tell me how much do you agree with each statement

| | **Strongly disagree** (1) | **Somewhat disagree** (2) | **Neither agree nor disagree** (3) | **Somewhat agree** (4) | **Strongly agree** (5) |
|---|---|---|---|---|---|
| This bird should be protected for future generations | ○ | ○ | ○ | ○ | ○ |
| It would be sad if this bird would no longer exist | ○ | ○ | ○ | ○ | ○ |

LTMA_Q8 Do you have any memories with this bird? Do you have any comments about this bird?

________________________________________________________________

________________________________________________________________

________________________________________________________________

________________________________________________________________

________________________________________________________________

Start of Block: White-throated Magpie-Jay

WTMJ **Please watch and listen carefully**

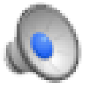

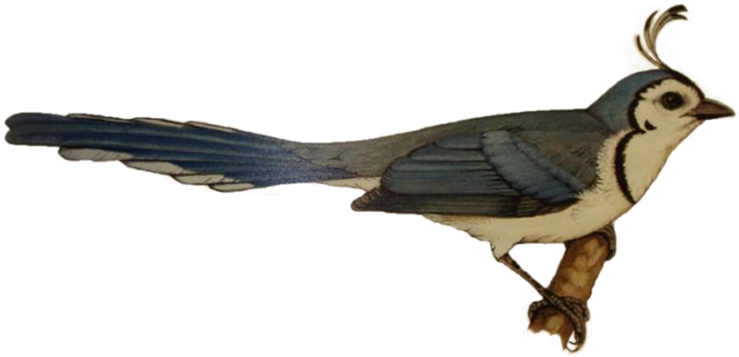

Illustration by Robert Dean (Garrigues & Dean, 2009), song from xeno-canto.org

WTMJ_Q1 How much do you like this bird?

- ◯ Dislike a great deal (1)
- ◯ Dislike somewhat (2)
- ◯ Neither like nor dislike (3)
- ◯ Like somewhat (4)
- ◯ Like a great deal (5)

WTMJ_Q2 Why do you like or dislike this bird?

________________________________________________________________

________________________________________________________________

WTMJ_Q3 How often do you see or hear this bird in a typical month?

- ○ Never (1)
- ○ Once (2)
- ○ A few times (3)
- ○ Many times (4)
- ○ Every day (5)

WTMJ_Q4 Do you know what this bird is?

- ○ Yes (1)
- ○ No (3)

*Skip To: WTMJ_Q7 If Do you know what this bird is? = No*

WTMJ_Q5 What do you call this bird? (You can enter N/A if you do not have a name for this bird)

________________________________________________________________

WTMJ_Q6 On a scale from "strongly disagree" to "strongly agree" please tell me how much do you agree with each statement (Presented in random order)

| | **Strongly disagree** (1) | **Somewhat disagree** (2) | **Neither agree nor disagree** (3) | **Somewhat agree** (4) | **Strongly agree** (5) |
|---|---|---|---|---|---|
| This bird is like my neighbor and makes me feel at home | ○ | ○ | ○ | ○ | ○ |
| This bird helps make this place what it is | ○ | ○ | ○ | ○ | ○ |
| This bird is beautiful and I enjoy watching it | ○ | ○ | ○ | ○ | ○ |
| This bird has a beautiful song | ○ | ○ | ○ | ○ | ○ |
| I am excited to find this bird | ○ | ○ | ○ | ○ | ○ |
| This bird causes problems to the crops or the farms, for example by eating the crop | ○ | ○ | ○ | ○ | ○ |
| This bird helps the crops, for example by controlling insect pests or rodents | ○ | ○ | ○ | ○ | ○ |
| This bird causes problems to other species that are important for me | ○ | ○ | ○ | ○ | ○ |
| I like learning about or studying this bird, where it lives and what it does | ○ | ○ | ○ | ○ | ○ |
| I like teaching others about this bird and its habitat | ○ | ○ | ○ | ○ | ○ |
| I find this bird annoying because it's too noisy | ○ | ○ | ○ | ○ | ○ |
| I dislike this bird because their droppings make a mess or they build nests in inconvenient places | ○ | ○ | ○ | ○ | ○ |

WTMJ_Q7 On a scale from "strongly disagree" to "strongly agree" please tell me how much do you agree with each statement

| | **Strongly disagree** (1) | **Somewhat disagree** (2) | **Neither agree nor disagree** (3) | **Somewhat agree** (4) | **Strongly agree** (5) |
|---|---|---|---|---|---|
| This bird should be protected for future generations | ○ | ○ | ○ | ○ | ○ |
| It would be sad if this bird would no longer exist | ○ | ○ | ○ | ○ | ○ |

WTMJ_Q8 Do you have any memories with this bird? Do you have any comments about this bird?

________________________________________________________________

________________________________________________________________

________________________________________________________________

________________________________________________________________

________________________________________________________________

Start of Block: Swallow-tailed Kite

STKI **Please watch and listen carefully**

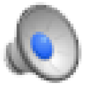
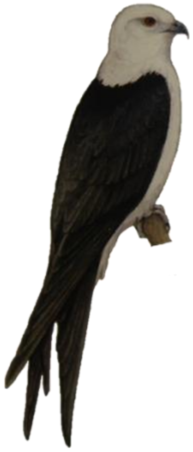

Illustration by Robert Dean (Garrigues & Dean, 2009), song from xeno-canto.org

STKI_Q1 How much do you like this bird?

○ Dislike a great deal (1)

○ Dislike somewhat (2)

○ Neither like nor dislike (3)

○ Like somewhat (4)

○ Like a great deal (5)

STKI_Q2 Why do you like or dislike this bird?

________________________________________________________________

________________________________________________________________

STKI_Q3 How often do you see or hear this bird in a typical month?

- ○ Never (1)
- ○ Once (2)
- ○ A few times (3)
- ○ Many times (4)
- ○ Every day (5)

STKI_Q4 Do you know what this bird is?

- ○ Yes (1)
- ○ No (3)

*Skip To: STKI_Q7 If Do you know what this bird is? = No*

STKI_Q5 What do you call this bird? (You can enter N/A if you do not have a name for this bird)

________________________________________________________________

STKI_Q6 On a scale from "strongly disagree" to "strongly agree" please tell me how much do you agree with each statement (Presented in random order)

| | **Strongly disagree** (1) | **Somewhat disagree** (2) | **Neither agree nor disagree** (3) | **Somewhat agree** (4) | **Strongly agree** (5) |
|---|---|---|---|---|---|
| This bird is like my neighbor and makes me feel at home | ○ | ○ | ○ | ○ | ○ |
| This bird helps make this place what it is | ○ | ○ | ○ | ○ | ○ |
| This bird is beautiful and I enjoy watching it | ○ | ○ | ○ | ○ | ○ |
| This bird has a beautiful song | ○ | ○ | ○ | ○ | ○ |
| I am excited to find this bird | ○ | ○ | ○ | ○ | ○ |
| This bird causes problems to the crops or the farms, for example by eating the crop | ○ | ○ | ○ | ○ | ○ |
| This bird helps the crops, for example by controlling insect pests or rodents | ○ | ○ | ○ | ○ | ○ |
| This bird causes problems to other species that are important for me | ○ | ○ | ○ | ○ | ○ |
| I like learning about or studying this bird, where it lives and what it does | ○ | ○ | ○ | ○ | ○ |
| I like teaching others about this bird and its habitat | ○ | ○ | ○ | ○ | ○ |
| I find this bird annoying because it's too noisy | ○ | ○ | ○ | ○ | ○ |
| I dislike this bird because their droppings make a mess or they build nests in inconvenient places | ○ | ○ | ○ | ○ | ○ |

STKI_Q7 On a scale from "strongly disagree" to "strongly agree" please tell me how much do you agree with each statement

| | **Strongly disagree** (1) | **Somewhat disagree** (2) | **Neither agree nor disagree** (3) | **Somewhat agree** (4) | **Strongly agree** (5) |
|---|---|---|---|---|---|
| This bird should be protected for future generations | o | o | o | o | o |
| It would be sad if this bird would no longer exist | o | o | o | o | o |

STKI_Q8 Do you have any memories with this bird? Do you have any comments about this bird?

________________________________________________________________

________________________________________________________________

________________________________________________________________

________________________________________________________________

________________________________________________________________

Start of Block: Lesser Swallow-tailed Swift

LSTS **Please watch and listen carefully**

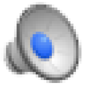

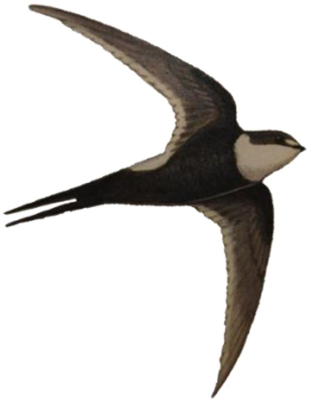

Illustration by Robert Dean (Garrigues & Dean, 2009), song from xeno-canto.org

LSTS_Q1 How much do you like this bird?

- ◯ Dislike a great deal (1)
- ◯ Dislike somewhat (2)
- ◯ Neither like nor dislike (3)
- ◯ Like somewhat (4)
- ◯ Like a great deal (5)

LSTS_Q2 Why do you like or dislike this bird?

________________________________________________________________

________________________________________________________________

LSTS_Q3 How often do you see or hear this bird in a typical month?

- ○ Never (1)
- ○ Once (2)
- ○ A few times (3)
- ○ Many times (4)
- ○ Every day (5)

LSTS_Q4 Do you know what this bird is?

- ○ Yes (1)
- ○ No (3)

*Skip To: LSTS_Q7 If Do you know what this bird is? = No*

LSTS_Q5 What do you call this bird? (You can enter N/A if you do not have a name for this bird)

________________________________________________________________

LSTS_Q6 On a scale from "strongly disagree" to "strongly agree" please tell me how much do you agree with each statement (Presented in random order)

| | **Strongly disagree** (1) | **Somewhat disagree** (2) | **Neither agree nor disagree** (3) | **Somewhat agree** (4) | **Strongly agree** (5) |
|---|---|---|---|---|---|
| This bird is like my neighbor and makes me feel at home | o | o | o | o | o |
| This bird helps make this place what it is | o | o | o | o | o |
| This bird is beautiful and I enjoy watching it | o | o | o | o | o |
| This bird has a beautiful song | o | o | o | o | o |
| I am excited to find this bird | o | o | o | o | o |
| This bird causes problems to the crops or the farms, for example by eating the crop | o | o | o | o | o |
| This bird helps the crops, for example by controlling insect pests or rodents | o | o | o | o | o |
| This bird causes problems to other species that are important for me | o | o | o | o | o |
| I like learning about or studying this bird, where it lives and what it does | o | o | o | o | o |
| I like teaching others about this bird and its habitat | o | o | o | o | o |
| I find this bird annoying because it's too noisy | o | o | o | o | o |
| I dislike this bird because their droppings make a mess or they build nests in inconvenient places | o | o | o | o | o |

LSTS_Q7 On a scale from "strongly disagree" to "strongly agree" please tell me how much do you agree with each statement

| | **Strongly disagree** (1) | **Somewhat disagree** (2) | **Neither agree nor disagree** (3) | **Somewhat agree** (4) | **Strongly agree** (5) |
|---|---|---|---|---|---|
| This bird should be protected for future generations | ○ | ○ | ○ | ○ | ○ |
| It would be sad if this bird would no longer exist | ○ | ○ | ○ | ○ | ○ |

LSTS_Q8 Do you have any memories with this bird? Do you have any comments about this bird?

________________________________________________________________

________________________________________________________________

________________________________________________________________

________________________________________________________________

________________________________________________________________

Start of Block: Wood Stork

WOST **Please watch and listen carefully**

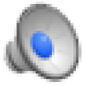

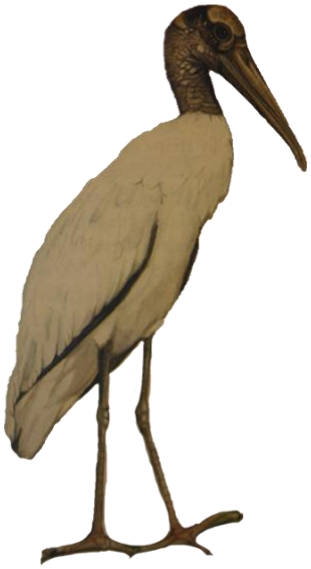

Illustration by Robert Dean (Garrigues & Dean, 2009), song from xeno-canto.org

WOST_Q1 How much do you like this bird?

- ◯ Dislike a great deal (1)
- ◯ Dislike somewhat (2)
- ◯ Neither like nor dislike (3)
- ◯ Like somewhat (4)
- ◯ Like a great deal (5)

WOST_Q2 Why do you like or dislike this bird?

________________________________________________________________

________________________________________________________________

WOST_Q3 How often do you see or hear this bird in a typical month?

- Never (1)
- Once (2)
- A few times (3)
- Many times (4)
- Every day (5)

WOST_Q4 Do you know what this bird is?

- Yes (1)
- No (3)

*Skip To: WOST_Q7 If Do you know what this bird is? = No*

WOST_Q5 What do you call this bird? (You can enter N/A if you do not have a name for this bird)

________________________________________________________________

WOST_Q6 On a scale from "strongly disagree" to "strongly agree" please tell me how much do you agree with each statement (Presented in random order)

| | **Strongly disagree** (1) | **Somewhat disagree** (2) | **Neither agree nor disagree** (3) | **Somewhat agree** (4) | **Strongly agree** (5) |
|---|---|---|---|---|---|
| This bird is like my neighbor and makes me feel at home | | | | | |
| This bird helps make this place what it is | | | | | |
| This bird is beautiful and I enjoy watching it | | | | | |
| This bird has a beautiful song | | | | | |
| I am excited to find this bird | | | | | |
| This bird causes problems to the crops or the farms, for example by eating the crop | | | | | |
| This bird helps the crops, for example by controlling insect pests or rodents | | | | | |
| This bird causes problems to other species that are important for me | | | | | |
| I like learning about or studying this bird, where it lives and what it does | | | | | |
| I like teaching others about this bird and its habitat | | | | | |
| I find this bird annoying because it's too noisy | | | | | |
| I dislike this bird because their droppings make a mess or they build nests in inconvenient places | | | | | |

WOST_Q7 On a scale from "strongly disagree" to "strongly agree" please tell me how much do you agree with each statement

| | **Strongly disagree** (1) | **Somewhat disagree** (2) | **Neither agree nor disagree** (3) | **Somewhat agree** (4) | **Strongly agree** (5) |
|---|---|---|---|---|---|
| This bird should be protected for future generations | ○ | ○ | ○ | ○ | ○ |
| It would be sad if this bird would no longer exist | ○ | ○ | ○ | ○ | ○ |

WOST_Q8 Do you have any memories with this bird? Do you have any comments about this bird?

________________________________________________________________

________________________________________________________________

________________________________________________________________

________________________________________________________________

________________________________________________________________

Start of Block: Laughing Falcon

LAFA **Please watch and listen carefully**

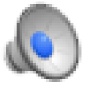

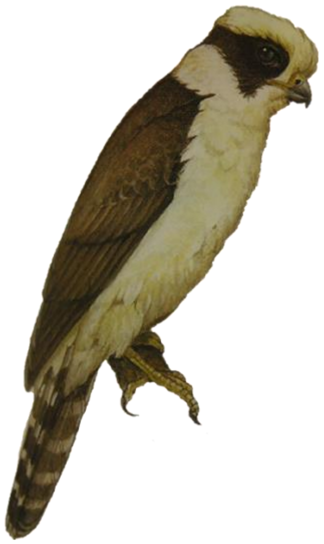

Illustration by Robert Dean (Garrigues & Dean, 2009), song from xeno-canto.org

LAFA_Q1 How much do you like this bird?

- ○ Dislike a great deal (1)
- ○ Dislike somewhat (2)
- ○ Neither like nor dislike (3)
- ○ Like somewhat (4)
- ○ Like a great deal (5)

LAFA_Q2 Why do you like or dislike this bird?

________________________________________________________________

________________________________________________________________

LAFA_Q3 How often do you see or hear this bird in a typical month?

- ○ Never (1)
- ○ Once (2)
- ○ A few times (3)
- ○ Many times (4)
- ○ Every day (5)

LAFA_Q4 Do you know what this bird is?

- ○ Yes (1)
- ○ No (3)

*Skip To: LAFA_Q7 If Do you know what this bird is? = No*

LAFA_Q5 What do you call this bird? (You can enter N/A if you do not have a name for this bird)

________________________________________________________________

LAFA_Q6 On a scale from "strongly disagree" to "strongly agree" please tell me how much do you agree with each statement (Presented in random order)

| | **Strongly disagree** (1) | **Somewhat disagree** (2) | **Neither agree nor disagree** (3) | **Somewhat agree** (4) | **Strongly agree** (5) |
|---|---|---|---|---|---|
| This bird is like my neighbor and makes me feel at home | ○ | ○ | ○ | ○ | ○ |
| This bird helps make this place what it is | ○ | ○ | ○ | ○ | ○ |
| This bird is beautiful and I enjoy watching it | ○ | ○ | ○ | ○ | ○ |
| This bird has a beautiful song | ○ | ○ | ○ | ○ | ○ |
| I am excited to find this bird | ○ | ○ | ○ | ○ | ○ |
| This bird causes problems to the crops or the farms, for example by eating the crop | ○ | ○ | ○ | ○ | ○ |
| This bird helps the crops, for example by controlling insect pests or rodents | ○ | ○ | ○ | ○ | ○ |
| This bird causes problems to other species that are important for me | ○ | ○ | ○ | ○ | ○ |
| I like learning about or studying this bird, where it lives and what it does | ○ | ○ | ○ | ○ | ○ |
| I like teaching others about this bird and its habitat | ○ | ○ | ○ | ○ | ○ |
| I find this bird annoying because it's too noisy | ○ | ○ | ○ | ○ | ○ |
| I dislike this bird because their droppings make a mess or they build nests in inconvenient places | ○ | ○ | ○ | ○ | ○ |

LAFA_Q7 On a scale from "strongly disagree" to "strongly agree" please tell me how much do you agree with each statement

| | **Strongly disagree** (1) | **Somewhat disagree** (2) | **Neither agree nor disagree** (3) | **Somewhat agree** (4) | **Strongly agree** (5) |
|---|---|---|---|---|---|
| This bird should be protected for future generations | ○ | ○ | ○ | ○ | ○ |
| It would be sad if this bird would no longer exist | ○ | ○ | ○ | ○ | ○ |

LAFA_Q8 Do you have any memories with this bird? Do you have any comments about this bird?

________________________________________________________________

________________________________________________________________

________________________________________________________________

________________________________________________________________

________________________________________________________________

Start of Block: Olivaceous Woodcreeper

OLWC **Please watch and listen carefully**

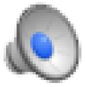 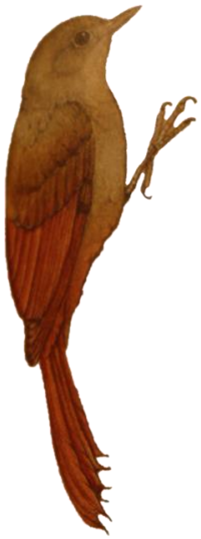

Illustration by Robert Dean (Garrigues & Dean, 2009), song from xeno-canto.org

OLWC_Q1 How much do you like this bird?

- ○ Dislike a great deal (1)
- ○ Dislike somewhat (2)
- ○ Neither like nor dislike (3)
- ○ Like somewhat (4)
- ○ Like a great deal (5)

OLWC_Q2 Why do you like or dislike this bird?

________________________________________________________________

________________________________________________________________

OLWC_Q3 How often do you see or hear this bird in a typical month?

- Never (1)
- Once (2)
- A few times (3)
- Many times (4)
- Every day (5)

OLWC_Q4 Do you know what this bird is?

- Yes (1)
- No (3)

*Skip To: OLWC_Q7 If Do you know what this bird is? = No*

OLWC_Q5 What do you call this bird? (You can enter N/A if you do not have a name for this bird)

________________________________________________________________

OLWC_Q6 On a scale from "strongly disagree" to "strongly agree" please tell me how much do you agree with each statement (Presented in random order)

| | **Strongly disagree** (1) | **Somewhat disagree** (2) | **Neither agree nor disagree** (3) | **Somewhat agree** (4) | **Strongly agree** (5) |
|---|---|---|---|---|---|
| This bird is like my neighbor and makes me feel at home | ○ | ○ | ○ | ○ | ○ |
| This bird helps make this place what it is | ○ | ○ | ○ | ○ | ○ |
| This bird is beautiful and I enjoy watching it | ○ | ○ | ○ | ○ | ○ |
| This bird has a beautiful song | ○ | ○ | ○ | ○ | ○ |
| I am excited to find this bird | ○ | ○ | ○ | ○ | ○ |
| This bird causes problems to the crops or the farms, for example by eating the crop | ○ | ○ | ○ | ○ | ○ |
| This bird helps the crops, for example by controlling insect pests or rodents | ○ | ○ | ○ | ○ | ○ |
| This bird causes problems to other species that are important for me | ○ | ○ | ○ | ○ | ○ |
| I like learning about or studying this bird, where it lives and what it does | ○ | ○ | ○ | ○ | ○ |
| I like teaching others about this bird and its habitat | ○ | ○ | ○ | ○ | ○ |
| I find this bird annoying because it's too noisy | ○ | ○ | ○ | ○ | ○ |
| I dislike this bird because their droppings make a mess or they build nests in inconvenient places | ○ | ○ | ○ | ○ | ○ |

OLWC_Q7 On a scale from "strongly disagree" to "strongly agree" please tell me how much do you agree with each statement

| | **Strongly disagree** (1) | **Somewhat disagree** (2) | **Neither agree nor disagree** (3) | **Somewhat agree** (4) | **Strongly agree** (5) |
|---|---|---|---|---|---|
| This bird should be protected for future generations | ○ | ○ | ○ | ○ | ○ |
| It would be sad if this bird would no longer exist | ○ | ○ | ○ | ○ | ○ |

OLWC_Q8 Do you have any memories with this bird? Do you have any comments about this bird?

________________________________________________________________

________________________________________________________________

________________________________________________________________

________________________________________________________________

________________________________________________________________

Start of Block: Barred Antshrike

BAAS **Please watch and listen carefully**

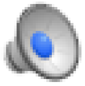

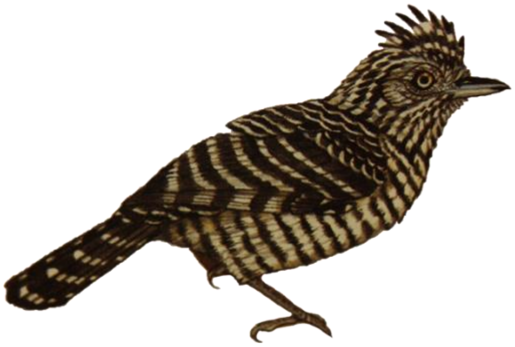

Illustration by Robert Dean (Garrigues & Dean, 2009), song from xeno-canto.org

BAAS_Q1 How much do you like this bird?

○ Dislike a great deal (1)

○ Dislike somewhat (2)

○ Neither like nor dislike (3)

○ Like somewhat (4)

○ Like a great deal (5)

BAAS_Q2 Why do you like or dislike this bird?

________________________________________________________________

________________________________________________________________

BAAS_Q3 How often do you see or hear this bird in a typical month?

- ○ Never (1)
- ○ Once (2)
- ○ A few times (3)
- ○ Many times (4)
- ○ Every day (5)

BAAS_Q4 Do you know what this bird is?

- ○ Yes (1)
- ○ No (3)

*Skip To: BAAS_Q7 If Do you know what this bird is? = No*

BAAS_Q5 What do you call this bird? (You can enter N/A if you do not have a name for this bird)

________________________________________________________________

BAAS_Q6 On a scale from "strongly disagree" to "strongly agree" please tell me how much do you agree with each statement (Presented in random order)

| | **Strongly disagree** (1) | **Somewhat disagree** (2) | **Neither agree nor disagree** (3) | **Somewhat agree** (4) | **Strongly agree** (5) |
|---|---|---|---|---|---|
| This bird is like my neighbor and makes me feel at home | o | o | o | o | o |
| This bird helps make this place what it is | o | o | o | o | o |
| This bird is beautiful and I enjoy watching it | o | o | o | o | o |
| This bird has a beautiful song | o | o | o | o | o |
| I am excited to find this bird | o | o | o | o | o |
| This bird causes problems to the crops or the farms, for example by eating the crop | o | o | o | o | o |
| This bird helps the crops, for example by controlling insect pests or rodents | o | o | o | o | o |
| This bird causes problems to other species that are important for me | o | o | o | o | o |
| I like learning about or studying this bird, where it lives and what it does | o | o | o | o | o |
| I like teaching others about this bird and its habitat | o | o | o | o | o |
| I find this bird annoying because it's too noisy | o | o | o | o | o |
| I dislike this bird because their droppings make a mess or they build nests in inconvenient places | o | o | o | o | o |

BAAS_Q7 On a scale from "strongly disagree" to "strongly agree" please tell me how much do you agree with each statement

| | **Strongly disagree** (1) | **Somewhat disagree** (2) | **Neither agree nor disagree** (3) | **Somewhat agree** (4) | **Strongly agree** (5) |
|---|---|---|---|---|---|
| This bird should be protected for future generations | ○ | ○ | ○ | ○ | ○ |
| It would be sad if this bird would no longer exist | ○ | ○ | ○ | ○ | ○ |

BAAS_Q8 Do you have any memories with this bird? Do you have any comments about this bird?

________________________________________________________________

________________________________________________________________

________________________________________________________________

________________________________________________________________

________________________________________________________________

Start of Block: Brown-crested Flycatcher

BCFL **Please watch and listen carefully**

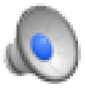

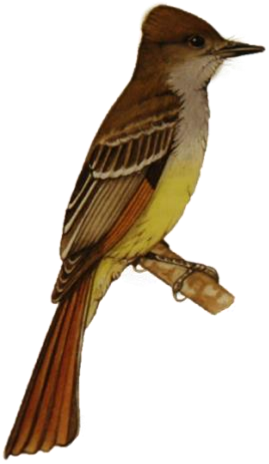

Illustration by Robert Dean (Garrigues & Dean, 2009), song from xeno-canto.org

BCFL_Q1 How much do you like this bird?

◯ Dislike a great deal (1)

◯ Dislike somewhat (2)

◯ Neither like nor dislike (3)

◯ Like somewhat (4)

◯ Like a great deal (5)

BCFL_Q2 Why do you like or dislike this bird?

________________________________________________________________

________________________________________________________________

BCFL_Q3 How often do you see or hear this bird in a typical month?

- Never (1)
- Once (2)
- A few times (3)
- Many times (4)
- Every day (5)

BCFL_Q4 Do you know what this bird is?

- Yes (1)
- No (3)

*Skip To: BCFL_Q7 If Do you know what this bird is? = No*

BCFL_Q5 What do you call this bird? (You can enter N/A if you do not have a name for this bird)

________________________________________________________________

BCFL_Q6 On a scale from "strongly disagree" to "strongly agree" please tell me how much do you agree with each statement (Presented in random order)

| | **Strongly disagree** (1) | **Somewhat disagree** (2) | **Neither agree nor disagree** (3) | **Somewhat agree** (4) | **Strongly agree** (5) |
|---|---|---|---|---|---|
| This bird is like my neighbor and makes me feel at home | ○ | ○ | ○ | ○ | ○ |
| This bird helps make this place what it is | ○ | ○ | ○ | ○ | ○ |
| This bird is beautiful and I enjoy watching it | ○ | ○ | ○ | ○ | ○ |
| This bird has a beautiful song | ○ | ○ | ○ | ○ | ○ |
| I am excited to find this bird | ○ | ○ | ○ | ○ | ○ |
| This bird causes problems to the crops or the farms, for example by eating the crop | ○ | ○ | ○ | ○ | ○ |
| This bird helps the crops, for example by controlling insect pests or rodents | ○ | ○ | ○ | ○ | ○ |
| This bird causes problems to other species that are important for me | ○ | ○ | ○ | ○ | ○ |
| I like learning about or studying this bird, where it lives and what it does | ○ | ○ | ○ | ○ | ○ |
| I like teaching others about this bird and its habitat | ○ | ○ | ○ | ○ | ○ |
| I find this bird annoying because it's too noisy | ○ | ○ | ○ | ○ | ○ |
| I dislike this bird because their droppings make a mess or they build nests in inconvenient places | ○ | ○ | ○ | ○ | ○ |

BCFL_Q7 On a scale from "strongly disagree" to "strongly agree" please tell me how much do you agree with each statement

| | **Strongly disagree** (1) | **Somewhat disagree** (2) | **Neither agree nor disagree** (3) | **Somewhat agree** (4) | **Strongly agree** (5) |
|---|---|---|---|---|---|
| This bird should be protected for future generations | ○ | ○ | ○ | ○ | ○ |
| It would be sad if this bird would no longer exist | ○ | ○ | ○ | ○ | ○ |

BCFL_Q8 Do you have any memories with this bird? Do you have any comments about this bird?

________________________________________________________________

________________________________________________________________

________________________________________________________________

________________________________________________________________

________________________________________________________________

Start of Block: Rufous-browed Peppershrike

RBPS **Please watch and listen carefully**

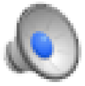

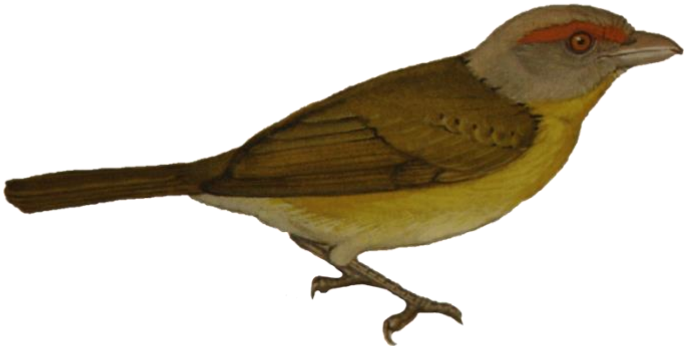

Illustration by Robert Dean (Garrigues & Dean, 2009), song from xeno-canto.org

RBPS_Q1 How much do you like this bird?

- Dislike a great deal (1)
- Dislike somewhat (2)
- Neither like nor dislike (3)
- Like somewhat (4)
- Like a great deal (5)

RBPS_Q2 Why do you like or dislike this bird?

________________________________________________________________

________________________________________________________________

RBPS_Q3 How often do you see or hear this bird in a typical month?

- ◯ Never (1)
- ◯ Once (2)
- ◯ A few times (3)
- ◯ Many times (4)
- ◯ Every day (5)

RBPS_Q4 Do you know what this bird is?

- ◯ Yes (1)
- ◯ No (3)

*Skip To: RBPS_Q7 If Do you know what this bird is? = No*

RBPS_Q5 What do you call this bird? (You can enter N/A if you do not have a name for this bird)

________________________________________________________________

RBPS_Q6 On a scale from "strongly disagree" to "strongly agree" please tell me how much do you agree with each statement (Presented in random order)

| | **Strongly disagree** (1) | **Somewhat disagree** (2) | **Neither agree nor disagree** (3) | **Somewhat agree** (4) | **Strongly agree** (5) |
|---|---|---|---|---|---|
| This bird is like my neighbor and makes me feel at home | ○ | ○ | ○ | ○ | ○ |
| This bird helps make this place what it is | ○ | ○ | ○ | ○ | ○ |
| This bird is beautiful and I enjoy watching it | ○ | ○ | ○ | ○ | ○ |
| This bird has a beautiful song | ○ | ○ | ○ | ○ | ○ |
| I am excited to find this bird | ○ | ○ | ○ | ○ | ○ |
| This bird causes problems to the crops or the farms, for example by eating the crop | ○ | ○ | ○ | ○ | ○ |
| This bird helps the crops, for example by controlling insect pests or rodents | ○ | ○ | ○ | ○ | ○ |
| This bird causes problems to other species that are important for me | ○ | ○ | ○ | ○ | ○ |
| I like learning about or studying this bird, where it lives and what it does | ○ | ○ | ○ | ○ | ○ |
| I like teaching others about this bird and its habitat | ○ | ○ | ○ | ○ | ○ |
| I find this bird annoying because it's too noisy | ○ | ○ | ○ | ○ | ○ |
| I dislike this bird because their droppings make a mess or they build nests in inconvenient places | ○ | ○ | ○ | ○ | ○ |

RBPS_Q7 On a scale from "strongly disagree" to "strongly agree" please tell me how much do you agree with each statement

| | **Strongly disagree** (1) | **Somewhat disagree** (2) | **Neither agree nor disagree** (3) | **Somewhat agree** (4) | **Strongly agree** (5) |
|---|---|---|---|---|---|
| This bird should be protected for future generations | o | o | o | o | o |
| It would be sad if this bird would no longer exist | o | o | o | o | o |

RBPS_Q8 Do you have any memories with this bird? Do you have any comments about this bird?

________________________________________________________________

________________________________________________________________

________________________________________________________________

________________________________________________________________

________________________________________________________________

Start of Block: Scarlett Macaw

SCMA **Please watch and listen carefully**

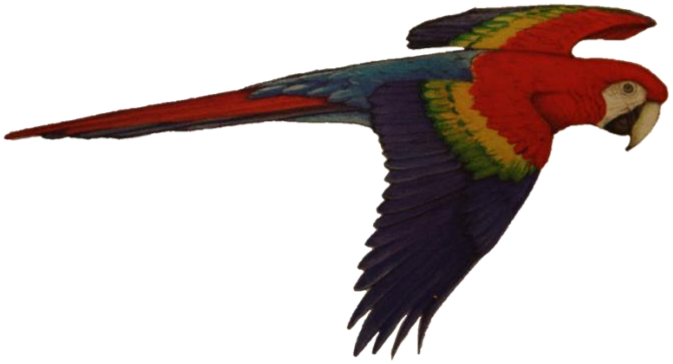

Illustration by Robert Dean (Garrigues & Dean, 2009), song from xeno-canto.org

SCMA_Q1 How much do you like this bird?

- ○ Dislike a great deal (1)
- ○ Dislike somewhat (2)
- ○ Neither like nor dislike (3)
- ○ Like somewhat (4)
- ○ Like a great deal (5)

SCMA_Q2 Why do you like or dislike this bird?

________________________________________________________________

________________________________________________________________

SCMA_Q3 How often do you see or hear this bird in a typical month?

- ◯ Never (1)
- ◯ Once (2)
- ◯ A few times (3)
- ◯ Many times (4)
- ◯ Every day (5)

SCMA_Q4 Do you know what this bird is?

- ◯ Yes (1)
- ◯ No (3)

*Skip To: SCMA_Q7 If Do you know what this bird is? = No*

SCMA_Q5 What do you call this bird? (You can enter N/A if you do not have a name for this bird)

________________________________________________________________

SCMA_Q6 On a scale from "strongly disagree" to "strongly agree" please tell me how much do you agree with each statement (Presented in random order)

| | **Strongly disagree** (1) | **Somewhat disagree** (2) | **Neither agree nor disagree** (3) | **Somewhat agree** (4) | **Strongly agree** (5) |
|---|---|---|---|---|---|
| This bird is like my neighbor and makes me feel at home | ○ | ○ | ○ | ○ | ○ |
| This bird helps make this place what it is | ○ | ○ | ○ | ○ | ○ |
| This bird is beautiful and I enjoy watching it | ○ | ○ | ○ | ○ | ○ |
| This bird has a beautiful song | ○ | ○ | ○ | ○ | ○ |
| I am excited to find this bird | ○ | ○ | ○ | ○ | ○ |
| This bird causes problems to the crops or the farms, for example by eating the crop | ○ | ○ | ○ | ○ | ○ |
| This bird helps the crops, for example by controlling insect pests or rodents | ○ | ○ | ○ | ○ | ○ |
| This bird causes problems to other species that are important for me | ○ | ○ | ○ | ○ | ○ |
| I like learning about or studying this bird, where it lives and what it does | ○ | ○ | ○ | ○ | ○ |
| I like teaching others about this bird and its habitat | ○ | ○ | ○ | ○ | ○ |
| I find this bird annoying because it's too noisy | ○ | ○ | ○ | ○ | ○ |
| I dislike this bird because their droppings make a mess or they build nests in inconvenient places | ○ | ○ | ○ | ○ | ○ |

SCMA_Q7 On a scale from "strongly disagree" to "strongly agree" please tell me how much do you agree with each statement

| | **Strongly disagree** (1) | **Somewhat disagree** (2) | **Neither agree nor disagree** (3) | **Somewhat agree** (4) | **Strongly agree** (5) |
|---|---|---|---|---|---|
| This bird should be protected for future generations | o | o | o | o | o |
| It would be sad if this bird would no longer exist | o | o | o | o | o |

SCMA_Q8 Do you have any memories with this bird? Do you have any comments about this bird?

________________________________________________________________

________________________________________________________________

________________________________________________________________

________________________________________________________________

________________________________________________________________

**Start of Block: Environmental attitudes**

A1 Have you gone birdwatching in the past two years?

- ◯ Yes (1)
- ◯ No (2)

A2 How many species of birds can you identify in Guanacaste?

- ◯ Less than 10 (1)
- ◯ 10-20 (2)
- ◯ 21-40 (3)
- ◯ 41-60 (4)
- ◯ more than 60 (5)

A3 How often do you watch TV shows, or read something about birds?

- ◯ Never (1)
- ◯ Rarely (2)
- ◯ Sometimes (3)
- ◯ Often (4)

---

**Start of Block: Demographics**

D1 In this last section of the survey, we would like to learn more about your background and your current household characteristics. You can be assured that all your answers will be kept confidential. This information will only be used to report results among groups of people. We will never identify individuals or households with these responses.

D2 Please type in your email address if you want an electronic version of research results (optional)

________________________________________________________________

D3 Age (in years)

________________________________________________________________

D4 Gender

- ◯ Female (5)
- ◯ Male (6)
- ◯ Other (7)
- ◯ Prefer not to answer (8)

D5 Where do you live? (City or town, Country)

________________________________________________________________

D6 Where do you consider home? (City or town, Country)

________________________________________________________________

D7 What is the highest level of education you have completed?

- ○ Primary school (last year graded) (1) ________________________________________________
- ○ High school or equivalent (2)
- ○ Vocational/Technical school (3)
- ○ Bachelor's degree (4)
- ○ Specialization (5)
- ○ Licenciature (6)
- ○ Master's degree (8)
- ○ Doctoral degree (9)
- ○ Other (10)

D8 Which of the following categories best describes your area of employment (regardless of your actual position)? Check all that apply

- ☐ Unemployed (2)
- ☐ Retired (3)
- ☐ Agriculture (4)
- ☐ Cattle production (5)
- ☐ Renting property (6)
- ☐ Renting machinery (7)
- ☐ Scientific or technical, research (including biologist) (8)
- ☐ Birdwatching (10)
- ☐ Education (11)
- ☐ Business, marketing, administration (13)
- ☐ Government and public administration (14)
- ☐ Health Care, social assistance (15)
- ☐ Transportation (16)
- ☐ Arts, entertainment, publicity (19)
- ☐ Construction (17)
- ☐ Other (18) ________________________________________________

D12 What do you consider your place of residence to be?

○ Large city or urban area (1)

○ Small city or town (3)

○ Rural area, on a farm or ranch (5)

D13 Please answer the next question only if you feel comfortable answering it. How would you rate your own financial situation today? Would you say it is excellent, good, only fair, or poor? In the following scale from 1 to 10, where 1=poor and 10=excellent, where would you place yourself?

| | Poor | | Only fair | | | Good | | Excellent | |
|---|---|---|---|---|---|---|---|---|---|
| | 1 | 2 | 3 | 4 | 5 | 7 | 8 | 9 | 10 |
| Your financial situation () | | | | | | | | | |

D14 **Thank you for participating in this survey!**

If you have any additional comments, feel free to type them here.
Please do not hesitate to contact us if you have any questions regarding our research project or how the results will be used.

Alejandra Echeverri Ochoa, Co-investigator and PhD Candidate, Institute for Resources, Environment, and Sustainability, University of British Columbia.

________________________________________________________

________________________________________________________
